# Supervised discovery of interpretable gene programs from single-cell data

**DOI:** 10.1101/2022.12.20.521311

**Authors:** Russell Z. Kunes, Thomas Walle, Tal Nawy, Dana Pe’er

**Author notes:** equal contribution (Russell Z. Kunes, Thomas Walle).

## Abstract

Factor analysis can drive biological discovery by decomposing single-cell gene expression data into a minimal set of gene programs that correspond to processes executed by cells in a sample. However, matrix factorization methods are prone to technical artifacts and poor factor interpretability. We have developed Spectra, an algorithm that identifies user-provided gene programs, modifies them to dataset context as needed, and detects novel programs that together best explain expression covariation. Spectra overcomes the dominance of cell-type signals by modeling cell-type-specific programs, and can characterize interpretable cell states along a continuum. We show that it outperforms existing approaches in challenging tumor immune contexts; Spectra finds factors that change under immune checkpoint therapy, disentangles the highly correlated features of CD8^+^ T-cell tumor reactivity and exhaustion, finds a novel program that explains continuous macrophage state changes under therapy, and identifies cell-type-specific immune metabolic programs.

## Intro

Deciphering the genetic mechanisms that underlie cellular function is a central goal in biology. We frequently wish to know, for instance, how different cell types respond to external stimuli and how this alters processes within the cell. Although clustering analysis can delineate cell types in single-cell RNA sequencing (scRNA-seq) data, it is difficult to retrieve coherent, interpretable gene programs representing these cellular processes, and to quantify them in response to perturbation.

Gene programs are sets of genes defined by common tasks, such as metabolic pathways and responses to inflammatory cues or growth signals. Gene set scoring (e.g. scanpy score_genes) is a simple and widely used approach^1^ to query which known gene programs are active in which cells, but it is often confounded by gene set overlap and technical factors. Single-cell sequencing is particularly well suited to identify gene programs, since programs tend to be regulated by mechanisms that are shared across specific cell subpopulations. Coregulation creates collinearity in gene expression levels, lending low-dimensional structure to high-dimensional cell-by-gene count matrices. Matrix factorization is a means of mining this structure to identify candidate gene programs^2,3^, and it has become a core tool in single-cell analysis; for example, factorization by principal component analysis (PCA) appears early in most scRNA-seq analysis pipelines.

In principle, the power of factorization lies in summarizing biological activity as a compact set of cellular building blocks—it can provide a minimal vector representing the degree to which a cell activates each gene program, rather than a noisy vector of all observed genes or a single label denoting cell-type. Yet matrix factorization is an unconstrained problem (there are many ways to decompose a matrix), and unsupervised approaches such as PCA and non-negative matrix factorization (NMF) produce factors that are often uninterpretable or driven by technical artifacts^3,4^. Some methods take a supervised approach, applying known gene sets as priors to make detected factors more interpretable^5,6^. However, the results may not be faithful to the data being analyzed, because gene sets are typically defined in biological contexts that differ from those under study. In addition, cell-type factors tend to prevail in factor analysis because differences between cells are dominated by cell type^4^. The popular practice of partitioning data by cell type and factoring each subset separately mitigates this issue, but makes it impossible to find shared programs.

A successful factor analysis method should identify all active gene programs in a dataset, including variations specific to biological context as well as novel factors, and it should quantify the degree to which each gene program is executed by each cell type. We have developed the factorization method Spectra (supervised pathway deconvolution of interpretable gene programs) to provide meaningful annotations of cell function by balancing prior knowledge with data-driven discovery (https://github.com/dpeerlab/spectra). Spectra incorporates existing gene sets and cell type labels, which are robustly determined by clustering, as biological priors. It explicitly models cell type and represents input gene sets as a gene-gene knowledge graph, using a penalty function to guide factorization towards the prior graph. The graph representation enables Spectra to generate factors that reflect the biological context under study, by modifying the input gene graph in a data-driven fashion, and the algorithm can also identify novel gene programs from residual unexplained variation. The degree of reliance on biological priors can be tuned with a global parameter.

Spectra’s ability to minimize the influence of cell type allows it to identify factors that are shared across cell types. We show that it outperforms existing approaches by solving longstanding challenges in tumor immune contexts; Spectra finds factors that change under immune checkpoint therapy (ICT), disentangles the highly correlated features of CD8^+^ T cell tumor reactivity and exhaustion under ICT, learns a novel gene program that explains the continuum of macrophage state changes under therapy, and identifies metabolic programs specific to different immune cell types.

## Results

### Spectra factor analysis identifies interpretable gene programs from single-cell data

To model gene expression, we assume that each cell executes a small number of gene programs and that the observed expression in a cell is determined by the sum of its active programs. Spectra decomposes the cell-by-gene expression matrix into a cell-by-factor matrix that identifies and quantifies the programs executed by each cell and a factor-by-gene matrix representing the genes in each gene program (**Fig. 1** and Methods). As input, the algorithm receives a normalized cell-by-gene count matrix, a cell-type annotation for each cell, and either a list of gene sets, or gene-gene relationships in the form of knowledge graphs. As output, Spectra provides a set of normalized global and cell-type-specific factor matrices, representing the gene loadings for each identified factor (gene scores); a sparse matrix of normalized factor loadings for each cell (cell scores); and the posterior gene knowledge graph representing factors inferred from the data (see Methods for a technical description of Spectra and parameter settings).

**Fig. 1.**
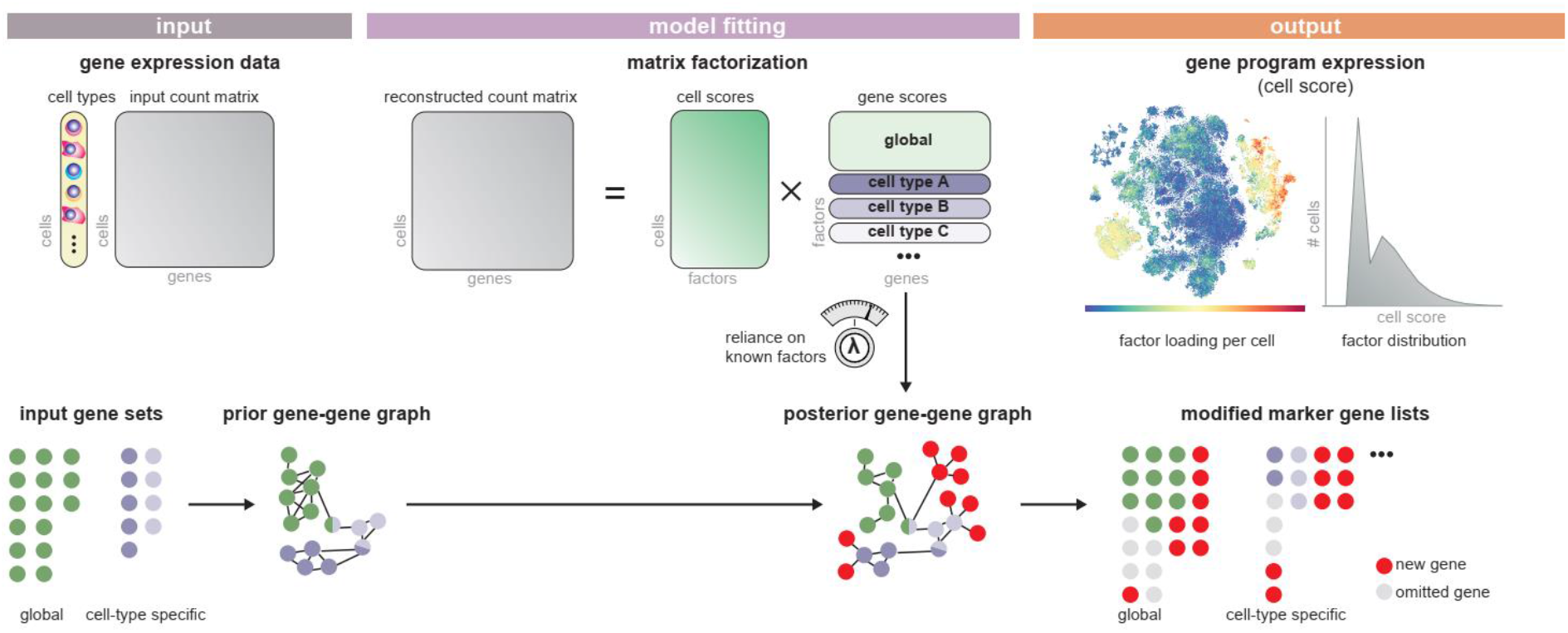
Spectra retrieves gene programs from scRNA-seq data using biological priors. As input, Spectra receives a gene expression count matrix with cell type labels for each cell, as well as pre-defined gene sets, which it converts to a gene-gene graph. The algorithm fits a factor analysis model using a loss function that optimizes reconstruction of the count matrix and guides factors to support the input gene-gene graph. As output, Spectra provides factor loadings (cell scores) and gene programs corresponding to cell types and cellular processes (factors).

Spectra attempts to balance prior knowledge and interpretability with faithfulness to the data. Two key features distinguish it from other factorization methods. First, Spectra uses known cell type information and allows for cell-type-specific factors; by incorporating cell-type-specific gene weights that explain away constitutively expressed cell-type marker genes, it mitigates their influence on the factors. Second, Spectra uses existing gene sets as a prior, which it represents as a gene-gene knowledge graph, enabling their data-driven modification as well as the derivation of entirely new factors. Together, these features allow Spectra to identify more interpretable factors and to discover new biology.

Given that clustering is usually superior to factor analysis for identifying cell types, we provide cell type labels as input to Spectra, which models the influence of a factor on gene expression relative to baseline expression per cell type. Modeling cell types also enables the incorporation of both global and cell-type-specific factors for improved inference. For example, while the T-cell receptor (TCR) activation program should be limited to T cells, many of its genes are also activated by different programs in other cell types, which confuses traditional factor analysis. Cell type information enables Spectra to decompose gene expression in a cell-type-specific manner rather than assuming that all cell types use identical gene programs.

Spectra’s likelihood function ensures that after decomposition, the reconstituted matrix closely matches the input matrix, and its penalty function guides gene factorization towards the gene-gene knowledge prior (Methods). To encourage factors to capture our prior, we use binary gene-gene relationships as evidence for gene participation across similar factors. Spectra takes input gene sets and turns each into a fully connected clique in the input graph, indicating that each gene pair in the set is related. Factors are thus scored by how well they match the data, as well as how many edges in the gene-gene graph support them.

Most established gene sets are derived from bulk sequencing data and multiple biological contexts. In contrast, analysis typically involves a dataset from a specific biological context that only utilizes subsets or variations of these gene programs. Spectra can take a broad compilation of input gene sets and determine which are supported by the data, ignoring those that are dissimilar to its identified factors. Encoding the prior as a graph facilitates computational efficiency, and more importantly, it allows Spectra to adapt gene programs by adding or removing edges in the prior graph to generate the posterior graph. The algorithm incorporates background edge and non-edge rates, provided as input parameters or learned from the data, to determine edge addition and removal rates. A critical feature of Spectra is that it can detach factors from graph penalization to learn entirely new factors. Effectively, Spectra attempts to explain as many of the input gene counts as possible by adapting the input gene graph (providing highly interpretable factors), and uses the residual unexplained counts to identify non-penalized factors that can capture entirely novel biology.

By design, Spectra is thus empowered to reduce the dominance of cell type while detecting more subtle global and cell-type-specific gene programs; it uniquely balances prior information for maximal interpretability with the data-driven modification of existing gene sets and discovery of new programs.

### Application of Spectra to an immuno-oncology dataset

We applied Spectra to scRNA-seq data from non-metastatic breast cancer patients before and after pembrolizumab (anti-PD-1) treatment (‘Bassez dataset’), representing a challenging and potentially impactful context for factorization (**Fig. 2a**)^7^. The original study used clustering and gene set analysis to identify therapy-induced changes, and employed TCR sequencing to define patients’ clonal T-cell expansion status under anti-PD-1 as a surrogate for ICT response^7^. We use this data to demonstrate Spectra’s features, evaluate its performance and showcase its ability to elucidate biological insight.

**Fig. 2.**
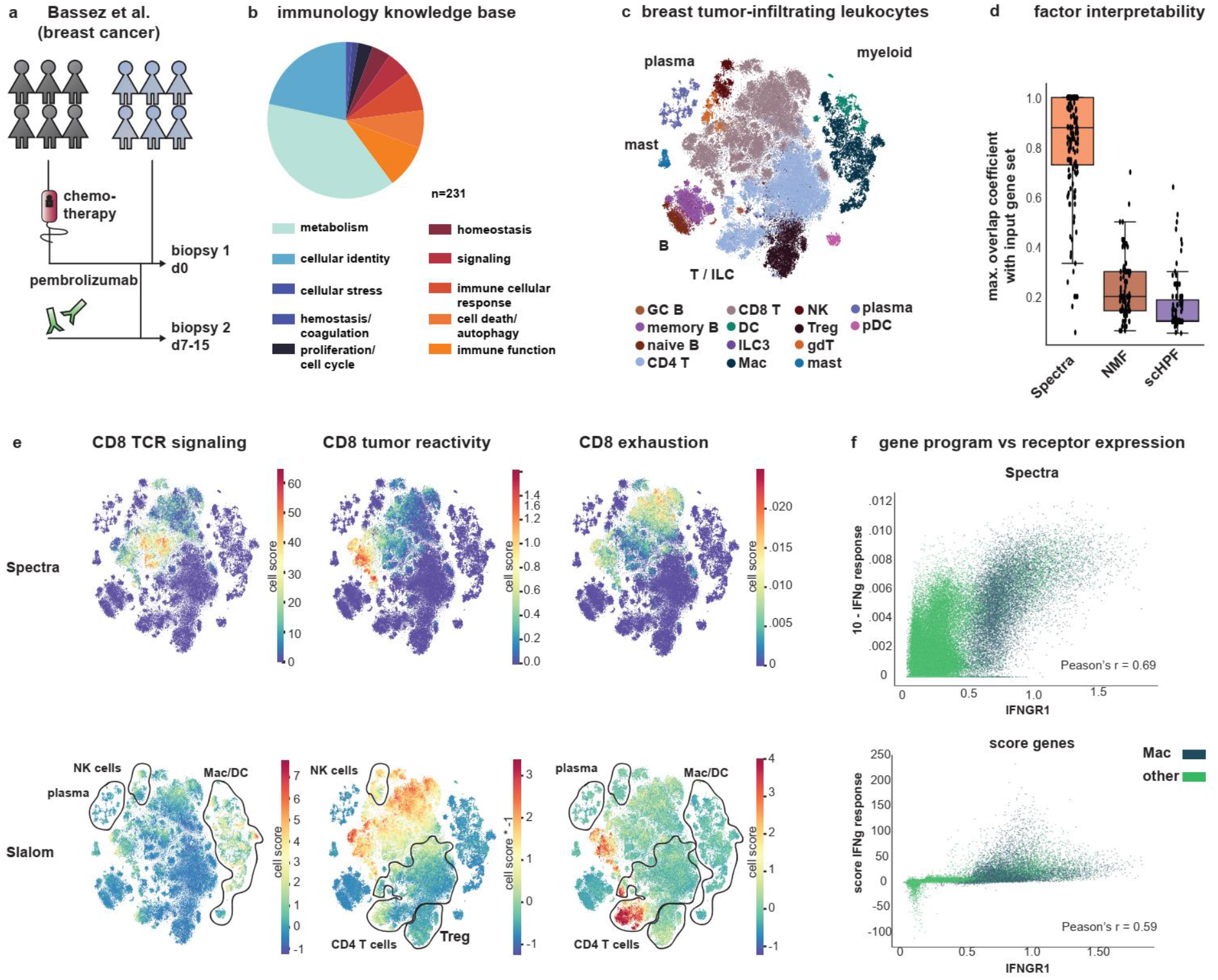
Spectra identifies interpretable factors that change under PD-1 perturbation. **a**, Treatment and scRNA-seq sampling regime of breast cancer patients in Bassez et al., 2021^7^ (‘Bassez dataset’). We used Spectra to analyze tumor infiltrating leukocytes from this data. **b**, Gene set categories in the immunology knowledge base (n=231 gene sets). **c**, t-SNE embedding of tumor-infiltrating leukocytes (n=97,863 cells), colored by cell type. **d**, Maximum overlap coefficient of every factor generated by Spectra (n=152), NMF (n=100) or scHPF (n=100) with every input gene set. Boxes and line represent interquartile range (IQR) and median, respectively; whiskers represent ±1.5 x IQR. **e**, t-SNE projection colored by cell scores for the Spectra or Slalom factors best representing CD8^+^ T-cell tumor reactivity and CD8^+^ TCR signaling (n=97,863 cells). Black contours highlight aberrant expression patterns. **f**, Cell scores for Spectra and scanpy.score_genes factors plotted against MAGIC-imputed (t = 3) *IFNGR1* expression for each cell, colored by macrophage (Mac) or non-macrophage (other) cell type (n=97,863 cells). d: day, T: T cell, gdT: gamma-delta T cell, pDC: plasmacytoid dendritic cell, Mac: macrophage, NK: Natural Killer cell, B: B cell, Mac: macrophage, mast: mast cell, mono: monocyte, TCR: T cell receptor, Treg: regulatory T cell, DC: dendritic cell, GC: germinal center, plasma: plasma cell.

We first curated a general resource of immunology gene sets that can be input to Spectra for analyzing any immune-related dataset. Our knowledge base contains 231 relevant gene sets, including 50 ‘cellular identity’ gene sets to define input cell types and 181 ‘cellular process’ gene sets (**Fig. 2b**, **Supplementary Table 1** and Methods). To generate the resource, we developed 97 new gene sets, including 14 from perturbation experiments, and added these to 134 gene sets from publications and external databases, some of which we modified. Of the cellular processes, 150 apply to most cell types in the data (e.g. leukocytes) and are designated as global, and 31 apply to individual cell types. We designed the cellular process gene sets to have comparable size (median n = 20 genes per gene set) and relatively little overlap (median pairwise overlap coefficient 40%) to enable dissection of a large number of processes and to avoid size-driven effects.

To ensure consistent cell typing at a resolution best suited for Spectra, we independently annotated the dataset into 14 broad cell types of established discrete lineages (e.g. CD8^+^ T cells, macrophages, dendritic cells, plasma cells), leaving Spectra to infer factors associated with finer cell type distinctions such as T-cell activation or macrophage polarization (**Fig. 2c**, **Supplementary Fig. 1**, **Supplementary Table 2** and Methods). We provided the broad cell type labels and our immunology knowledge base as input, and fit the Spectra model to the Bassez dataset using default parameters (Methods), resulting in 152 global and 45 cell-type-specific factors. Most cell-type-specific factors refer to CD4^+^ T cells (n = 12), CD8^+^ T cells (n = 7) and myeloid cells (n = 6).

We first assessed whether Spectra is able to identify biologically interpretable gene programs. For every factor, Spectra estimates a dependence parameter (η) between zero and one that quantifies reliance on the input gene-gene graph. Most factors (171, or 86.8%) are strongly constrained by the gene-gene graph (η ≥ 0.25), whereas 26 (13.2%) are novel (**Supplementary Fig. 2**). We found that factors with η ≥ 0.25 overlap substantially (≥ 0.5 of genes) with an input gene set, enabling their interpretation. In contrast, widely used unbiased factorization approaches, NMF and scHPF^3,8^, typically produce factors that do not agree well with annotated gene sets (**Fig. 2d**), underscoring the difficulty of interpreting gene programs derived by these approaches.

We next assessed whether Spectra can provide more biologically sensible factor loadings (assignments of gene programs to cells) than other supervised methods when using identical input gene sets. Spectra uses cell-type labels and cell-type-specific input gene sets to restrict factors to their appropriate cell type; for example, it limits CD8-specific TCR signaling, tumor reactivity and exhaustion factors to CD8^+^ T cells (**Fig. 2e**). In contrast, Slalom^5^, a factorization method based on gene set priors, misassigns some TCR activity, CD8^+^ T cell exhaustion, and tumor reactivity to the myeloid, NK cell and plasma cell lineages (**Fig. 2e**), likely because many genes in these factors participate in multiple programs. For example, *MAFF* is induced by TCR signaling^9^ but also participates in monocyte inflammatory responses^10^.

Pleiotropy similarly confounds the most widely used single-cell gene-set annotation tool, score_genes^11^. For example, Spectra’s interferon gamma (IFNg) response factor is well correlated with the IFNg receptor upstream of this gene program and correctly captures it across all cell types, whereas score_genes IFNg response is detected almost exclusively in the myeloid population (**Fig. 2f**). This myeloid bias is due to differences in baseline expression across cell types, especially HLA-II genes, which are preferentially expressed by myeloid antigen-presenting cells (**Supplementary Fig. 3**). Spectra overcomes pleiotropy by penalizing the attribution of one gene to multiple factors—it attributes the gene to the factor best supported by total expression in a given cell. Spectra is able to identify IFNg activity and its previously reported activation by ICT^12,13^ across expected immune cell types^14^ because it learns these factors in a cell-type-specific manner, accounting for differences in baseline expression across cell types.

Thus, in addition to yielding more interpretable gene programs than other supervised methods, Spectra is better at inferring which cells these programs are active in, enabling it to detect subtle effects of ICT on multiple cell types that are missed by score_genes.

### Spectra outperforms other methods in delineating and assigning gene programs

We systematically benchmarked Spectra against other widely used methods by measuring how well they identify coherent gene programs and assign activity to cells. A key feature of Spectra is that it can modify input gene sets in a data-driven manner. We first evaluated the quality of output gene programs by holding out 30% of genes from 20 input gene sets and tracking whether these genes are identified in the resulting factors (Methods). Spectra factors recover substantially more genes than Slalom (**Fig. 3a**); for example, among the 50 genes with highest gene scores (top 50 marker genes) for the MYC factor, Spectra identifies 7 of 33 held-out genes (*GLN3*, *NOP16*, *PAICS*, *APEX1*, *PA2G4*, *TSR1* and *TRAP1*) relevant to MYC signaling, while MYC signaling was not retained by Slalom (Methods), and likewise performs well for other cellular processes (**Fig. 3b** and **Supplementary Fig. 4**). Among the genes with highest gene scores, Spectra also recovers known MYC targets *DKC1* and *TOMM40*, which are absent from the training and hold-out sets.

**Fig. 3.**
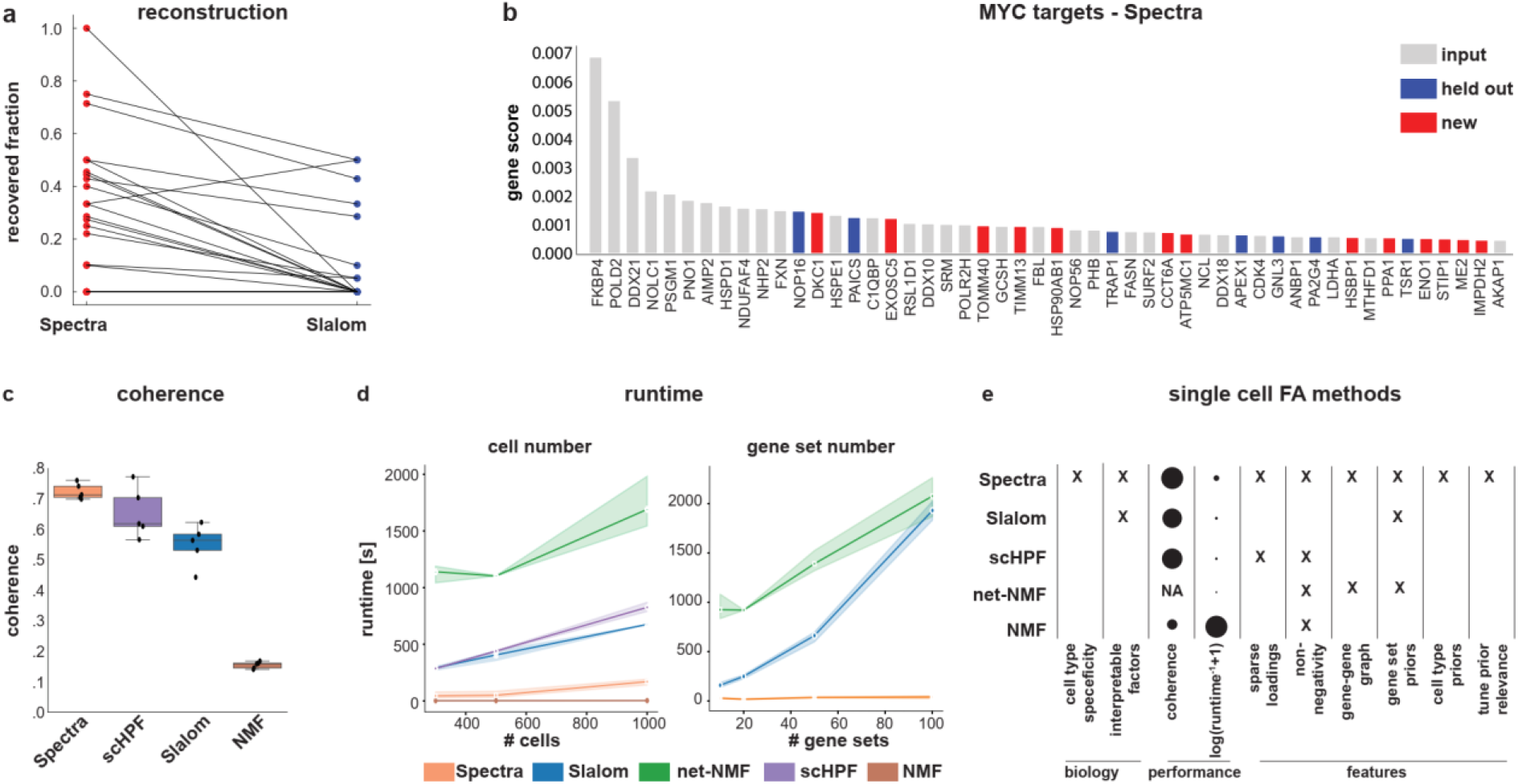
Spectra outperforms other factor analysis methods. **a**, Proportion of held-out genes recovered by Spectra or Slalom from the Bassez dataset^7^, for each input gene set tested. Lines connect identical input gene sets. **b**, Gene scores for the top 50 marker genes of the MYC target Spectra factor, highlighting new genes (red) and held-out genes (blue) that were recovered. **c**, Coherence (mean pairwise log-normalized co-occurrence rate among top 50 markers) of factors generated by various factor analysis methods, using a random sample of 10,000 cells from the Bassez dataset, with 14 cell types and 181 input gene sets. The experiment was repeated n = 5 times. Boxes and line represent interquartile range (IQR) and median, respectively; whiskers represent ±1.5 x IQR. **d**, Runtime dependence on cell number (left panel) and gene set number (right panel). The experiment was repeated n = 3 times; shading indicates 95% confidence interval. **e**, Properties and performance metrics of factor analysis methods. Dot diameter is proportional to coherence for the experiment in **c**, or inversely proportional to runtime in seconds for the experiment described in **d** (left panel) using 1000 cells.

To provide a more systematic evaluation of new genes, we reasoned that genes belonging to a shared program should exhibit coherence by appearing together in the same cells. Spectra explicitly uses this signal to add new genes; to ensure that the data is not overfit, we applied factor analysis with held-out cells (not used in training), and evaluated the coherence of inferred factors in the test set (Methods). Spectra and other methods that take the sparsity of scRNA-seq data into account (Slalom, scHPF) perform well, while generic models (NMF) do not (**Fig. 3c**). The key advantage of supervised approaches is that by seeding inference with a known gene set, coherent genes are more likely to be biologically meaningful.

Matrix factorization methods rely on objective functions that implicitly encourage the estimation of diverse gene programs, such that programs expressed in similar contexts (e.g. T-cell activation and exhaustion) are often recovered as a single merged factor. To test the ability to robustly identify correlated programs, we simulated gene expression data from a generic factor analysis model with both correlated and uncorrelated factors (Methods). Spectra’s use of prior knowledge allowed it to separate highly correlated factors, unlike other methods (**Supplementary Fig. 5a**). We next asked whether Spectra can accurately quantify the activity of inferred factors across cells, which is particularly challenging because pleiotropy creates correlation between gene programs. Given the lack of ground truth, we synthesized data with features similar to real data and benchmarked the assignment of factor loadings to cells (Methods). As gene set overlap increases, score_genes surges in false positive score estimates, while Spectra is able to correctly assign expressed factors to cells (**Supplementary Fig. 5b**). Due to their multivariate nature and sparsity, factorization methods select the factors that best explain the data globally, such that each factor accounts for expression that is not already explained by other factors. Factor analysis is thus superior to score_genes even for the simple task of scoring gene sets.

In contrast to Spectra, Slalom suffers a substantial drop in accuracy as the number of active gene sets increases (**Supplementary Fig. 5c**). Moreover, Slalom can only assess a few dozen gene-sets before its runtime becomes prohibitive, whereas Spectra’s graph-based representation allows it scale to hundreds of thousands of cells and hundreds of gene programs, with a runtime that outperforms both scHPF and Slalom (**Fig. 3d**). Our benchmarking demonstrates that Spectra is faster and infers programs with superior interpretability and coherence, while retrieving more ground truth factors (**Fig. 3e**).

### Spectra disentangles correlated features of CD8^+^ T cell tumor reactivity and exhaustion

To understand how tissues respond to cancer treatments and to ultimately improve therapeutic efficacy, we seek to quantify gene program changes under therapy on a per-cell-type basis. One population that is particularly important to track under ICT consists of non-dysfunctional tumor-reactive CD8^+^ T cells, a subset of T cells that recognize tumor-associated antigens^15^ and are also cytotoxic^16,17^. These cells express clonal TCRs and specific markers (e.g. *ENTPD1, CXCL13*)^18,19^, and can accumulate upon PD-1/PD-L1 checkpoint blockade in a process called clonal expansion^20,21^. Conversely, T cells that expand clonally under ICT are likely to be tumor-reactive^17,20^. These T cells may also gradually become exhausted (lose effector capacity) upon prolonged antigen exposure in the tumor microenvironment^22,23^. Although exhaustion and tumor reactivity lead to different cellular behaviors with highly consequential phenotypes, their gene programs are correlated and challenging to discriminate computationally; clustering approaches, for example, typically generate groupings that combine exhaustion, tumor reactivity and cytotoxicity features^7,24^.

We evaluated Spectra’s ability to deconvolve tumor reactivity and exhaustion programs and to quantify therapy-induced changes in each, focusing on CD8^+^ T cells in the Bassez dataset (**Fig. 4a**). Genes from these two programs are correlated in this data (**Supplementary Fig. 6a**), likely explaining why they were not distinguished in prior work^7,24^. Consequently, their cell scores were found to be identically distributed in responders and non-responders (**Fig. 4b**). Yet the absence of tumor-reactive, non-terminally exhausted states in responders is inconsistent with the treatment-induced clonal expansion of these states observed in mouse models^25,26^ and longitudinal phenotyping of cancer patients^20,21^, and it conflicts with the proven efficacy of ICT in this clinical setting^27^.

**Fig. 4.**
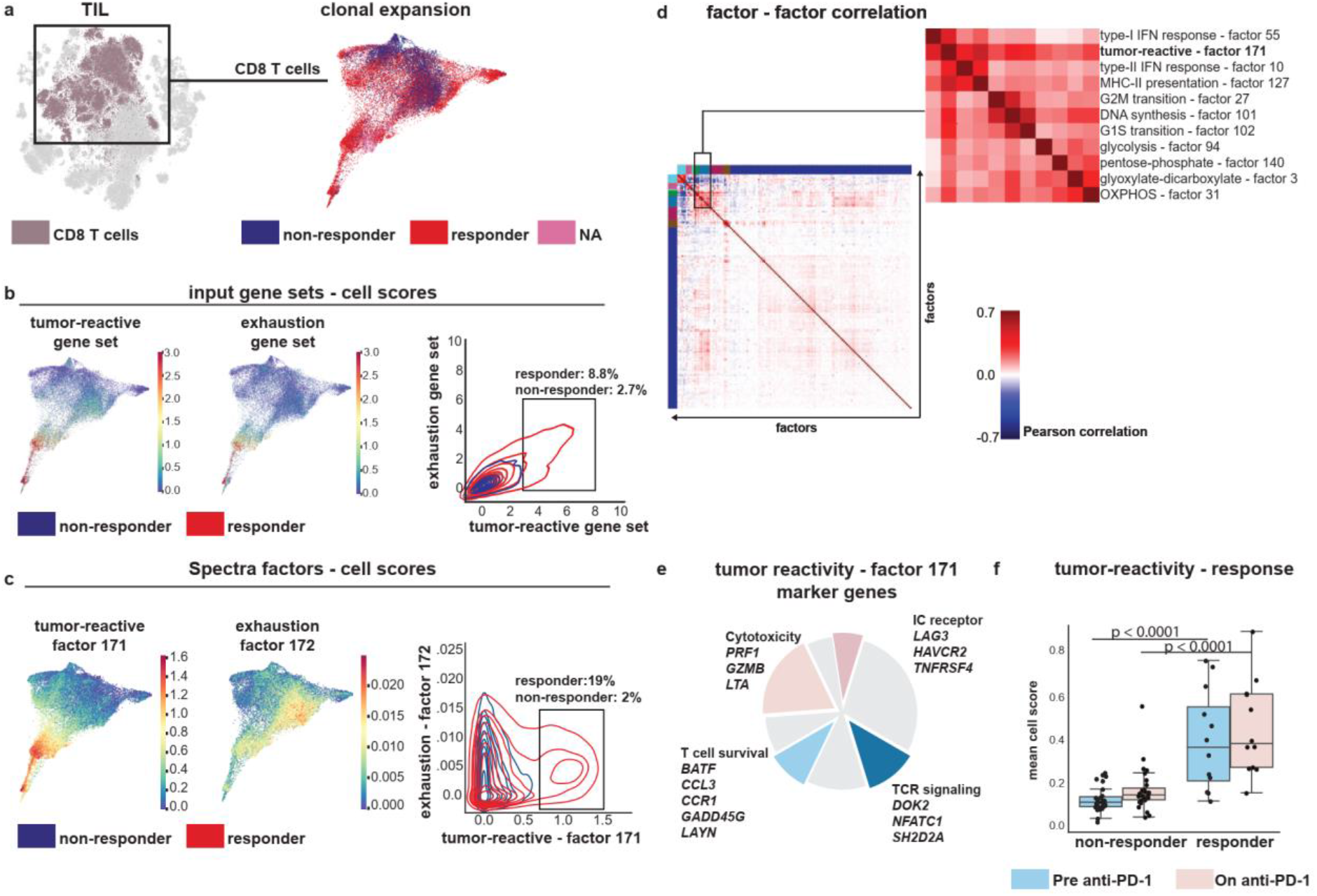
Spectra deconvolves the highly correlated features of tumor reactivity and exhaustion in CD8^+^ T cells. Analysis of breast cancer infiltrating leukocytes from the Bassez scRNA-seq dataset (n = 42 patients)^7^. **a**, t-SNE map of the entire dataset highlighting CD8^+^ T cells (left), and force-directed layout (FDL, n=43,066 cells) of CD8^+^ T cells labeled by evidence of clonal T-cell expansion in the donor (right). T cells that expand clonally under ICT are likely to be tumor-reactive^17,20^. **b**,**c**, FDL of CD8^+^ T cells (n=43,066) colored by tumor reactivity (left) or exhaustion (right) cell scores, and contour plots depicting cell score density distribution from patients with (red) or without (blue) clonal expansion of T cells under therapy. Cell scores were obtained using scanpy.score_genes (**b**) or Spectra (**c**). Only Spectra disentangles processes with correlated gene expression. **d**, Pearson correlation coefficients of factor cell scores (n=43,066 cells). Tumor reactivity is highly correlated with factors related to CD8^+^ T cell effector function and ICT response. **e**, New genes identified by Spectra (n = 38) among the genes with highest gene scores (top 50 marker genes) in the tumor reactivity factor, highlighting processes known to be involved in tumor reactivity. See **Supplementary Table 3** for full list of genes. **f**, Per-sample mean cell scores for the Spectra tumor reactivity factor in positive cells (loading > 0.01). Boxes and line represent interquartile range (IQR) and median, respectively; whiskers represent ±1.5 x IQR. P values (two-sided) were calculated using Mann-Whitney-U tests; pre-anti-PD-1 (n = 40): 3.835 x 10^−5^ statistic: 308 Cohen’s d: 1.510; on anti-PD-1 (n = 40): p = 2.001 x 10^−5^, statistic = 313 Cohen’s d: 1.491.

Whereas gene set scores are markedly correlated and fail to distinguish expanding from non-expanding clones (**Fig. 4b**), Spectra clearly disentangles them, identifying a substantial tumor-reactive population, with varying degrees of exhaustion, that is almost exclusive to responders (**Fig. 4c**). In support of these observations, incompletely exhausted ‘pre-dysfunctional’ T cells are known to be critical for ICT efficacy^15^. Importantly, Spectra extracts gene programs directly from the unlabeled data and does not need response status to successfully dissect these features. Spectra’s likelihood function discourages overlap between gene programs when a single program is sufficient to explain the observed count matrix, harnessing unique features of each gene-set to associate cells with the best fit program. We calculated the covariance of individual genes with their respective gene set scores, and identified *CXCL13* as having highest covariance with both tumor reactivity and exhaustion, indicating that it is a driver of the overlap in these scores (**Supplementary Fig. 6b,c**). Spectra assigns a high weight for *CXCL13* in the tumor reactivity but not in the exhaustion factor, and strongly weights genes related to TCR signaling, T cell activation and cytotoxicity in the tumor reactivity factor. We find that only 4 genes overlap among the 50 genes with highest gene scores for each factor, whereas the exhaustion factor mostly includes exhaustion-inducing transcription factors (*TOX, NR4A1*) and *PDCD1* (PD-1).

After establishing that the tumor reactivity factor describes biologically relevant features, we explored its correlation with other factors in CD8^+^ T cells. Reactivity correlates with proliferative programs, as expected for cells that expand under ICT, as well as oxidative phosphorylation and glycolysis, processes associated with enhanced CD8^+^ T cell effector function^28^, and IFN-γ signaling, a key mediator of ICT efficacy^12^ (**Fig. 4d**). Of the 50 genes in tumor-reactive CD8^+^ T cells with highest gene scores, 43 are outside the input gene set, but recent studies support their roles in tumor reactivity (**Fig. 4e** and **Supplementary Table 3**). Four of the new genes are among 21 found to be differentially expressed in functionally validated tumor reactive CD8^+^ T cells in lung cancer patients^29^. This includes the TCR signaling target *BATF*, which was recently shown to enhance effector function and counteract exhaustion^30,31^. Moreover, CCL3 and its cognate receptor CCR1 are upregulated during CD8^+^ T cell priming^32,33^ and lure CD8^+^ T cells into tumors^34^.

Separating tumor reactivity also allowed us to assess its independent contribution to clonal expansion and therapeutic response. We found that expression of this factor is higher in responders (patients with clonal T cell expansion) at baseline than non-responders, and in responders it increases further under therapy (**Fig. 4f**), consistent with the reported association between tumor-reactive cell clusters and therapeutic response^35,36^. Spectra thus cleanly disentangles a CD8^+^ T cell tumor reactivity program that is associated with response to ICT at the cell and patient levels, enabling us to further identify correlated processes and novel mediators of reactivity which may be nominated as candidate targets for enhancing ICT efficacy.

On the same dataset, Slalom only found a factor strongly enriched for exhaustion genes, which overlaps with the highest scoring factor for tumor reactivity by 35 genes (**Supplementary Fig. 6d,e**). scHPF factors show a low enrichment for both reactivity and exhaustion gene sets (**Supplementary Fig. 6d**). Unlike Spectra, neither Slalom nor scHPF were able to distinguish a clonally expanding tumor-reactive T cell population that is specific to responders (**Supplementary Fig. 6e,f**). Moreover, Slalom and scHPF factors failed to associate with patient-level response, defined as a significant difference between expression in responders and non-responders before or under ICT (**Supplementary Fig. 6g**). Spectra can disentangle tumor reactivity from exhaustion because of its capacity to separate highly correlated gene programs (**Supplementary Fig. 5a**).

### Uncovering metabolic pathway utilization patterns across tumor infiltrating leukocytes

Metabolic processes are fundamental to cancer progression and therapeutic response, in part because cancer and immune cells compete for scarce nutrients such as essential amino acids^37,38^. Analyzing cellular metabolism has proven very challenging, as participating genes are involved in multiple pathways^38^. We asked whether Spectra’s ability to deconvolve overlapping gene programs (**Supplementary Fig. 6b,c**) can empower the inference of metabolic processes from tumor immune cells in the Bassez breast cancer dataset^7^. Spectra identified gene programs related to all 89 metabolic input gene sets (overlap coefficient > 0.25) and determined their expression across cell types, recapitulating known macrophage metabolic characteristics such as iron uptake, iron storage^39,40^ and cholesterol synthesis by cytochrome P450 enzymes^41,42^, as well as DNA synthesis in cycling germinal center B cells (**Fig. 5a**).

**Fig. 5.**
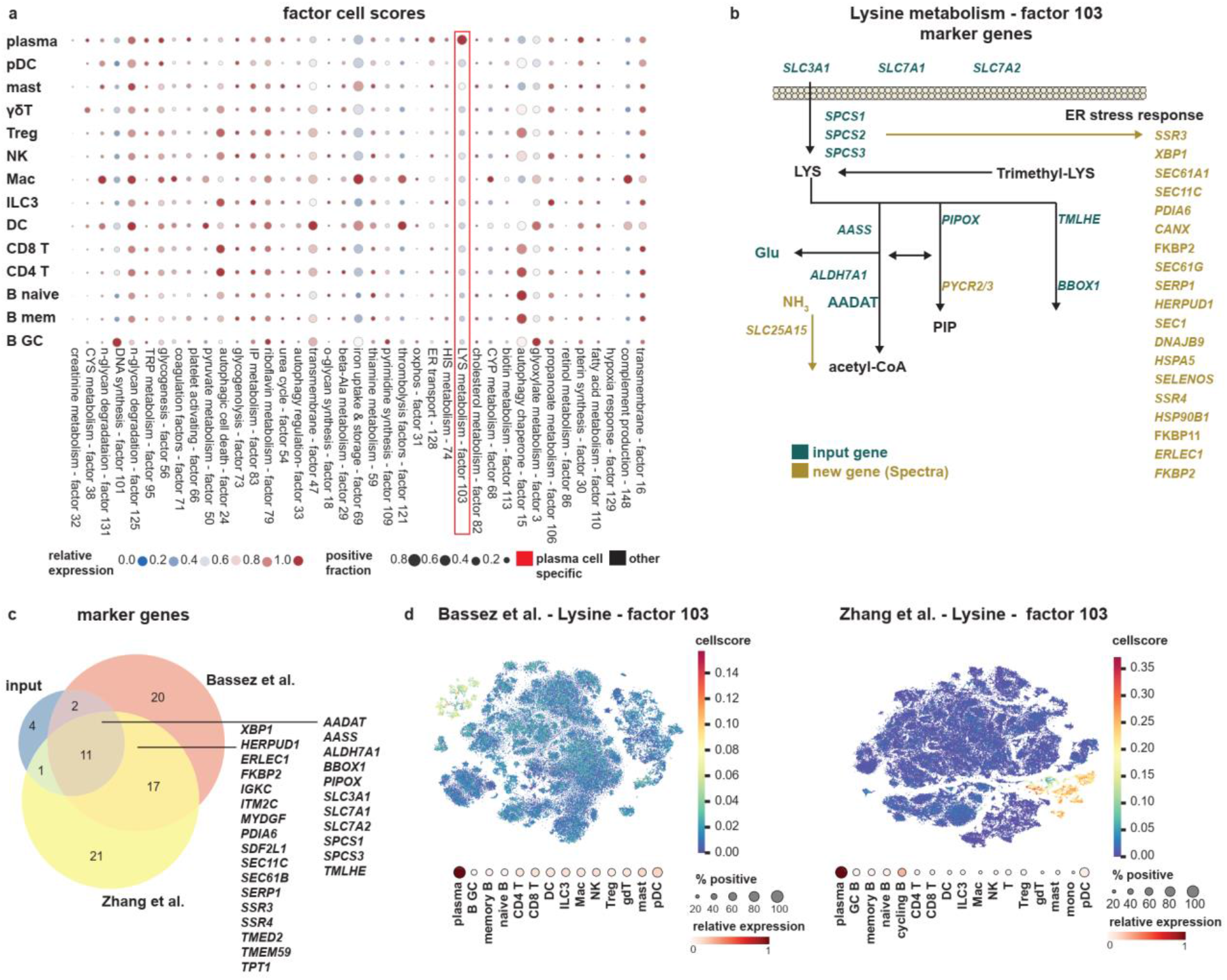
Spectra reveals cell-type-specific metabolic profiles in breast cancer data. **a**, Mean cell score among positive cells (score > 0.01) per cell type (y-axis) for each Spectra metabolic factor (x-axis) identified in the Bassez dataset^7^ (n=97,863 leukocytes). **b**, Input gene sets and new genes inferred by Spectra in the lysine metabolism pathway. **c**, Overlap of genes from the input lysine metabolism gene set with the top 50 marker genes from lysine metabolism factors identified in the Bassez^7^ and Zhang^24^ datasets. **d**, t-SNE embeddings of tumor infiltrating leukocytes, colored by Spectra factor cell scores in the Bassez (n=97,863 leukocytes) and Zhang datasets (n=150,985 leukocytes). ILC3: innate lymphoid cell type 3, T: T cell, gdT: gamma-delta T cell, pDC: plasmacytoid dendritic cell, Mac: macrophage, mono: monocyte, NK: Natural Killer cell, B: B cell, mast: mast cell, Treg: regulatory T cell, DC: dendritic cell, GC B: germinal center B cell, plasma: plasma cell.

Spectra also uncovered novel cell-type-specific expression of amino acid metabolic factors, such as cysteine metabolism in gamma delta T cells and lysine metabolism in plasma cells (**Fig. 5a**). Lysine is an essential amino acid found at lower concentrations in malignant lesions than adjacent normal breast tissue, likely due to the high metabolic demands of the tumor^43^. Among the top 50 marker genes of the lysine factor, Spectra retained 72% of the input gene set, including all key metabolic enzymes; it removed redundant amino acid transporters (*SLC25A21*, *SLC38A4*, *SLC6A14*, *SLC7A3*); and it retrieved degradation genes (*PYCR2, PYCR3, SLC25A15*) not found in the input set (**Fig. 5b**). Moreover, Spectra added genes involved in the unfolded protein response, including the two pivotal initiators *XBP1* and *ATF6* and their downstream targets (*ERLEC1*^44^, *SDF2L1*^45^, *HERPUD1*^46^, *PDIA6* ^47^). These genes are expressed more coherently and at higher levels in plasma cells than other cells, suggesting coordinated expression as a gene program (**Supplementary Fig. 7a**). ER stress regulates the capacity of plasma cells to produce immunoglobulins^48^, likely because large quantities of misfolded antibodies^49^ must be degraded, generating significant lysine^50^.

Slalom identified a factor in plasma cells with poorer resemblance to lysine metabolism (overlap coefficient 0.17 vs 0.72) and less biological coherence (**Supplementary Fig. 7b**), possibly because Slalom, unlike Spectra, does not use cell-type-specific gene scalings. The Slalom lysine factor misses ER stress genes and is contaminated with cell type markers (e.g. *SDC1*, *MZB1*), likely for the same reason (**Supplementary Fig. 7c**). scHPF lacks gene set priors and failed to identify a factor enriched for lysine metabolism genes; moreover, the scHPF factor with the highest number of lysine genes (2 of 18) is not expressed in plasma cells (**Supplementary Fig. 7d**).

To gauge Spectra’s stability and the reproducibility of its lysine factor, we fitted an independent Spectra model onto data from patients with metastatic breast cancer biopsied before and during paclitaxel chemotherapy with or without anti-PD-L1 (‘Zhang dataset’)^24^(**Supplementary Fig. 1b**), using identical parameters. Of the top 50 marker genes identified in the Bassez dataset, 28 were also identified independently in the Zhang dataset (**Fig. 5c**). This includes 17 of the 37 new genes learned independently and directly from the data in both datasets, and encompassed ER stress genes. The lysine metabolism factors from both datasets were specifically expressed in plasma cells, supporting the reproducible identification of both marker genes and cell scores by Spectra (**Fig. 5d**). Our results link lysine metabolism and ER stress as features of tumor infiltrating plasma cells in breast cancer.

### A novel gene program describes continuous macrophage state changes under therapy

Macrophages critically shape anti-tumor immunity and mediate resistance to immune checkpoint therapy by adopting immunosuppressive phenotypes under therapy (adaptive resistance); however, the effect of ICT on macrophage gene programs and their association with response remains unclear^51,52^. Bassez et al.^7^ linked a macrophage cluster expressing the complement gene C3 to therapy resistance (**Supplementary Fig. 8a,b**). While marker genes can help to identify cell populations that change under ICT, they do not necessarily represent biological processes. For example, complement genes such as CFB (which activates C3 (ref. ^53^)) exhibit opposite trends to C3 and are more highly expressed in responders (**Supplementary Fig. 8b,c**).

We evaluated whether Spectra can identify more interpretable gene programs underlying macrophage cell states and adaptive resistance mechanisms, using diffusion components to effectively visualize continuous states^54^. Specifically, we found that diffusion components 2 and 4 (DC2 and DC4) are most relevant for capturing maturation from monocyte-like to macrophage states, and for separating responders from non-responders, respectively (**Fig. 6a**). Cell scores for Spectra factors form gradients along DC2 with successive peaks from monocyte to macrophage states, beginning with CYP enzyme activity and TNF-α signaling (required for monocyte survival^55^), followed by glycolytic activity (likely required for monocyte activation^56^), a novel factor containing invasive and angiogenic mediators (‘invasion program’), and finally complement production, a key feature of mature macrophages^57^. Along DC4, Spectra identified programs for high type-II IFN signaling and MHC-II antigen presentation at one extreme, followed by IL-4/IL-13 response, and hypoxia signaling and the invasion program at the other (**Fig. 6a**; see **Supplementary Table 4** for factors associated with each DC). Spectra can thus characterize continuous macrophage states in tumor tissue, including hypoxia and related invasion programs, which may represent critical states along the macrophage spectrum.

**Fig. 6.**
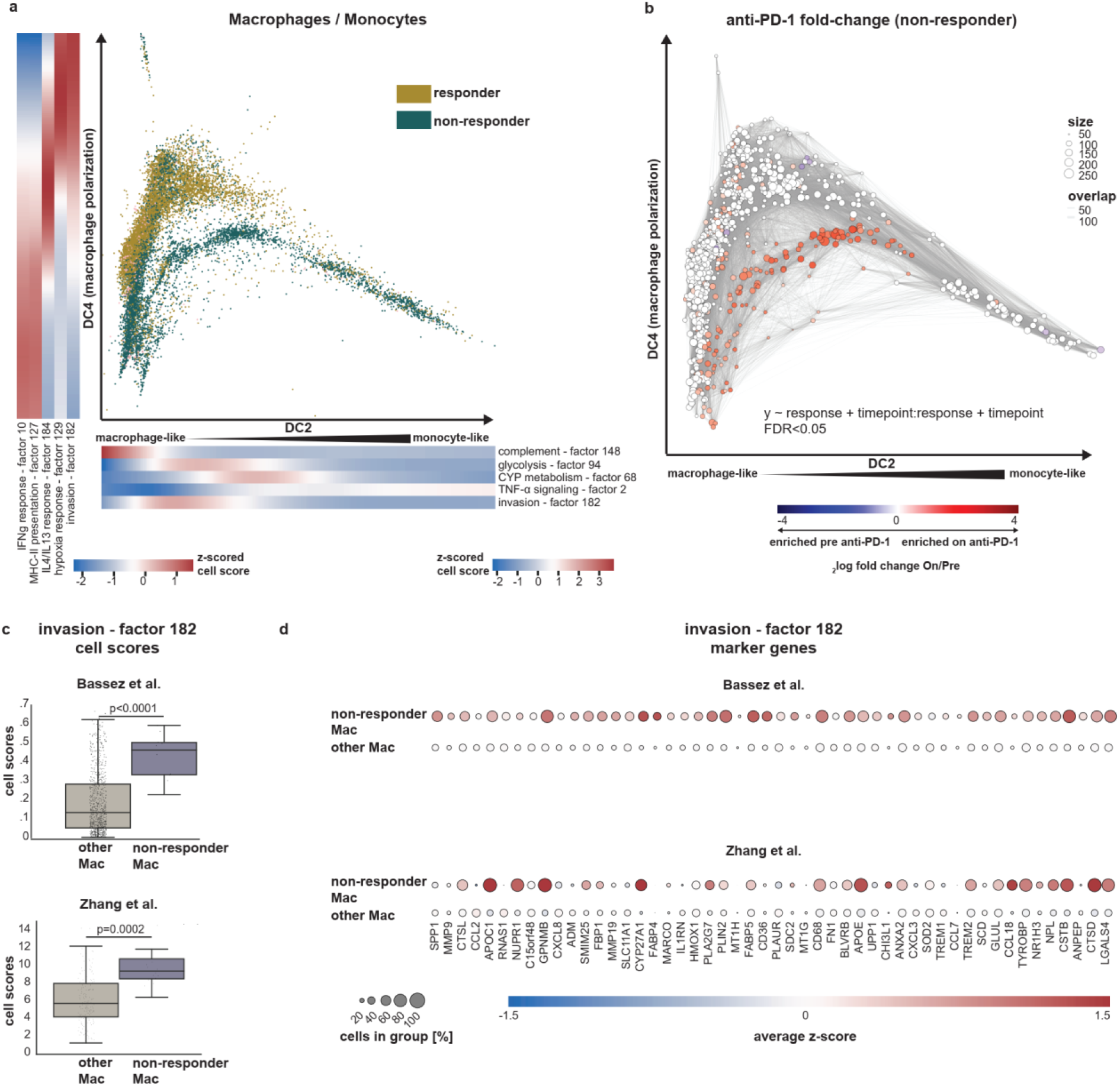
Spectra reveals therapy-induced macrophage gene expression programs. **a**, Macrophage (Mac) cells plotted along diffusion components DC2 and DC4, colored by patient-level T cell expansion status (responder, non-responder) in the Bassez et al dataset^7^ (n=12,132 cells). Heatmaps along the x and y axes indicate z-scored gene program cell scores, smoothened by fitting a generalized additive model (Methods). **b**, Graph with nodes representing cellular neighborhoods (n=858) plotted along DC2 and DC4, and edges representing overlap, colored by 2log(fold change) under anti-PD-1 in non-responders as estimated with Milo (Methods). The 2log(fold change) of non-significant (FDR ≥ 0.05) neighborhoods is set to 0. **c**, Average cell scores of macrophage neighborhoods (n=858) enriched in non-responders under therapy, and cell scores for all other macrophage neighborhoods in the independent Bassez and Zhang breast cancer datasets. Cell scores were calculated using the Spectra invasion factor (factor 182 from the Bassez dataset) or by using scanpy.score_genes on the top 50 marker genes of factor 182 in the Zhang dataset. P values (two-sided) were calculated using Mann-Whitney U tests. Bassez et al.: P value: 4.958×10^−5^ statistic: 1060 Cohen’s d: 1.494; Zhang et al.: P value: 3.739×10^−12^ statistic: 600886 Cohen’s d: 1.030.Boxes and line represent interquartile range (IQR) and median, respectively; whiskers represent ±1.5 x IQR. **d**, Mean expression, z-scored across cells (n=12,132 cells) in color code below and dot size indicating the percentage of cells with at least one detected copy of the indicated genes with legend below, of factor genes in non-responder macrophage populations and other macrophage populations in the Bassez (n=12,132 cells) and Zhang datasets (n=3,206 cells).

To find macrophage states that only change in non-responders under ICT—and could therefore confer adaptive resistance in patients—we used the Milo algorithm^58^, which avoids discretizing the macrophage phenotypic continuum. Milo revealed overlapping cellular neighborhoods (states) that only expand under anti-PD-1 therapy in non-responders (**Fig. 6b**) and are high in the novel invasion program (**Fig. 6c**). This invasion program exhibits low graph dependence (η = 0.24) (**Supplementary Fig. 8d**), indicating that it does not correspond to any input gene set; moreover, Slalom and scHPF do not identify a similar program (**Supplementary Fig. 8e**). Individual invasion program genes are coherently expressed in macrophages, only increase in non-responders, and include known invasion and metastasis mediators (*CTSL*^5*9*^, *CTSD*^60^, *CTSB*^61^, *CHI3L1*^62^, *SPP1*^63^, *PLIN2*^64^). Moreover, the invasion program includes cholesterol metabolism genes (*APOE*^65,66^, *APOC1*^67^, *CYP27A1*^68^, *GPNMB*^69^), some which have been linked to the suppression of inflammatory cytokine release (IL-6, TNF-alpha). Our results suggest that patients who do not respond to ICT may express these genes coordinately, constituting a new macrophage program (**Fig. 6d**). By focusing on residual expression that is not well explained by the gene knowledge graph, Spectra can thus find a novel gene program that is both interpretable and related to ICT response, unlike fully unsupervised approaches such as scHPF.

To test for replication on independent data, we scored expression of the top 50 marker genes of the Spectra invasion factor in the Zhang dataset^24^. We ran Milo and identified macrophage populations that expand in radiological non-responders. Despite the different clinical setting of metastatic tumors, we observed high expression in macrophage populations enriched in non-responders (**Fig. 6c**). The same invasion and cholesterol metabolism genes as identified in the Bassez dataset also showed higher expression in the Zhang dataset, validating our invasion program (**Fig. 6d**). Spectra thus identifies a novel pro-metastatic gene program that is upregulated under anti-PD-1/PD-L1 in therapy-resistant breast cancer patients, with implications for understanding adaptive resistance mechanisms and macrophage polarization.

## Discussion

Spectra infers interpretable gene programs from single-cell expression data by anchoring data-driven factors with prior biological knowledge. Factors that Spectra estimates on held-out data are more coherently expressed than those found by other methods, and they yield more interpretable marker lists that are not polluted by cell type markers. Upon identifying a factor, Spectra modifies it to the dataset’s biological context by upweighting novel genes that are tightly expressed with the bulk of factor genes and downweighting less coherently expressed genes. This enables Spectra to dissect programs that are highly correlated and missed by other methods, such as T cell exhaustion and T cell tumor reactivity in an immuno-oncology dataset. We demonstrate that expression of this T cell tumor reactivity program separates patients by their clonal expansion status after anti-PD1 treatment, while less coherent factors estimated by other methods fail to do so.

A unique design feature underlying Spectra’s interpretability is its unified approach for dealing with expression variance at multiple scales. Cell type heterogeneity dominates the marginal gene-gene covariance matrix, so that higher-resolution cell-type-conditional covariances exhibit markedly different structure. To address this, Spectra accepts biological prior information in the form of cell type labels, and explicitly models separate cell-type-specific factors that can account for local correlation patterns. As a result, Spectra reliably identifies programs that are conserved across multiple cell types relating to metabolism, response to cytokine signaling, differentiation and growth, while separately estimating the cell-type-specific components of these programs.

Spectra’s interpretability depends on two key components, its probabilistic model and the quality of prior knowledge used as input. We compiled an immunology knowledge base of high confidence gene sets for 50 cell types and 181 cellular processes, which can be used to improve the analysis of immune scRNA-seq data using Spectra or other supervised methods. However, Spectra does not require high-quality gene sets that are highly relevant to the dataset being analyzed as input. Spectra factor deconvolution is biologically grounded by a prediction model for gene-gene relationships that flexibly accounts for potentially noisy observations of edges in the input knowledge graph. The algorithm adaptively tunes its reliance on prior information based on concordance of the prior and observed expression data. Furthermore, novel factors are adaptively allocated when prior information is insufficient to explain the observed expression data. Exploiting this property of our method, we discovered a novel cancer invasion gene program describing an axis of variation in tumor-associated macrophages that is strongly related to anti-PD-1 therapy resistance. The association of the invasion program with non-responders to ICT replicated upon transfer to an independent dataset.

The main simplifying assumption made by Spectra and other factor analysis methods is that factors combine linearly to drive gene expression. Though occasionally violated, this approximation establishes a direct relationship between estimated model parameters and each factor’s contribution to gene expression; moreover, it avoids severe issues of non-identifiability and indeterminacy during optimization. The tradeoff is that nonlinear cooperative effects of multiple factors on gene expression cannot be determined via matrix decomposition. T cell priming, for example, represents a nonlinear interaction between TCR stimulation, coactivation and cytokine signaling, that initiates differentiation along a terminal effector memory precursor trajectory. Uncovering nonlinear relationships in an interpretable manner is a future goal of factorization.

We anticipate that Spectra will be useful in assessing the determinants of heterogeneity in large scale scRNA-seq studies, due to its simplicity, interpretability, and scalability. The strong association of some factors with immunotherapy response suggests that Spectra can delineate features that may be explored as clinical biomarkers. Spectra’s unique ability to adapt the knowledge graph to the data allowed us to infer novel candidate therapeutic targets that were added to immunotherapy response programs. In addition, the ability to transfer factors learned from one dataset to another can advance our ability to iteratively transfer and refine knowledge across scRNA-seq studies without requiring data integration.

## Methods

Full methods are provided in Supplementary Methods.

## Data availability

Count matrices of the Bassez dataset^7^ were kindly provided by the authors and are also available at (http://biokey.lambrechtslab.org). Raw read counts are available in the European Genome-phenome Archive (EGA) (EGAS00001004809, EGAD00001006608). Count matrices for the Zhang dataset^24^ were downloaded from the Gene Expression Omnibus (GEO, https://www.ncbi.nlm.nih.gov/geo/) using the following accession number: GSE169246.

## Code availability

Spectra is available as an open-source python package at https://github.com/dpeerlab/spectra and the immune knowledge base at https://github.com/wallet-maker/cytopus ^70^. Notebooks to reproduce figures are available at: https://github.com/dpeerlab/SpectraReproducibility.

## Acknowledgements

We thank Andrew E. Cornish and Samuel A. Rose for critically reading this manuscript. This work was funded by NCI Human Tumor Atlas Network grant U2C CA233284 (D.P.), U54 CA209975 (D.P.) and Cancer Center Support Grant P30 CA08748 as well as the Functional Genomics Initiative, Center for Epigenetics Research and Alan and Sandra Gerry Metastasis and Tumor Ecosystems Center at Memorial Sloan Kettering (D.P.). R.K. was supported by NSF Graduate Research Fellowship 2020297401.

## Author contributions

R.K. and D.P. conceived the algorithmic approach and strategy. R.K. developed the algorithm and coded software. R.K. and T.W. carried out testing, applications and data analysis. T.W. developed the immune knowledge base. R.K., T.W., T.N. and D.P. wrote the manuscript and D.P. supervised the work.

## Spectra Supplementary Methods

### 1 Introduction

Our goal is to use factor analysis to make single-cell RNA-seq (scRNA-seq) more interpretable by decomposing the expression matrix into a set of gene programs, where each cell is executing a subset of these programs. Gene expression is modular, where a *gene program* is a set of genes defined by a common biological task and shared regulatory mechanisms. This coregulation creates collinearity in gene expression levels, lending low-dimensional structure to high-dimensional cell-by-gene count matrices. Thus we assume each cell is executing a set of coregulated gene programs that combine linearly to define the observed gene expression. These gene expression programs are often estimated using a matrix factorization approach, which decomposes the gene expression count matrix into two matrices: (1) a set of factors *θ* (gene programs), each described by weights over the entire set of genes, and (2) a set of loadings *α* that describe each cell’s expression of each factor. A cell’s expression of any given gene is approximated by a weighted sum of factors, weighted by the cell loadings (see Fig. M1).

The parameters underlying the factor and loading matrices are chosen via an optimization process to best describe the observed data. To maintain parsimonious explanation of the observed data, each factor puts high weight on co-expressed genes and facilitates the aggregation of genes into gene programs. A successful factor analysis method should identify all active gene programs in a dataset, including variations specific to biological context as well as novel factors, and it should quantify the degree to which each gene program is executed by each cell type. A number of matrix factorization methods have been applied to estimate gene programs from single cell transcriptomic data including principal components analysis, non-negative matrix factorization, zero-inflated factor analysis, and slalom (Pierson and Yau [2015], Lee and Seung [1999], Buettner et al. [2017]). Since matrix factorization is an unconstrained problem, each individual method differs in terms of the optimization constraints and the reconstruction loss function used to estimate the factors and loadings. For example, non-negative matrix factorization (NMF, Lee and Seung [1999]) imposes a positivity constraint on both factors and loadings and can optimize either a Poisson-derived or least squares reconstruction loss. Similarly, Bayesian priors or penalty functions can be used to prioritize among solutions that explain the observed data equally well (Buettner et al. [2017], Gopalan et al. [2015], Levitin et al. [2019]) by choosing ones that are closest to the prior.

A number of aspects of single cell expression data have hindered the success of matrix factorization methods in deriving interpretable biological factors. Before developing Spectra, we scrutinized some of aspects that confound matrix factorization of scRNA-seq. First, the biggest source of variation in the expression matrix is typically driven by cell-type identity and therefore cell-type identity often dominates the derived factors. Moreover, expression covariance occurs at multiple scales and existing methods often fail to estimate factors associated with local covariance structure (e.g. cytokine responses or activation programs) within a cell-type. There are a number confounding aspects, even when applying factor analysis to a single cell-type: Important biological processes are often correlated, however recovery of correlated factors is discouraged by the reconstruction loss functions that matrix factorization methods use; which, in order to return a parsimonious explanation of the data, prioritize factorizations that account for diverse components of transcriptional heterogeneity. Furthermore, a considerable component of the variation can be attributed to either technical artifacts or highly variable housekeeping genes. Most methods can not distinguish between technical and biological variation and mix genes whose variation is driven by technical reasons (e.g. gene capture rate) with biological programs obscuring their interpretation. Even methods that use gene-set priors to guide factor analysis towards known gene programs Buettner et al. [2017] are limited by the quality of the provided gene-sets. Specifically, most gene sets are derived from bulk data and different biological contexts, whereas gene programs vary considerably across cell-types and biological contexts. To overcome these obstacles, we developed Spectra.

### 2 Spectra (Supervised pathway deconvolution of interpretable gene programs) algorithm

#### 2.1 Overview of Spectra

Spectra (https://github.com/dpeerlab/spectra) addresses these issues by grounding datadriven factors with prior biological knowledge. First, Spectra takes in biological prior information in the form of cell type labels and explicitly models separate cell type specific factors that can account for local correlation patterns. This explicit separation of cell type specific and global factors enables the estimation of factors at multiple scales of resolution. Secondly, Spectra resolves indeterminacy of the reconstruction loss function via a penalty derived from a gene-gene knowledge graph that encourages solutions that assign similar latent representations to genes with edges between them. To account for prior information of variable relevance and quality, Spectra adaptively tunes its reliance on prior information based on concordance of the prior and observed expression data. Finally, novel factors are adaptively allocated when prior information is insufficient to explain the observed expression data.

In the first step of Spectra, a set of gene-gene similarity graphs is built by aggregating information across gene sets and/or other sources. This graph representation is flexible and can accommodate various types of prior knowledge: gene sets can be incorporated into graphs by including edges between genes that are annotated to the same pathway, while existing datasets can be used to generate annotations by thresholding partial correlations or factor similarity scores. This representation lends to computational convenience as the graph dimensions are fixed regardless of the size of the input annotations. The annotations are either labeled as cell type specific, or have global scope. A separate graph is thus built for each cell type alongside a global graph.

In the second step, Spectra learns a multidimensional parameter for each cell and each gene, representing each cell and each gene’s distribution over gene expression programs. Similarity of the parameters between genes indicate that these genes are likely to have an edge joining them while similarity of the parameters between a cell and a gene indicate that the cell is likely to express that gene. Hence, the graph encodes the prior that genes with edges between them are likely to be expressed by the same set of cells. In practice, we take a number of additional steps to fulfill the desiderata: (1) factors not represented in the annotations can be discovered (2) low quality annotations can be removed (3) discrete cell types are assumed to be fixed and known and therefore not captured as factors by the model.

In order to avoid penalizing novel factors that have no relation to the annotations, we introduce a weighting matrix that scales the computation of gene-gene similarity scores by factor specific weights that are learned from the data. Factors that have low weight are not used in computing edge probabilities while factors with high weights influence the edge probabilities directly. Hence Spectra can estimate similar parameters for two genes without forcing a high edge probability between them, so long as the factors corresponding to these genes also have low weight (Fig. M5). These weights allow the addition of new, unbiased factors that are not influenced by the input annotations. Importantly, weights are estimated from the data allowing for an adaptive determination of the relative number of unbiased and biased factors. An estimated background rate of edges in the graph allows for the removal of annotations with little supporting evidence from gene expression data. Finally, Spectra explicitly separates global and cell type specific factors by enforcing a cell type determined block sparsity pattern in the cell loading matrix. Cell type specific factors capture within cell type variation while global factors capture any variation that is shared across multiple cell types. To reduce the burden of modeling constitutively expressed cell type marker genes, each factor’s contribution to gene expression is multiplied by a cell type specific gene weight. These cell type specific gene weights explain away the influence of cell type marker genes and hence mitigate the tendency of these marker genes to influence the factors themselves.

#### 2.2 Components of the Spectra objective function

Broadly speaking, Spectra fits a set of factors and cell loadings by minimizing an objective function with two components. The first component of the objective function, 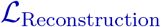, measures how well the estimated model parameters can reconstruct (or predict) the observed expression count data using the set of all model parameters, Θ. We write 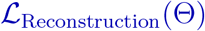 to emphasize that 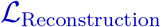 is a function that maps a set of model parameters to a corresponding objective value. The second component of the objective function measures how well the set of model parameters Θ correspond to our biological prior information. This second component is denoted 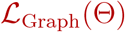. We weight this term by a user defined hyperparameter *λ* which allows a user to control the level of confidence placed in the given biological prior information. The general form of the Spectra objective function is:

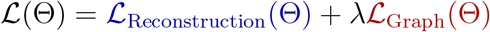

**Figure M1:**
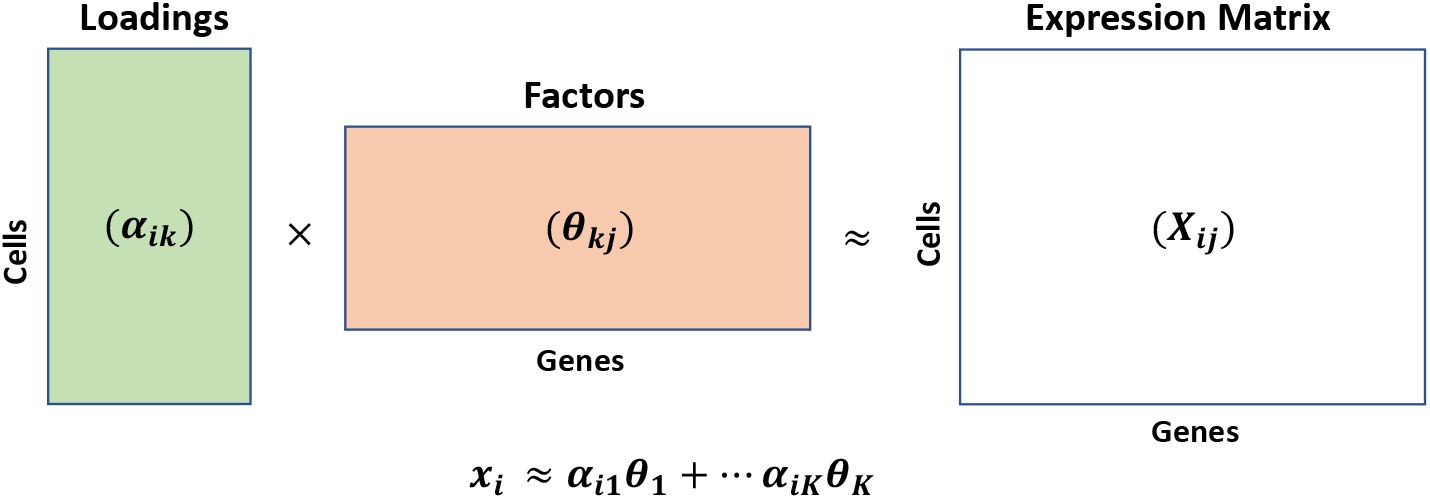
Matrix decomposition methods applied to single cell RNA sequencing data decompose the expression count matrix as a product of cell level loadings (*α_i_*) and latent factors (*θ_k_*)

Below we describe the precise functional forms of each of the objective function components.

##### 2.2.1 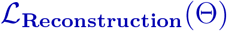: Modeling gene expression as a low rank product

We assume that the expression variation observed in the count matrix is driven by variation in the activity of different biologically meaningful gene programs, as well as technical variation which often involves highly expressed genes. Therefore, our model of gene expression needs to account for both components. In more detail, interpretation of factors estimated from single cell RNA sequencing data is often hindered by highly expressed genes, which factor analysis methods based on reconstruction loss functions must account for. Housekeeping genes required for basal cellular function such as GAPDH, ACTB, and ribosomal genes are expressed at high levels, and hence unduly influence the reconstruction loss function despite the fact that their expression variance is explained in large part by overall levels of transcription. As a result, existing matrix decomposition methods tend to put high weight on such non-specifically expressed genes; though post hoc corrections can be applied for the interpretation of individual factors. On the other hand, certain important cytokines and chemokines (e.g. IL4, IL6, IL2, IL10), receptors (CXCR1,CXCR2), and transcription factors (RORC, BATF3) are expressed in low mRNA copy number. Normalization strategies that rescale features empirically tend to amplify measurement uncertainty associated with lowly expressed genes, leading matrix factorization methods to overfit and return low quality gene expression programs. To address this, we introduce gene scale factors *g_j_* that are estimated from the data and allow the model to explain high expression and variability of certain genes without increasing the magnitude of these genes’ factor weights. Since lowly expressed genes are correspondingly noisier we bound the minimum gene scale factors below by a tuning parameter *δ*.

By way of notation, **X** refers to the processed gene expression matrix, with entry **X**_*ij*_ containing the gene expression value for cell *i* and gene *j*. The matrix **X** has *n* rows (the number of cells) and *p* columns (the number of genes). *K* refers to the number of gene expression programs unless otherwise specified. Additionally, for a given cell indexed by *i* the cell loading, a set of weights across the set of factors, is denoted by *α_i_*. The distribution across factors for gene *j* is denoted as *θ_j_* which sums to 1 over *K* gene expression programs, 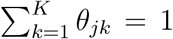. Unsubscripted variables refer to the collection containing all possible subscripts, e.g. *θ* refers to the collection of all *θ_j_*. The base expression model describing the gene expression measurement for cell *i* and gene *j* is:

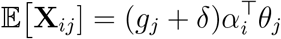

with *g_j_* ∈ [0, 1] a gene scaling parameter, 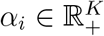 and *θ_j_* ∈ Δ^*K−*1^ (where Δ^*K−*1^ is the set of positive *K* vectors that sum to 1). The low rank decomposition of this expression model can be visualized in Fig. M2.

**Figure M2:**
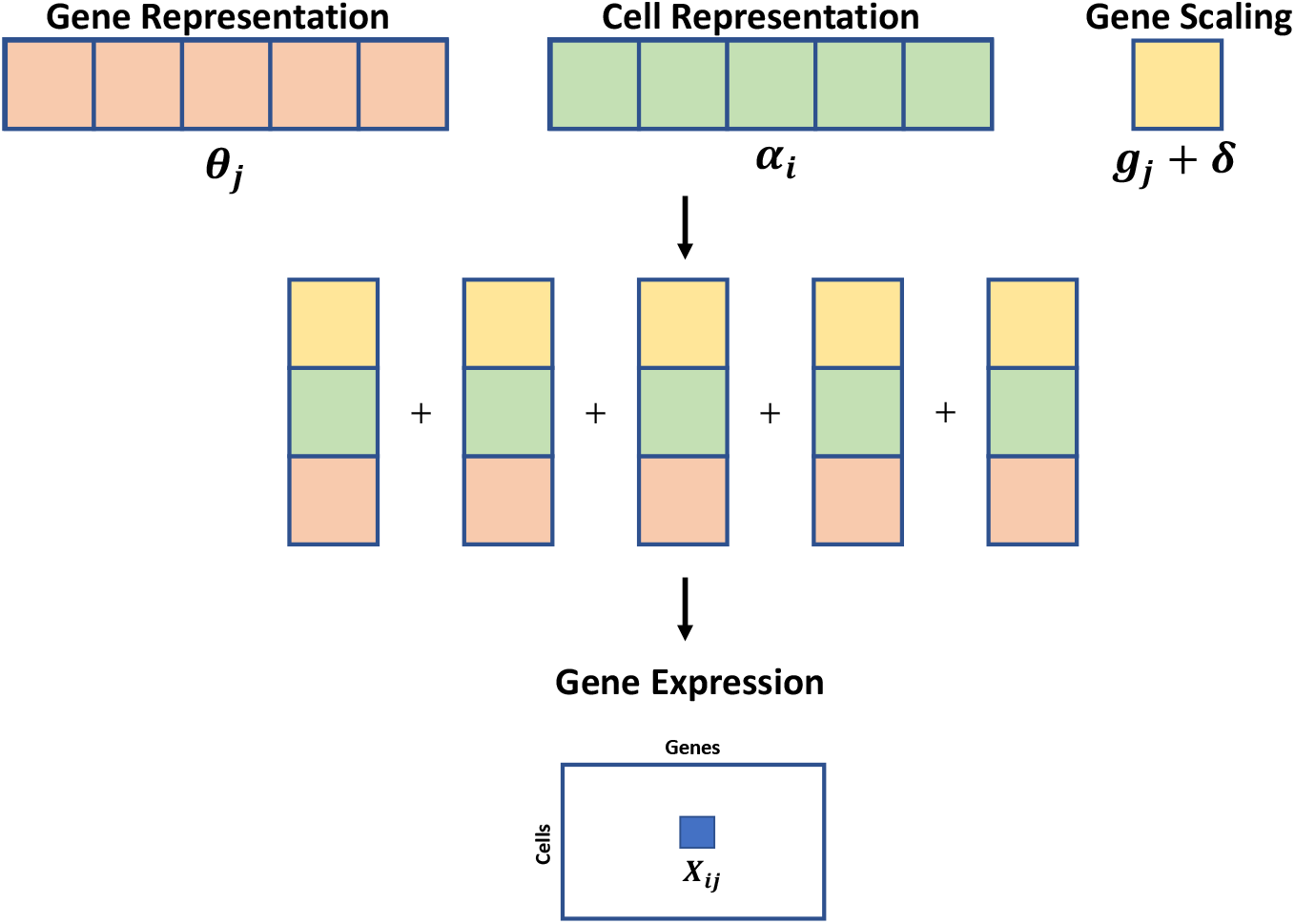
Backbone of the Spectra expression model. Each element of the expression matrix **X**_*i,j*_ is modeled as an inner product of gene representations (*θ_j_*) and cell representations (*α_i_*), weighted by gene scalings *g_j_* + *δ*

###### Incorporating cell types into modeling expression variation

Because expression variation is dominated by cell types, existing methods generally fit factors that are polluted with cell type markers or alternatively must be run on a subset of the data. For example, T cell receptor activation programs - consisting of markers such as NFATC1 and NFATC2 - are confounded with T cell identity and existing factor analysis methods tend to return identity markers such as CD3, CD4 and CD8. Similarly, programs representing metabolic pathways are often confounded with plasmacytoid dendritic cell (IL3R, BDCA2) or B cell identity markers (CD19, CD79A). While it is challenging to fit a biologically meaningful factor model, successful cell-typing of scRNA-seq data is a solved problem. Therefore, to mitigate this issue, Spectra assumes that discrete cell types are known and therefore not captured as factors by the model, instead Spectra explicitly fits cell type specific and global factors - allowing Spectra to effectively deal with expression variance at multiple scales. To perform this cell type integrative factor analysis, for cell type *c* and cell *i* the model is extended to:

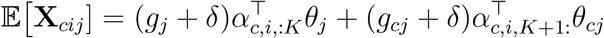

where *c* is the cell type label for cell *i*, *g_cj_* is cell-type specific gene scaling, 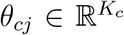 is a cell type specific gene representation with 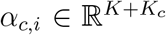. Single subscript variables such as *g_j_* and *θ_j_* denote global parameters, while the notation *α*_:*K*_ indicates the first *K* elements of a vector (typically denoting global elements) and *α_K_*_+1:_ indicates the tail of the vector from the *K* + 1’st element (typically denoting cell type specific elements). The threshold *δ* restricts the maximum ratio of gene scaling factors to 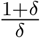.

**Figure M3:**
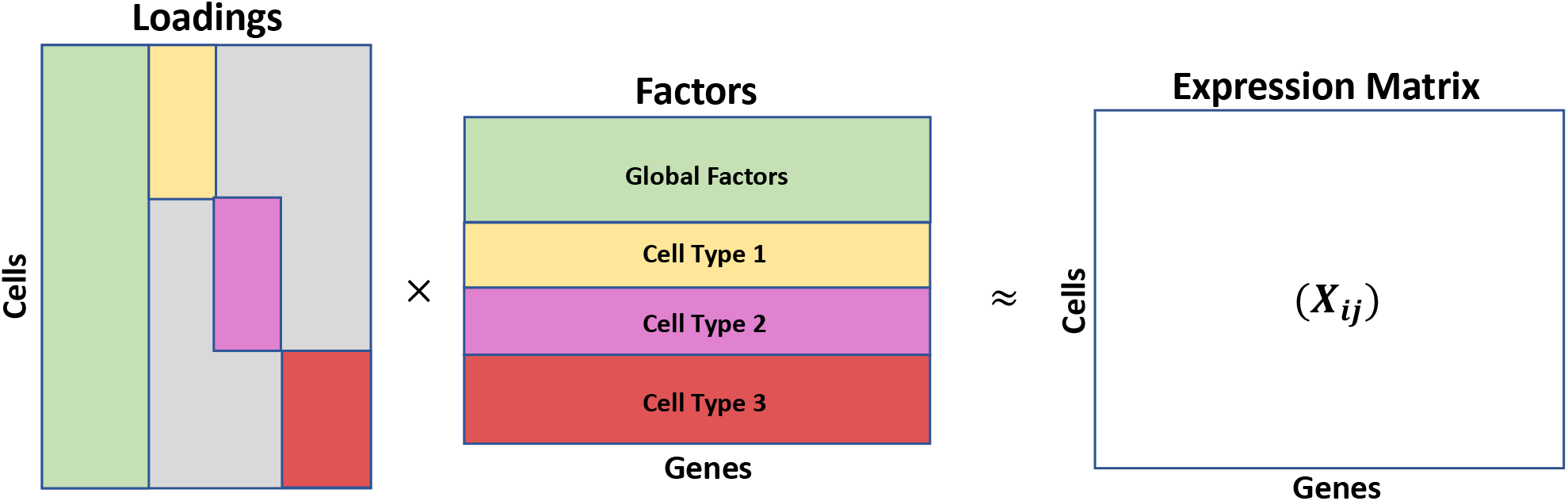
A matrix decomposition method that fits cell type specific and global factors. Green colored global factors correspond to green colored loadings which are potentially non-zero for all cells, while the three cell type specific factors have sparsity pattern in their loadings corresponding to cell type identity. The model still decomposes the expression count matrix as a product of cell level loadings (*α_i_*) and latent factors (*θ_k_*)

Spectra models the presence of gene programs with highly limited scope in that they can only be activated by a specific cell type, which can be represented by a hard coded sparsity pattern in the cell loading matrix (Fig. M4). The cell type specific gene scalings (*g_cj_*) associated with these programs are encouraged to capture cell type identity markers and constitutively active genes, enabling factors themselves to capture variation across cell types and within cell types (Fig. M3). Spectra tends to assign constitutive genes such as EEF1A1 and ACTB as well as identity markers such as CD4 and CD3 high values of *g_j_*. Lowly expressed genes important for CD4 T cell specific gene programs such as IL21, IL13, and IL6 are often assigned small values *g_j_*, which allows Spectra to attend to gene expression differences that occur on a smaller scale (Fig. M3). By default, Spectra runs with at least one cell type specific factor per cell type so that global factors do not capture cell type identities.

###### Determinating Cell Type granularity

Spectra can accommodate cell type labels at any level of granularity, subject to a linear increase in computational burden with the number of cell types in the dataset. Additionally, as the granularity increases the effective sample size for estimating cell type specific factors decreases, leading to potentially lower quality cell type specific factors. The correct cell type granularity depends on the dataset and the specific scientific questions at hand. First, the analyst should incorporate cell types that are known to be discrete and easily identifiable in the dataset via standard clustering analysis (e.g. T cells, B cells, myeloid cells, and epithelial cells). If cell subtypes exist that are not included as input to the model, Spectra devotes factors to describing variation across these subtypes. Moreover, if intermediate differentiation states between subtypes exist in the data, these subtypes should generally not be included as input to the model because (1) coarser cell type specific factors can describe these intermediate states and (2) delineating between subtypes via clustering may be inaccurate.

##### 2.2.2 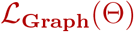: Modeling gene-gene relationships in relation to expression data

In addition to faithful approximation of the input count matrix, we would also like interpretable factors that correspond known gene programs and biological processes (prior). Therefore, the second component of our likelihood function is a penalty term that guides the solution towards this prior. A key novelty of Spectra is that it models this prior knowledge as a gene-gene community graph, which provides both computational efficiency and flexibility to adapt the graph structure to the data. In this graph nodes represent individual genes and edges between genes occur when each gene has a similar distribution over factors. Then communities within the graph, or densely connected subsets, represent gene programs while edges between communities contain information about genes that participate in multiple gene programs. Providing an imperfect, partially known graph structure as input, we can constrain our matrix factorization solution to respect the structure to yield interpretable gene programs. A main advantage of this approach is its flexibility. Gene sets are naturally incorporated into a graph by forming fully connected cliques among members of each set. Further, more complex prior knowledge graph structures can be used as input, for example, arising from gene programs estimated from a separate dataset or cell atlas. Most importantly, this the structure of this input gene-gene graph can be improved, by fitting it to the data and learning gene programs that are more faithful to the data.

**Figure M4:**
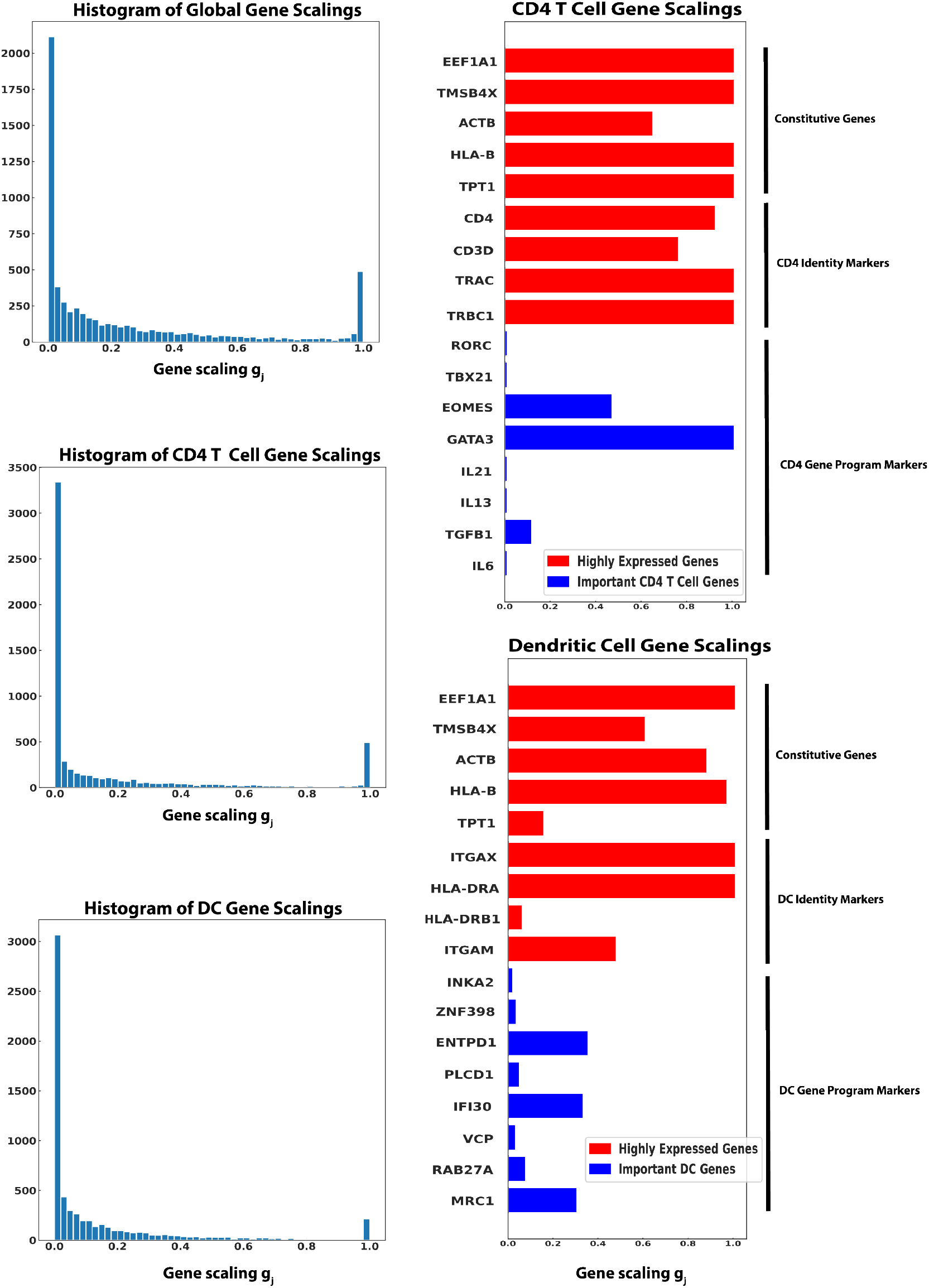
*Left* : Distribution of gene scale factors for global (top), CD4 T cells (middle), and dendritic cells (bottom) on Bassez data (*λ* = 0.01, *δ* = 0.001). Mode at 1 + *δ* is due to constraint on maximum ratio of scale factors. *Right* : Gene scale values for specific genes for CD4 T cells (top) and dendritic cells (bottom) including constitutive genes, identity markers, and genes involved in cell type specific gene programs

**Figure M5:**
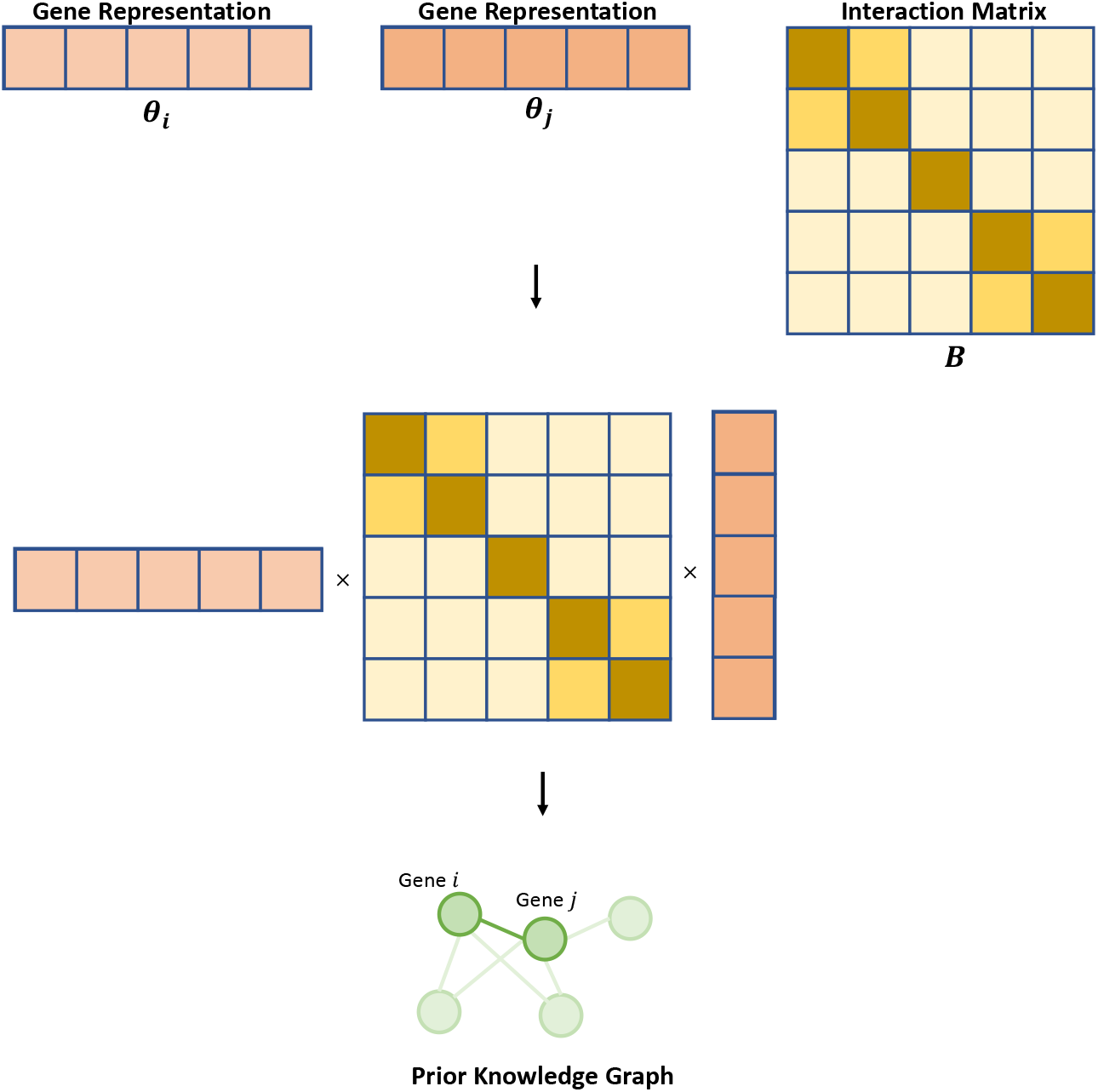
Backbone of the Spectra graph model. Spectra incorporates prior information by encouraging gene representations to be able to predict a set of annotated gene pairs, which can be represented by a graph. Given gene representations, *θ_i_* and *θ_j_*, an edge is predicted by a weighted inner product, weighted by interaction matrix *B*, 〈*θ_i_, Bθ_j_*〉

A second advantage of the graph prior is its scalability. While gene sets may be highly overlapping, especially when curated from several separate databases, this redundancy is eliminated when storing information at the level of gene-gene relationships. Redundant gene sets will be merged into highly overlapping communities, and so two redundant gene sets can be approximately described by a single factor. A further computational advantage over gene set priors is that the dimensions of the graph are fixed as the size of gene set database increases, with only the number of edges increasing and eliminates the need for iterating over the gene set dimension. Finally, operations involving the graph are implemented via efficient and parallelizable matrix multiplications with the graph adjacency matrix, thus allowing Spectra to efficiently scale to a large number of gene-sets and cells (Fig. 3D)

In order to encourage factors to capture our prior knowledge of gene programs, we assume that binary gene-gene relationships are evidence of a pair of genes having similar latent profiles. This assumption could be incorporated by assuming a model for edge probabilities depending on the similarity scores 〈*θ_i_, θ_j_*〉 for genes *i* and *j*. However, the naive inner product does not explicitly account for the fact that prior information is invariably imperfect in systematic ways. First, at the level of entire gene programs: not all gene programs are active in all datasets and therefore entire graph communities may be unnecessary for describing the observed expression data, while there are likely novel gene programs observed in the expression data that are not be represented by communities in the graph. Also gene programs are imperfect, both due to inaccuracy of annotation and more frequently, gene programs differ across biological contexts and our prior information is typically derived from a different biological context. Therefore, genes may be misclassified into gene sets to which they do not belong (corresponding to noisy edge observations) or gene sets may be incomplete (corresponding to missing edges). Spectra addresses these issues in two ways: (1) adaptively modeling background noise in the graph, allowing for the addition and removal of edges (Section 2.2.5) and (2) tuning the weight of the prior gene-gene matrix through the incorporation of a weight matrix, termed the factor interaction matrix, into the inner product between gene representations *θ_i_* and *θ_j_* (Section 2.2.3).

**Figure M6:**
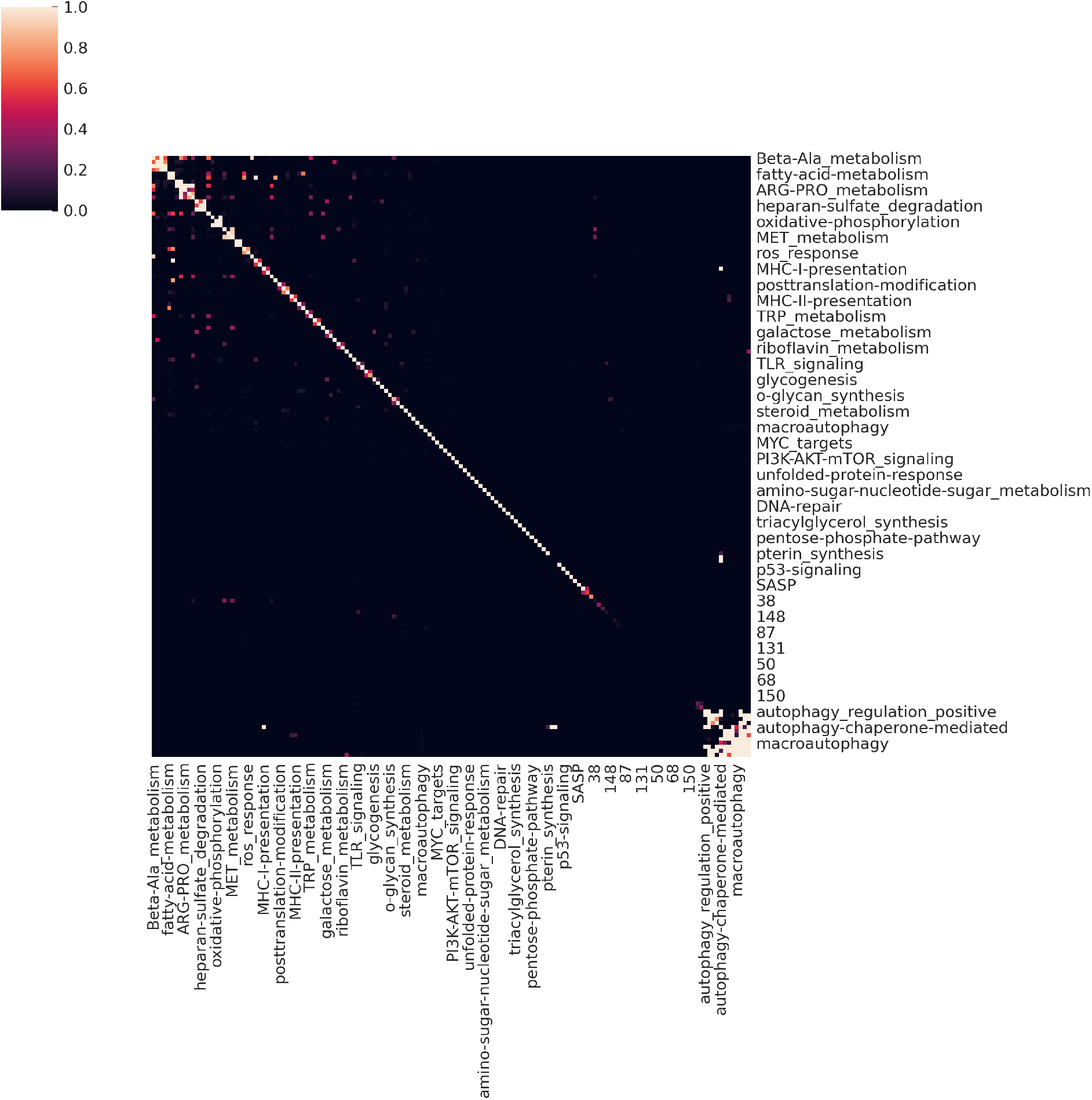
Global factor interaction matrix (*B*) estimated from the Bassez dataset. Gene programs with 0 values on the diagonal of the matrix are novel.

##### 2.2.3 The factor interaction matrix tunes the weight of the gene-gene prior

To understand the purpose of the factor interaction matrix, let’s first consider the ordinary inner product, measuring gene-gene similarity in terms of gene program representations:

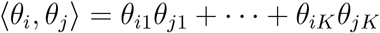

The maximum value of this product is 1, achieved only when gene *i* and gene *j* put all their weight into a single gene program. Consider what happens if genes *i* and *j* are important components of a gene program that exists only in the expression data and not in our prior information. Then *i* and *j* are not connected in the graph and so the inner product model encourages 〈*θ_i_, θ_j_*〉 ≈ 0. When 〈*θ_i_, θ_j_*〉 ≈ 0, gene *i* and *j* must be components of entirely separate programs. In this way, we see that the naive inner product discourages new factors from being estimated from the expression data. Such an inner product model estimates novel factors that are heavily biased by the graph.

Now instead of the naive inner product consider a weighted product, weighted by scalar values (*b*_1_, *b*_2_,…, *b_K_*) that are between 0 and 1:

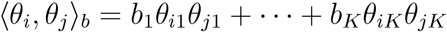

In order to model the data we can adjust the values of (*b*_1_,…, *b_K_*) to achieve the best fit. Consider the same situation as above, where *i* and *j* are not connected in the graph but they are components of a gene program supported by expression data alone. The product model again encourages 〈*θ_i_, θ_j_*〉_*b*_ ≈ 0; however, now this constraint does not necessarily encourage *θ_i_* and *θ_j_* to be dissimilar. To see this, suppose that *θ_i_* = [1, 0, 0] and *θ_j_* = [1, 0, 0]. If *b*_1_ = 0, then:

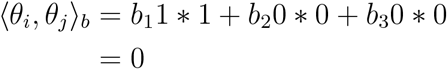

Hence, novel gene programs can be estimated so long as the value of *b_k_* corresponding to that program is pushed towards 0. We can interpret gene programs corresponding to low values of *b_k_* as novel and gene programs corresponding to high values of *b_k_* as supported by prior information. We could equivalently write each weight *b_k_* as one of the non-zero elements of a diagonal matrix

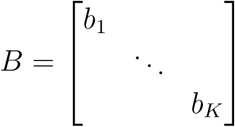

so that

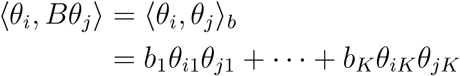

In practice, we allow the off diagonals of this matrix *B* to be estimated as non-zero (Fig. 5). The resulting matrix is termed the factor interaction matrix.

Allowing off diagonals of the factor interaction matrix to be non-zero serves two purposes: First, it allows the model to explain overlapping gene sets without forcing shared genes to have partial membership. For example, if two gene sets overlap but in reality represent two distinct biological processes that can be separated in the gene expression data, the model is not forced to assign partial membership to overlapping genes but can fully assign genes to one of two programs. To account for this, the off diagonal element corresponding to this pair of gene programs (*B_k,l_* for programs *k* and *l*) can be estimated as greater than 0. On real data, we see this occur for *β*-alanine metabolism and fatty acid metabolism (Fig. 6). Second, non-zero off diagonal elements of the factor interaction matrix serve to mitigate the effect of low quality edges in the prior graph by allowing edges between genes that are in separate gene expression programs to arise with non-zero probability.

#### 2.3 Full Spectra model

As notation we refer to the adjacency matrix of an input graph as *A* ∈ ℝ^*p×p*^ with element *A_ij_* = 1 if an edge exists between *i* and *j* and *A_ij_* = 0 otherwise. Following the discussion above, the Spectra generative model states (Fig. M5):

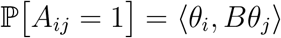

In the full Spectra model, each gene has a separate representation per cell type (in addition to its global representation), *θ_ci_*, where *c* indexes into the possible cell types. In order to supervise these representations in a cell type specific manner, the user (optionally) provides one graph for each cell type and a graph representing global gene-gene relationships (Fig. M7). These graphs are modeled separately - where each graph’s edges can only be predicted using factor representations specific to that cell type. The cell type specific graphs are denoted *A_c_* for cell type *c*, with *A_c,ij_* = 1 if there is a cell type specific annotation between genes *i* and *j* for cell type *c*. The cell type specific graphs can only influence cell type specific factors and vice versa:

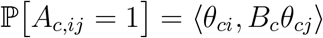

diagrammed in Fig. M7. Importantly, a separate factor interaction matrix, *B_c_*, is learned for each cell type with a prior graph provided.

The computational cost of including granular cell type specific prior information can be large, as each cell type requires its own graph.

##### 2.3.1 Background edge rates

Realistic annotation graphs have a number of edges that are not supported by expression data, and the model should be allowed the flexibility to attribute edges (or the lack thereof) in annotations to a background rate of noise. To allow flexibility in modifying the original graph we incorporate background edge and non-edge rates *κ* and *ρ* that reflect noise rates in the observed graph. These parameters serve two separate purposes: first, they deal with numerical stability issues by moving probabilities away from 0 and 1, and second they control the rate that edges are added and removed from the original graph. Intuitively, our inference procedure examines whether a relationship (or lack of a relationship) in the prior knowledge graph is consistent with expression data, and if not can ascribe this relationship to random noise.

The generative process of our model is that with some probability *ρ*, edges between gene *i* and *j* are blocked out and cannot occur irrespective of the corresponding factor values *θ_i_* and *θ_j_*. If this doesn’t occur, an edge will be generated by random chance with probability *κ*. Finally, if neither of these events occur, an edge is generated according to the factor similarity score 〈*θ_i_, Bθ_j_*〉. This yields the following distribution for the adjacency matrix:

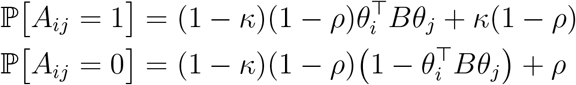

where *κ* and *ρ* are (cell type specific) background rates of 1 and 0 in the adjacency matrix respectively. *κ* and *ρ* can be estimated from the data or fixed to constants and treated as tunable hyperparameters.

**Figure M7:**
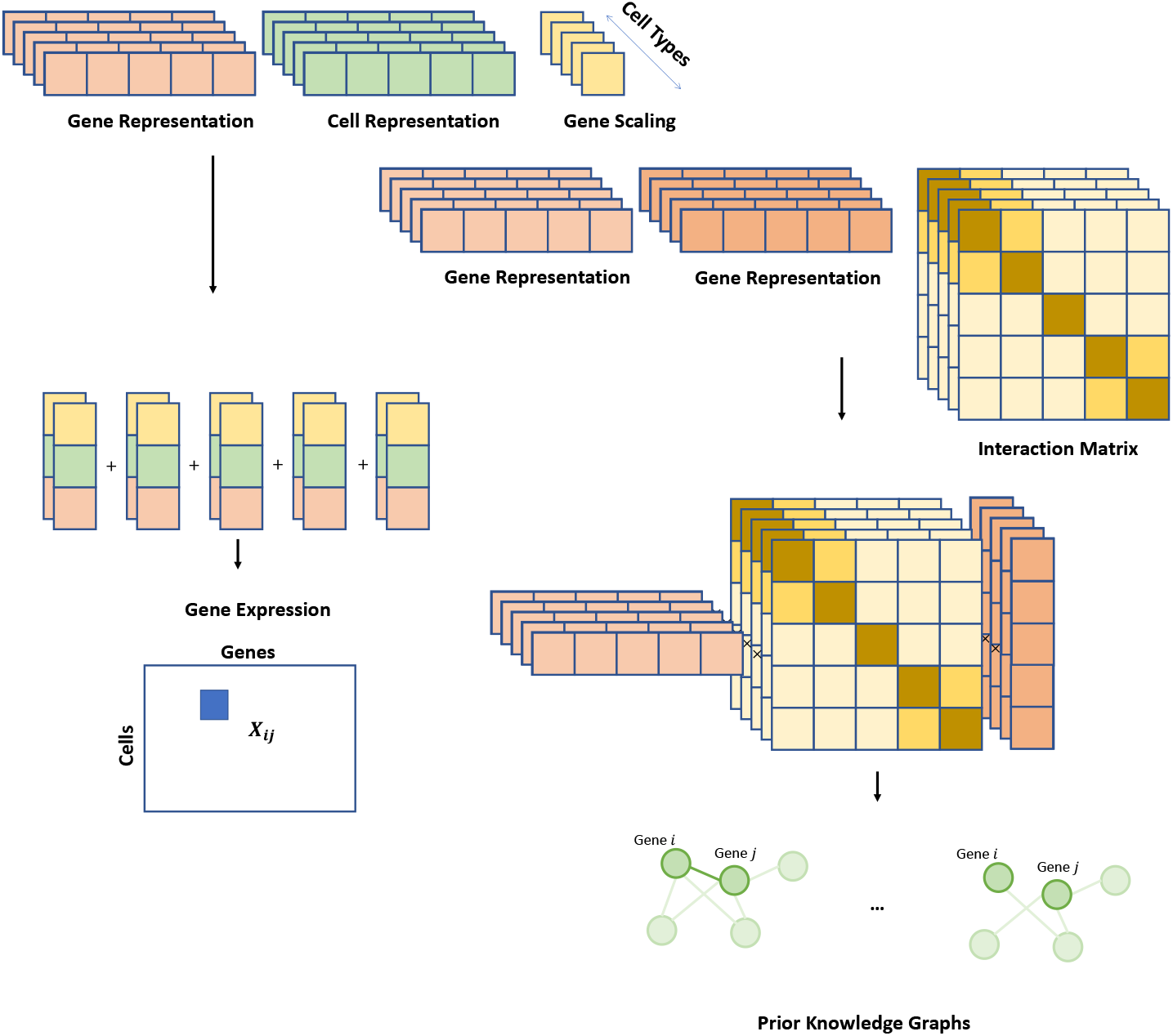
Spectra model. The full Spectra model combines the components described in Section 2.2.2. and Section 2.2.3. In addition to it’s global representation, each gene is also equipped with a per cell type representation. Each cell is also equipped with an additional per cell type representation that is non-zero only for the cells assigned cell type. The individual per cell type arms are aggregated into a single gene expression estimate for each cell and gene. For each cell type that has a prior information graph, a weighted inner product weighted by cell type specific factor interaction matrices estimates the edge probability for each pair of genes.

##### 2.3.2 Constructing the gene-gene prior graph

In most applications, Spectra receives a set of gene-sets, rather than a gene-gene graph as input and the gene-gene graph is constructed from these gene-sets. Large gene sets generally provide lower evidence that any given gene is crucial to the process that the gene set represents. For example, hallmark gene sets often contain hundreds of genes Liberzon et al. [2015], some of which are upregulated as distant downstream targets. Additionally, larger cliques represent a larger component of the likelihood function, potentially biasing Spectra solutions towards attending to the largest gene communities. Therefore, by default, when Spectra takes in gene sets as input, the edge weights used to down-weight the contribution of any individual graph edge proportionally to the size of the gene set that it’s derived from. The default weighting scheme is to weight edges by the total number of edges in the clique. For a given gene set *G_k_*, this involves downweighting by 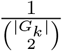:

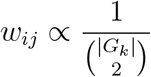

where |*G_k_*| is the size of a gene set *G_k_* containing genes *i* and *j*. The weights are rescaled so that the median weight across gene sets is 1. When a pair of genes exists in multiple gene sets, the weights accumulate additively. Another reasonable choice is 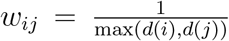, where *d*(*i*) is the degree of node *i*.

As an alternative weighting scheme, Spectra accommodates weighted graphs by scaling edges by edge specific weights. This feature allows users to annotate the prior information graphs with additional quantitative information representing relative confidence in each individual annotation.

##### 2.3.3 Likelihood function

The heretofore described model components describe the expected values of the expression data matrix, **X**, and the prior knowledge graph, *A* under the Spectra generative process. Together with specific observation distributions, this specifies a likelihood function that serves as the maximization objective of Spectra, fit via either first order methods or expectation maximization. For ease of exposition, we first describe the likelihood function assuming a single cell type. Recall the general form of the Spectra objective, consisting of a term that measures the ability of Spectra factors to recapitulate expression data and a term that measures the concordance of Spectra factors with the prior knowledge database:

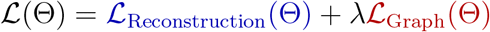

As edges are binary, combined with the assumption of independence, the log likelihood of *A_ij_* given a probability of 1, *p_ij_* ≔ ℙ[*A_ij_* = 1], is:

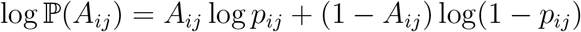

With *p_ij_* as described in Section 2.2.2. and 2.2.3.,

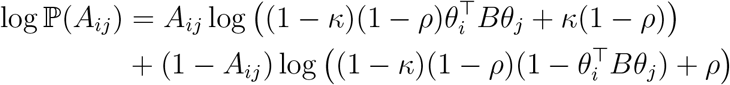

To incorporate weights (following Section 2.2.4), we weight likelihood terms corresponding to each edge in the graph by an edge specific weight *w_ij_*:

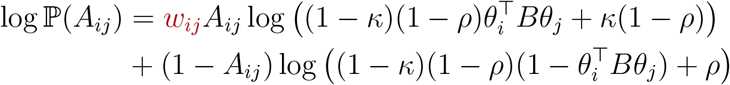

Combining across all observations (*i, j*), this leads to the expression for 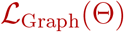 :

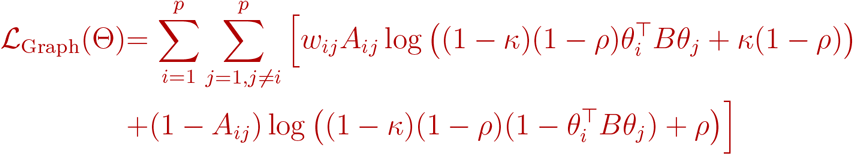

To model expression data, we use a Poisson observation model. The loss function derived from the Poisson distribution has been used widely for modeling single cell RNAseq counts Dann et al. [2022], L Lun et al. [2016], Levitin et al. [2019]. Here, we are primarily concerned with how the loss function behaves under changes in scale. For example, suppose we have an estimated expression value 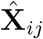. We can write 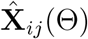 as our predicted gene expression as a function of the model parameters. The least squares loss 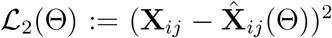 is quadratically dependent on the scale of **X**_*ij*_, since replacing both ground truth and estimate by scaled versions *φ***X**_*ij*_ and 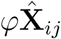 leads to a loss of 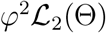. Similar to the issues addressed in 2.2.1, the squared loss function encourages factors to attend to highly expressed genes, since scale differences amplify the loss quadratically. At the other extreme, consider the Itakura-Saito loss (IS loss), given by (we briefly assume both ground truth and estimate are not 0):

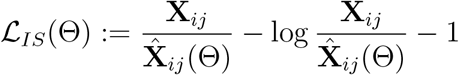

If we scale observed counts and prediction, *φ***X**_*ij*_ and 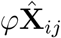, then the IS loss does not change. So, matrix factorization with the IS loss does not suffer from a bias towards highly expressed genes. However, forcing the model to predict all lowly expressed genes is not desirable - often leading to low quality factors. The Poisson log likelihood exhibits a practically convenient balance between these two extremes:

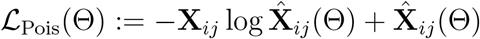

When **X**_*ij*_ and 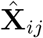 are scaled by *φ*, 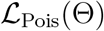 is scaled by *φ*. This linear dependence on expression scale achieves a good balance in the relative weighting between highly expressed and lowly expressed genes.

An additional advantage of this loss function is that the second term behaves as a lasso penalty (Tibshirani [1996]), inducing sparsity in the resulting estimates of 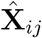 for sparse data **X**, noted by Gopalan et al. [2015]. This sparsity allows for a parsimonious explanation of a cell’s gene expression using as few factors as possible. In Spectra, we have: 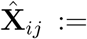 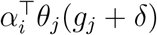, yielding the expression:

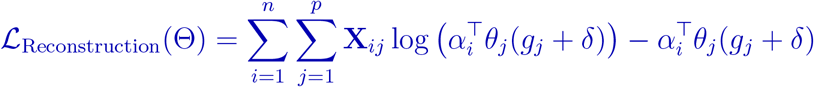

Combining the components, the log likelihood function is:

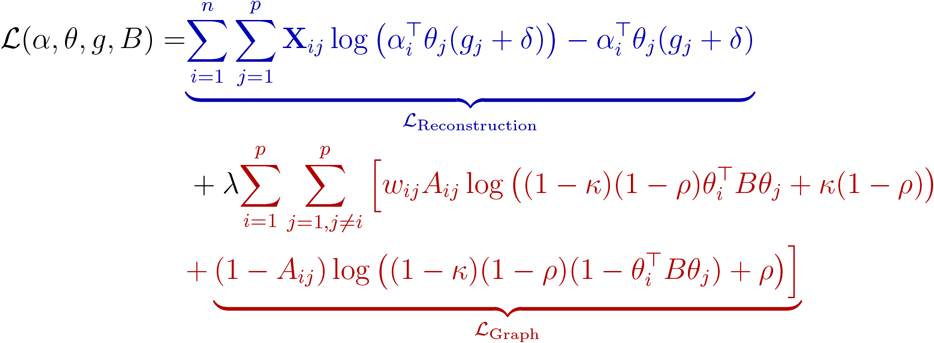

Again, **X** is the data matrix after processing, whereas (*α, θ, g, B*) are the four model parameters that need to be estimated. The first term in the likelihood function comes from the log likelihood of the Poisson distribution (also referred to as the KL Divergence loss function when multiplied by −1) while the second term is the log likelihood of a Bernoulli distribution with observations *w_ij_A_ij_* and 1 − *A_ij_* (the observations are no longer binary after scaling by edge weights). The likelihood function optimized by Spectra includes an optimization over cell type specific and global parameters and so an additional sum over cell types is included in the log likelihood (Fig. M7).

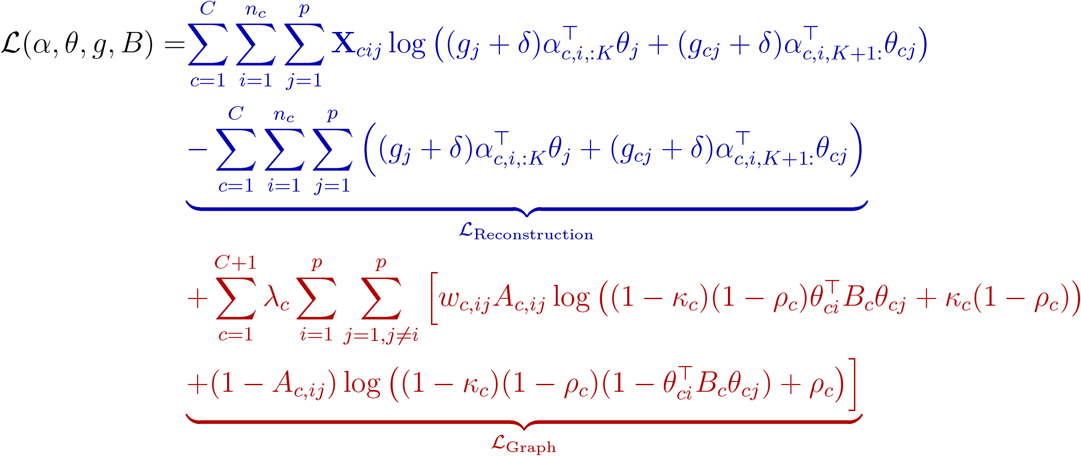

As all discrete parameters have been integrated out, this log likelihood can be directly maximized via first order methods such as gradient descent. Approximate second order methods are not ideal, due to the high dimension of the parameter space for practical problem sizes. However, for smaller sized problems (in terms of number of genes and factors) we develop an expectation maximization approach that yields intuitive coordinate ascent updates of model parameters.

#### 2.4 Spectra’s output

To describe the activity level of factor *k* in cell *i* we compute cell scores as `cell_score`_*ik*_ = *q_k_α_ik_* where 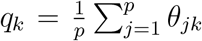. In other words, the cell scores are the loadings weighted by the total factor usage across all genes. This allows us to circumvent the non-identifiability of scale associated with factor analysis approaches.

To describe the relevance of gene *j* for factor *k* we compute gene scores for gene *j* and factor *k* as 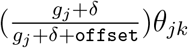. The first term is near 0 when *g_j_* is very small and near 1 when *g_j_* is large. This allows us to remove very lowly expressed genes from the factors while maintaining coherence. By default, the `offset` term is set to 1 can be tuned and in some cases set to 0, which yields the factors *θ_jk_* themselves. Each *θ_jk_* is more directly influenced by the prior than *g_j_θ_jk_* and so setting `offset` to 0 tends to yield marker lists closely resembling input gene sets.

Users can access additional parameters that facilitate interpretation of the gene scores and cell scores. The factor interaction matrix per cell type (*B*, Fig. M6) contains entries in the range [0, 1], where diagonal entries can be interpreted as the relevance of a given factor to the prior graph. Off diagonal entries can be interpreted as a background rate of edges between genes that are expressed in separate factors. For each cell type, users can access a posterior graph that is denoised using information from the expression data. The posterior graph is computed by the inner product 〈*θ_i_, Bθ_j_*〉 for each pair of genes *θ_i_* and *θ_j_* after estimating *θ_i_, θ_j_* and *B* from the data.

#### 2.5 Optimization

We develop two optimization schemes: an auxiliary latent variable expectation maximization (EM) approach and gradient descent based optimization via Adam (Kingma and Ba [2014]). EM converges quickly in many situations; however, the memory requirements are substantially larger than the gradient descent based optimization. Specifically the memory requirement of EM parameter storage is *O*(*npK* + *p*^2^*K*^2^) due to auxilliary parameter storage while the memory requirement of gradient descent is substantially lower: *O*(*nK* + *pK* + *K*^2^).

Though memory intensive, the EM solution is valuable for two reasons: (1) for problems with a small number of factors (< 20) and genes (< 2500), EM is fast and less sensitive to initialization than gradient descent (Salakhutdinov et al. [2003])(2) the EM updates are intuitive and give us understanding of how our algorithm balances evidence from the graph and expression data.

On the other hand, optimization with Adam can handle a large number of factors (> 200) and genes (> 10000), and can exhibit stability with the appropriate initialization. By default Spectra uses Adam for optimization.

##### 2.5.1 Expectation Maximization

For ease of exposition we describe the EM (expectation maximization) routine for the non-integrative model; the updates are easily extendable to incorporate cell type labels. To make expectation maximization possible, we exploit two facts about the distribution of (**X**, **A**) Liu and Wu [1999]. The first is that if *z_ijk_* ~ Pois (*g_j_* + *δ*)*α_ik_θ_jk_* and we define 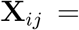 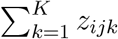, then **X**_*ij*_ still has the correct marginal distribution, due to standard properties of the Poisson distribution Gopalan et al. [2015]. Secondly, if we define 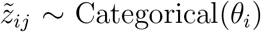, 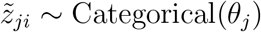 and define a conditional distribution for *A_ij_* as:

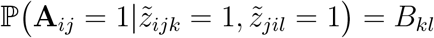

then **A**_*ij*_ still has the correct marginal distribution Airoldi et al. [2008]. As a result, we can optimize the marginal log likelihood via optimization of the expected complete data log likelihood 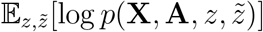 where the expectation is taken over the posterior 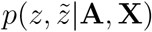.

The expected complete data log likelihood is given by:

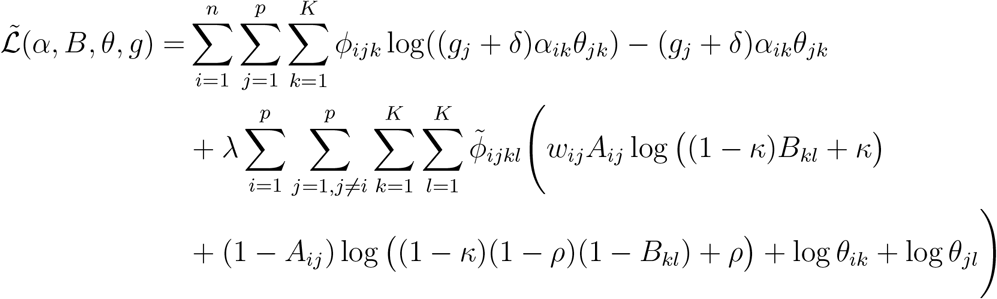

where 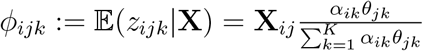 and

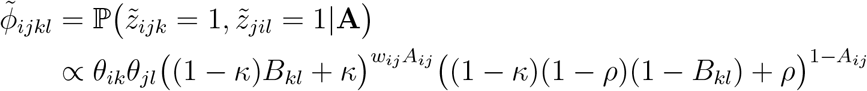

Importantly, this manipulation moves summations outside of the logs which permits analytic EM updates for *B*, *α* and *g* given by:

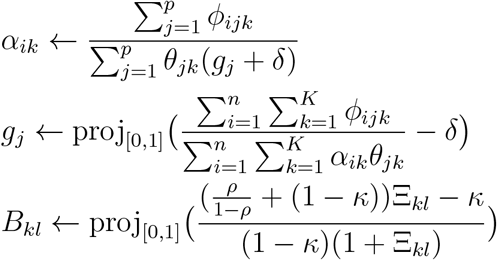

where 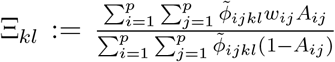, representing an odds ratio between Bernoulli outcomes.

Further, the complete data log likelihood has diagonal Hessian when viewed as a function of *θ* only, 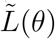, permitting linear time Newton Raphson updates:

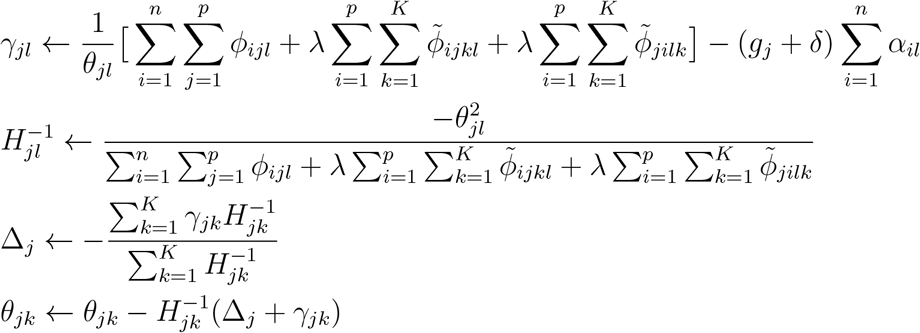

The integrative version of Spectra uses analogous updates, with the bounds of summations appropriately modified; specifically the E step updates are 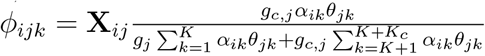 for cell type specific factors and 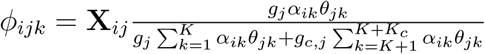 for global factors.

**Algorithm 1.**
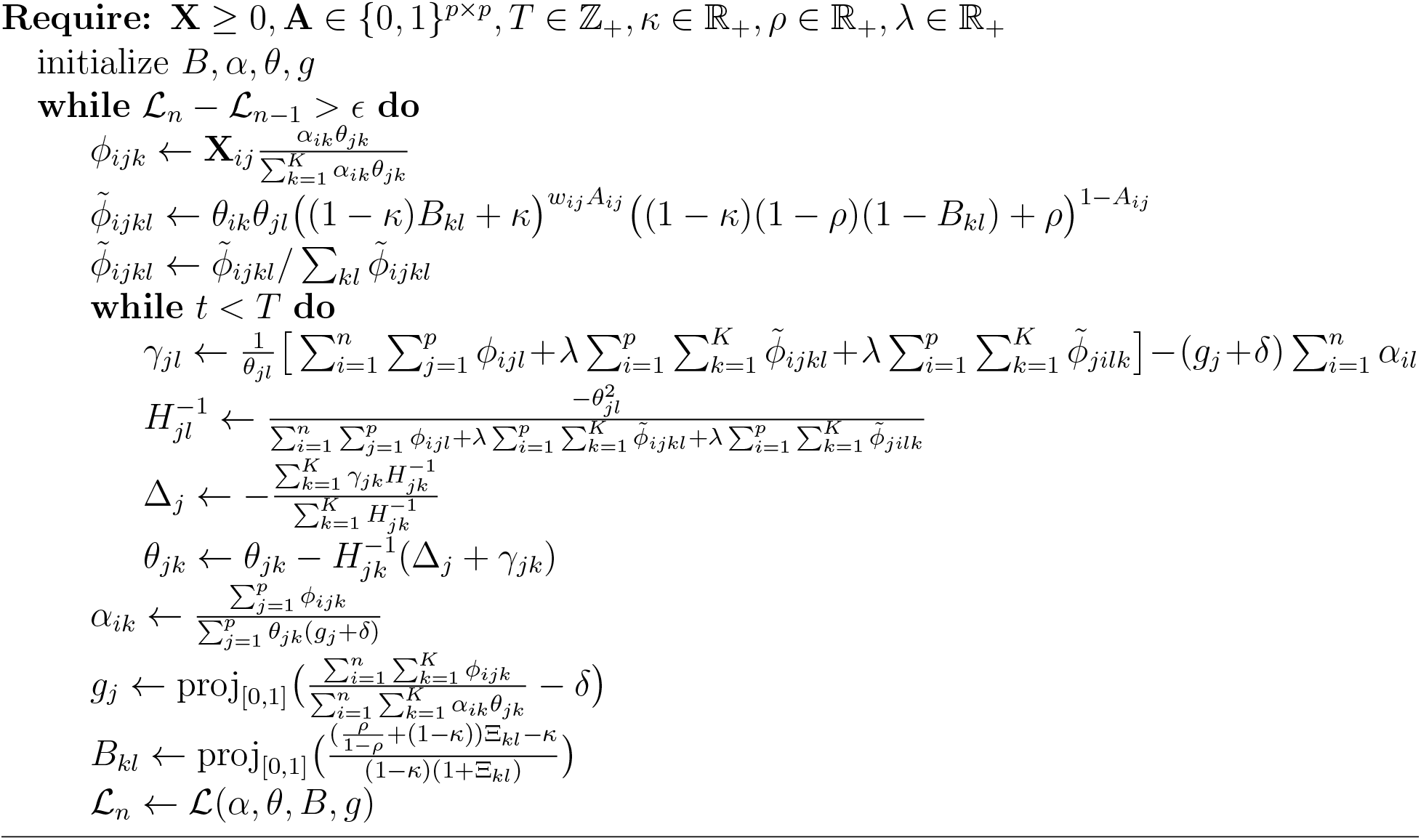
EM-`Spectra` routine

##### 2.5.2 Adam

For large single cell RNA-seq datasets, the memory requirement of EM for fitting a large number of factors is prohibitive. We optimize the marginal log likelihood with Adam Kingma and Ba [2014], a momentum based gradient descent optimizer implemented in `pytorch`, directly. In detail, the Adam hyperparameters *β*_1_ and *β*_2_ are set to default values 0.9 and 0.999 respectively. We use a learning rate schedule of [1.0, .5, .1, .01, .001, .0001] where training at subsequent learning rates occurs after convergence at higher learning rates. A maximum number of iterations is fixed to 10, 000. This default training scheme can be modified by the user. In particular, for faster convergence either the maximum number of iterations can be made smaller or the smallest learning rates can be removed, allowing for solutions that are not as fine tuned.

##### 2.5.3 Initialization

Since the Spectra objective function is non-convex and susceptible to suboptimal local maxima, initialization plays an important role in the quality of the eventual solutions. When Spectra is provided with gene sets as input, our strategy is to initialize factors as close to the gene sets as possible. Whenever the number of factors is greater than the number of gene sets, we resort to a gene set based initialization procedure:

First, a hyperparameter *t* controls the strength of the initialization. By default, *t* is set to 25. For a given cell type, whenever the number of factors is at least as large as the number of gene sets, we initialize log *θ_ij_* ← *t* when gene *i* belongs to gene set *j* for each gene set *j* = 1,…, *N*_gs_ and *N*_gs_ is the number of gene sets. Further, the factor interaction matrix is initialized with logit*B_jj_* ← *t* to encode the knowledge that this factor corresponds to a gene set. To encourage the last factor to capture genes that have no edges in the prior graph, we initialize the last row and column to small values, logit*B_K,j_* ← −*t* and logit*B_j,K_* ← −*t* for all *j* = 1,…, *K*. Corollary 1 explains why this leads to extremely fast convergence when *λ* is large.

For a given cell type, when the number of factors is not greater than the number of gene sets we resort to initialization with non-negative matrix factorization (Lee and Seung [1999]).

#### 2.6 Determining the number of factors

We adopt two approaches to determining the number of factors. The first is to set the number of factors for each cell type equal to the number of gene sets available for that cell type +1 (similar to the approach taken by slalom), and the second is to estimate the number of factors from the data via bulk eigenvalue matching analysis Ke et al. [2021]. Fitting a large number of factors is possible: in our experiments we fit a set of 197 factors.

The second approach involves three steps. In the first step we estimate a null distribution of eigenvalues based on sampling variances 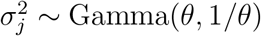 and then subsequently an *n* × *p* Gaussian matrix 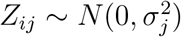. We sample *B* of these matrices and take the average of the sorted eigenvalues of 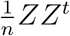 over *B* samples. Typically *B* is set to 100. Given the average sorted eigenvalues, we compute a regression coefficient without intercept between the “bulk” of these eigenvalues and the bulk of the observed eigenvalues of the data covariance matrix. The bulk of the distribution are the values between some lower and upper quantiles, which are hyperparameters of the method. We perform a line search on *θ* to find a value of *θ* that minimizes the sum of squared residuals of this regression. Denoting this regression coefficient as *β*, in the second step we simulate a background distribution based on sampling variance terms 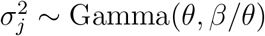, data from 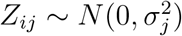, and eigenvalues from 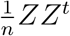. Finally, *K* is estimated as the (1 − *α*) quantile of the simulated distribution of leading eigenvalues. We apply this process for every cell type separately.

#### 2.7 Determining Spectra's input parameters

In Table 1, we summarize all user defined inputs to the Spectra algorithm. The data matrix and regularization strength *λ* must be provided by the user, while prior information can be provided in the form of a dictionary of cell type specific and global gene sets (note that Spectra can also be run by providing graph adjacency matrices directly). Optionally, cell type labels that align with keys of the gene set dictionary can be provided. The lower bound for gene scale factors, *δ*, controls the extent that gene expression is normalized and is set to a default value of 0.001. This translates to a maximum ratio of gene scale factors of ≈ 1000. By default the graph edge weights are set to be inversely proportional to the total number of edges induced by the gene set leading to a given edge, and accumulate additively for genes in multiple sets. The background rate of noise edges, *κ*, and the rate at which edges are randomly removed from the graph, *ρ*,can be provided as fixed parameters that provide users with an extra degree of control over the extent that the graph is modified. If they are set to None (default), they are estimated during the training process in the same manner that other model parameters are estimated.

**Table 1:** Spectra model inputs

| Input | Description |
| --- | --- |
| $\mathbf{X}$ | Expression matrix with $n$ cells and $p$ genes (Required). |
| $\lambda$ | Regularization strength of prior graph (Required). |
| Gene set dictionary | Dictionary with cell types as keys, gene sets as values (Optional). |
| Cell type labels | List of cell types corresponding to expression matrix (Optional). |
| $\delta$ | Parameter that bounds minimum gene scale factor (Optional). |
| $\mathbf{w}$ | Graph edge weights (Optional). |
| $\kappa$ | Background rate of edges (Optional). |
| $\rho$ | Background rate of edge deletion (Optional). |

For typically sized scRNAseq datasets, as a rule of thumb we recommend *λ* = 0.01 for studies in which factors should closely resemble the input gene sets and *λ* = 0.1 for studies where the factors should be allowed to deviate significantly from the gene sets. Values of *δ* ranging from 0.0001 to 0.01 yield similar results with *δ* > 0.01 providing solutions with typical highly expressed genes observed from NMF.

### 3 Validation metrics

#### 3.1 Marker list coherence metrics

To evaluate the quality of factors computed from data, we follow previous work (Stevens et al. [2012], González-Blas et al. [2022]) we use *coherence*, co-occurrence of factor genes in held-out data, to evaluate the quality of the inferred factors. For a given factor, we consider the 50 top marker genes with the highest gene scores for that factor. Between every pair of genes in the top 50 markers, we compute the *pointwise mutual information* as:

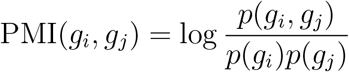

where probabilities denote the empirical occurrences in the held out data. Coherence is defined as the average of this quantity across the marker gene list. This metric is used in Fig. 3C. In order to assess the coherence of Spectra and other methods, we allocated 9787 cells as a hold out set to compute the coherence scores at evaluation time. The remaining 88, 076 cells were used to fit the model. For each experiment, we subsampled the 88, 076 cells in the training set to a size of 10, 000 without replacement (repeating this process 5 times to recapitulate the underlying data distribution). This number was chosen to be sufficiently large subject to the constraint that each of the methods under evaluation could run in a reasonable amount of time (< 2 days). For each subsampled dataset we computed the coherence score described above with the top 50 markers where marker lists are determined via the method suggested by the individual papers. For scHPF we used the `gene_scores` function from the scHPF package to get the top 50 markers Levitin et al. [2019]. For slalom, we multiplied the estimated parameter matrices, i.e. the continuous posterior mean 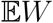 and Bernoulli posterior mean 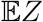, as in Buettner et al. [2017]. To evaluate non-negative matrix factorization, we derive marker lists based on the absolute values of the estimated factor matrix as is standard practice Barkley et al. [2022].

#### 3.2 Reconstruction of held out genes

To quantify the ability of methods to impute missing genes from gene sets, we ran Spectra and slalom (Section 6) on the full Bassez dataset but with randomly truncated gene sets. Due to slalom’s computational demands and size of the dataset, we choose a small set of 24 gene sets to evaluate for both methods, which are chosen a priori and held fixed throughout the experiment. We hold out 40 percent of genes (selected randomly) from the original set and measure the fraction of these genes recovered in the top 200 genes according to slalom and Spectra’s gene scores. In order to match factors to gene sets, for both methods we find the gene set (in our full database) with highest Szymkiewicz—Simpson overlap coefficient (overlap coefficient) to the given factor and label the factor as corresponding to that gene set. The overlap coefficient for the sets X and Y is defined as the size of the intersection divided by the size of the smaller set:

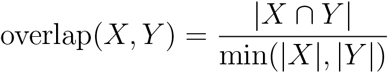

If two factors both have highest overlap coefficient to the same gene set, we take the one with higher overlap coefficient. The accuracy reported is the fraction of held out markers recovered in the top 200 highest gene scores (See Fig. 3A,B and S4).

### 4 Simulation Experiments

#### 4.1 Robustness to correlated factors

Matrix factorization methods rely on reconstruction based objective functions that implicitly encourage the estimation of a diverse set of gene programs. As a result, when gene programs are expressed in similar contexts (e.g. CD8 T cell activation, exhaustion, and tumor reactivity or TNF and IFN type II responses) matrix factorization methods often return a single program representing the combined set of correlated programs. Further, as the correlation between gene programs increases, the effective sample size of the estimation problem decreases, as most cells do not provide information to separate the gene programs. To illustrate that Spectra can incorporate prior information to maintain robust estimation in the presence of highly correlated gene programs, we simulated gene expression data from a generic factor analysis model where the cell loadings corresponding to factors 1 and 2 are simulated from a joint log Normal distribution with non-zero correlation terms ranging from 0.25 to 0.99 (Supplementary Fig. S5A). Factors themselves were simulated from a half-Cauchy distribution to achieve realistic levels of sparsity and extreme values. Conditional on simulated factors and loadings, gene expression was simulated from Poisson distribution with mean given by the matrix product of loadings and factors. A noisy prior knowledge graph was simulated by sampling the adjacency matrix from a Bernoulli distribution with parameters given by inner products between factors (as in the Spectra model) and used as input to Spectra. For each value of the correlation, we simulated 10 datasets and ran Spectra (lambda = 0.1), NMF, scHPF and Slalom (20 top genes per factor as input). We quantified estimation accuracy by the mean Pearson correlation of ground truth factors with estimated factors across genes, both for the two correlated factors and for a third factor uncorrelated with the first two. While the unbiased methods, NMF and scHPF, correctly recover the factors when factors are weakly correlated, estimation accuracy deteriorates as the correlation increases (though the inaccurate estimation of the correlated factors does not hurt performance on the uncorrelated factors). Spectra’s utilization of prior knowledge allows it to separate highly correlated factors.

In more detail, in our comparative simulation study factors are correlated in the sense that they tend to be expressed by the same cells (Supplementary Fig. S5A). We simulate ground truth factor matrices with *p* features and *K* factors with each entry independently distributed according to a half-Cauchy distribution (chosen to obtain realistically sparse factor matrices). In order to obtain correlated factors, the factor loadings, *α*, are independently drawn from a correlated LogNormal distribution:

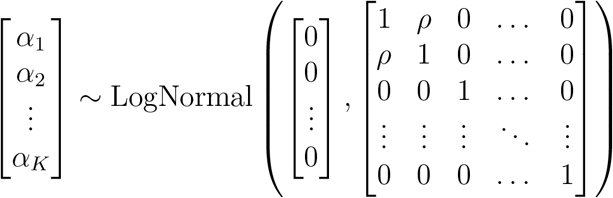

If we denote the *N* × *K* loading matrix by *α* and the *K* × *p* factor matrix by *θ*, the count data simulated by *X* ~ Pois(*αθ* + *ϵ*) where *ϵ* is a random noise term with variance *σ*^2^. An adjacency matrix is sampled coordinate-wise 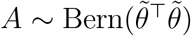 where 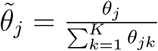. We run 10 independent trials, for 7 different levels of correlation *ρ* = {0.25, 0.5, 0.7, 0.85, 0.9, 0.95, 0.99}, totaling 70 simulated datasets. Since each of NMF, scHPF, slalom and Spectra estimates a factor matrix, we compared the estimated (normalized) factor matrices to 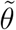 via Pearson correlation (*y*-axis of Supplementary Fig. S5A) after resolving the permutation of estimated factors that is closest to ground truth. Resolving the correct permutation for each estimate is done via finding the permutation that maximizes the average Spearman correlation between ground truth factors and estimates. Since we are interested in performance on correlated factors, we report the average correlation between estimation and ground truth for the first two correlated factors, across the 10 independent trials.

In our experiment, *N* = 20, *p* = 500, *K* = 3, *σ*^2^ = 4 (a setting with low signal to noise ratio). Spectra is provided with the simulated matrix *A* while slalom is provided with feature sets containing the correct top 20 features of each factor. Spectra uses a *λ* value of 0.1 and *δ* value of 0. All methods are run with the correctly specified number of factors and with default parameters.

#### 4.2 Biasedness of gene set averaging for overlapping gene sets

When gene sets corresponding to gene programs overlap, simple gene set averaging approaches produce false positive program activity calls. To illustrate this phenomenon, we simulated gene expression data driven by sets of overlapping gene programs with varying degrees of gene set overlap and showed that gene set averages are increasingly biased proxies for program activity as the degree of gene set overlap increases (Supplementary Fig. S5B). Specifically, we simulated factor matrices with known sparsity pattern determined by a set of gene sets (each non-zero entry independently Exponential(16)). Each gene set is designed to have overlap coefficient *ρ* with at least one other gene set, with *ρ* ranging from 0 to 0.75. Loadings are generated by first sampling each coordinate LogNormal(0, 1) independently, and then zeroing out components that are < 1 to induce sparsity. Simulated expression data is from a Poisson distribution with mean given by the matrix product of simulated loadings and factors.

For each possible value of the overlap coefficient *ρ* in {0.0, 0.3, 0.5, 0.75}, we create 3 simulated datasets and ran both Spectra and `score_genes` with the ground truth gene sets. Accuracy is measured by the Pearson correlation of estimated cell scores (or `score_genes` estimates) and the ground truth factor loadings from the data generation process (*y*-axis of Supplementary Fig. 5B). In this experiment, the gene set size is fixed to 20, the number of gene sets is 10, the number of features (genes) is 500, and the number of observations is 1000.

#### 4.3 Recovery of active gene sets

We compared Spectra to slalom, another factor analysis method that uses prior information in the form of gene sets, in a simulation experiment where we measured the ability of each method to recover the gene sets involved in the true data generating process. Here, we followed the simulation settings of the Buettner et al. [2017] closely. First background factors are generated from an Exponential distribution, as before. To simulate sparsity entries smaller than 2 are zeroed out. Next, loadings are generated LogNormal independently and entries < 1 are zeroed out. We then generate both active and control gene set based factors as in Buettner et al. [2017] and Section 4.2 where gene sets overlap with overlap coefficient 0.3. Loadings corresponding to active gene set based factors are also drawn from a standard LogNormal and zeroed out if less than 1. Next genes are randomly added to gene sets and removed from genesets to achieve a False Positive Rate (FPR) of 0.2 and False Negative Rate (FNR) of 0.2. As a measure of success we use the area under the ROC curve (AUC) based on slalom’s relevance score and Spectra’s average cell score for a given factor. Spectra was robust to increasing the number of gene sets while slalom suffered a drop in AUC as the number of active gene sets increased (Supplementary Fig. S5C).

In our experiments, the number of active pathways vary on the *x* axis of Supplementary Fig. S5C, the number of control pathways is 5, the gene set size is 20, the number of genes is 300, the number of cells is 300, the number of unbiased factors is 5, the gene set overlap coefficient is 0.3.

### 5 A human immunology knowledge base

Databases such as the Gene Ontology Resource (GEO) The Gene Ontology Consortium [2018], the Molecular Signatures Database (MSigDB) Liberzon et al. [2011], the Kyoto Encyclopedia of Genes and Genomes Kanehisa et al. [2016] and the Reactome database Croft et al. [2010] contain thousands of gene sets and their relationships, but they are noisy and often do not distinguish whether or not genes are transcriptionally regulated. For example, many genes with signaling pathway annotations are regulated at the post-translational level by phosphorylation or subcellular localization. Expression signatures in these databases are often derived from bulk sequencing data which may not represent responses in individual cells. Moreover, the databases do no have a framework for distinguishing which gene sets are cell-type specific. To address these issues, we created an immunology knowledge base with the following criteria:

1. Genes within a gene sets define a cellular process at the transcript level
2. Gene sets represent cellular processes at the single cell level
3. Gene sets can be specific to a defined cell type

Our knowledge base includes 181 gene sets representing ‘cellular processes’ to be queried by Spectra. Like Spectra, the knowledge base models gene sets as a graph wherein every gene set is a node connected to all individual gene nodes within the set, as well as to a cell type node. Cell type nodes (currently 50) are connected to ‘cellular identity’ gene sets, one for each cell type, which contain marker genes for their connected cell type. Metadata such as scientific publication the gene set was derived from, gene set version, and original gene set authors are stored as node properties. Cell type nodes are organized in a hierarchy, reflecting that cell types are frequently subsets of other cell types. This hierarchy starts with a cell type node labeled ‘all-cells’ to which gene sets for ‘cellular processes’ occurring in all cell types are connected. Thus, the knowledge base can be queried for a cellular processes that can be found in all cell types (e.g. glycolysis) or a cellular process specific to a cell type such as T cell receptor signaling which is only present in T cells. It also allows retrieving ‘cellular identity’ marker gene sets which define the queried cell type.

Within this resource, 150 cellular processes apply to all leukocytes and 31 apply to individual cell types. Of all 231 gene sets, 97 were gene sets newly curated from the literature, 14 used data from perturbation experiments, 11 were adopted from literature with modifications and 123 were taken from the literature and external databases without changes. Gene sets correspond to diverse cellular identities (*n* = 50) and cellular processes such as homeostasis (*n* = 9), stress response (*n* = 3), cell death and autophagy (*n* = 18), proliferation (*n* = 6), signaling (*n* = 12), metabolism (*n* = 90), immune function (*n* = 22), immune cell responses to external stimuli (*n* = 18) and hemostasis/coagulation (*n* = 3) (Fig. 2b). We designed the gene sets for cellular processes to have comparable size (median *n* = 20 genes per gene sets) and relatively little overlap (median pairwise overlap coefficient 40%) to enable dissection of a large number of cellular processes and avoid gene set size-driven effects.

To specify Spectra input, the user first defines cell types at a granularity of interest in their single-cell expression data, and retrieves the cell-type-specific cellular process gene sets and gene sets applying to all cell types from the knowledge base. Next, they can select cellular process gene sets pertaining to all cell types in the dataset, which should be set as ‘global’ in the Spectra model.

The user indicates which cellular processes can be considered global based on which cell types are present in the dataset under study. For example, if a dataset only contains T cells, all cellular processes pertaining to leukocytes and T cells should be considered global. If cellular processes apply to more than one but not all cell types in the data there are two options:

1. The gene set can be multiplied and one copy can be assigned to each cell type. This will ensure that the cell scores of the resulting factors will be specific to those cell types but may result in an separate factors for the same cellular process in each cell type.
2. The gene set can be set as global which will generally result in one factor. However, cell scores for this factor may be detected in other cell types also.

Users can take advantage of the hierarchical organization of cell types in the knowledge base by adding the children or parent classes of selected cell types. For example, cellular processes for both ‘CD4 T cells’ and its parent, ‘T cells’ can be retrieved and assigned to ‘CD4 T cells’, making it possible to find broader processes (e.g. T-cell receptor signaling) that are specific to CD4 T cells. Alternatively, cellular processes for CD4 T cells and for CD4 subtypes ‘TH1’, ‘TH2’ and ‘TH17’ can be retrieved and assigned to ‘CD4 T cells’, thereby pooling rare cell types which may not contain enough training data for the Spectra model to converge to a generalizable solution. Moreover, hierarchical classification is advantageous when cellular processes are ambiguously or incorrectly assigned. For example, CD4 T cell subtypes are often presented as distinct lineages with distinct cellular processes. However, mixed CD4 T cell subtypes have been reported, such as cells possessing both TH1 and TH2 polarization cellular processes Eizenberg-Magar et al. [2017], suggesting that CD4 T cells can be described using combinations of purportedly subtype-specific processes.

Our human immunology knowledge base is available on GitHub as the Cytopus python package https://github.com/wallet-maker/cytopus Walle [2022]. Users can load our default or a custom knowledge base using the `KnowledgeBase` class build on a NetworkX object. Cytopus includes methods to retrieve gene sets and corresponding cell types, visualize them as a graph and convert them into a Spectra-compatible dictionary. The `celltypes` method retrieves a list of available cell types; `processes` generates a dictionary of all ‘cellular processes’ gene sets; and `identities` generates a dictionary of all ‘cellular identity’ gene sets. Gene set metadata (e.g. author, topic, date of generation, version) can be accessed as node properties of the gene sets. The `get_celltype_processes` method retrieves cell-type-specific ‘cellular processes’ based on a user-provided list of cell types at the desired granularity (generally all cell types contained in the data).

Full cytopus documentation can be found at https://github.com/walletmaker/cytopus/. The tumor infiltrating leukocyte gene sets used in the paper are included in the Spectra package (https://github.com/dpeerlab/Spectra) as well as in Supplementary Table 1.

### 6 Single Cell RNAseq Data Preprocessing and Analysis

#### 6.1 Immunooncology datasets

To study Spectra in an immuno-oncology context we used two published scRNA-seq datasets of tumor-infiltrating leukocytes from female breast cancer patients treated with immunotherapy. We chose this immuno-oncology context for multiple reasons:

1. The abundant prior knowledge of cellular processes and well-characterized cell types in tumor infiltrating leukocytes enabled us to leverage the full power of gene set and cell type priors.
2. The availability of before- and on-treatment samples to test the sensitivity of factor cell scores to environmental perturbation with anti-PD-1/PD-L1 therapy
3. The clinical need for detecting cellular processes affected by anti-PD-1 in humans to improve current immunotherapy strategies.
4. The availability of two studies in similar biological settings to enable validating findings in an independent dataset.

##### 6.1.1 Bassez dataset

Bassez et al. [2021] was a prospective window-of-opportunity study reporting scRNAseq as an exploratory endpoint (’Bassez dataset’). The authors analyzed scRNAseq data from whole tumor single-cell suspensions from 42 operable breast cancer patients before and after anti-PD-1 immunotherapy (Pembrolizumab, NCT03197389). Patients received neoadjuvant chemotherapy (CTX) as per standard of care (CTX n=11, no CTX n=31) followed by a single dose of anti-PD-1. Breast resections were performed 7−14 days after anti-PD-1. Tissue from pre-anti-PD-1 biopsies (7-15 days before surgery) and from surgical resections was processed for scRNAseq. As a surrogate for response to therapy the authors of the original study quantified the clonal expansion of T-cells under therapy on the patient level using paired single cell T-cell receptor sequencing (scTCR-seq); we used these annotations to find response-associated cellular processes in the data. The authors categorized patients as either exhibiting (responders) or lacking (non-responders) T-cell clonal expansion under therapy. To classify patients, they quantified the number of expanding T cell clones (T-cells with identical T-cell receptor (TCR) sequences) per patient, labeling patients with > 30 expanding clones as responders and with ≤ 30 expanding clones as non-responders. A T-cell clone had to fulfill two criteria to be labeled as expanded:

1. Detected at least twice in the patient’s on-treatment sample
2. More frequent in the patient’s on-treatment as compared to the pre-treatment sample either by the absolute cell number in that clone or by the cell number in that clone relative to the number of cells with a TCR detected.

##### 6.1.2 Zhang dataset

Zhang et al. [2021] was a retrospective clinical study analyzing tumor-infiltrating leukocyte scRNAseq data of pre-on- and post-therapy samples from 22 advanced female breast cancer patients receiving either anti-PD-L1 (atezolizumab) combined with chemotherapy (paclitaxel, n=11) or chemotherapy alone (paclitaxel, n=11) (’Zhang dataset’). Notably, patients received corticosteroid pre-medication for paclitaxel. The authors assessed patient response to immunotherapy using radiological response according to RECIST v1.1 criteria Eisenhauer et al. [2009]. RECIST v1.1 are the standard criteria used for drug-approval relevant clinical trials and standard clinical management of cancer patients. The RECIST criteria classify patients into responders (combining partial and complete response labels) and non-responders (combining progressive and stable disease labels) based on the change in the sum of tumor lesion diameters under therapy. We used this classification to identify response-associated cellular processes in the Zhang dataset.

#### 6.2 Processing strategy

To minimize systematic differences in cell type annotations and normalization of gene expression counts, we performed the same pre-processing for the Bassez et al. [2021] and Zhang et al. [2021] data. After basic filtering, removing residual low-quality cells and doublets and subsetting to leukocytes with scanpy Wolf et al. [2018], we normalized the data using scran (L Lun et al. [2016]). We hierarchically annotated celltypes in the data first labeling major immune subsets (T/ILC cells, B/plasma cells, myeloid cells) by clustering on the most dominant principal components (PCs) only. We then partioned the data into these major immune subsets, re-normalized the data within every subset using scran, and clustered on more PCs to annotate granular cell types. We then combined the annotated data from major immune subsets for joint analysis using Spectra. We have outlined the details of the analysis strategy below.

**Table 2:** Clinical variables of the utilized scRNAseq datasets

|  | operable | cohort 1 (n) | cohort 2 (n) | cortison |
| --- | --- | --- | --- | --- |
| Bassez | Yes | anti-PD-1 (n=31) | CTX, anti-PD-1 (n=11) | - |
| Zhang | No | anti-PD-L1 + CTX (n=11) | CTX (n=11) | + |

##### 6.2.1 Retrieving single cell gene expression data

Count matrices of the Bassez et al. [2021] study were kindly provided by the authors and are also available at (http://biokey.lambrechtslab.org, 226, 635 cells). Raw read counts are available in the European Genome-phenome Archive (EGA, https://ega-archive.org/) under accession numbers: EGAS00001004809, EGAD00001006608. The count matrices for the Zhang et al. [2021] study were downloaded from the Gene Expression Omnibus (GEO, https://www.ncbi.nlm.nih.gov/geo/) using the following accession number: GSE169246 (489, 490 cells).

##### 6.2.2 Removing low quality cells

To prepare the data for clustering, we removed cells with less than 200 genes per cell and genes observed in less than 20 cells, as well as mitochondrial and ribosomal genes. This filtering procedure removed 2971 and 203 genes resulting in a total of 22, 639 and 20, 898 genes in the Bassez et al. [2021] and Zhang et al. [2021] data, respectively. We defined doublets in the data by running DoubletDetection Gayoso et al. [2020] for each sample individually using standard parameters (clustering algorithm: PhenoGraph, p-value threshold: 1*e* 16, voter threshold: 0.5). DoubletDetection detected 3270 (1.4%) and 12, 760 (2.6%) doublets as well as 27 (0.01%) and 8 (0.001%) ambiguous doublets in the Bassez et al. [2021] and Zhang et al. [2021] data respectively which we removed from the data.

##### 6.2.3 Retrieving tumor-infiltrating leukocytes for downstream annotation

While the Zhang et al. [2021] data contained sorted tumor infiltrating leukocytes the Bassez et al. [2021] data contained unsorted whole tumor single cell suspensions. To retrieve immune cells from the Bassez et al. [2021] data for downstream annotation, we first performed standard median library size normalization and log1p-transformed the data so that the normalized expression of every gene *j* in cell *i* is 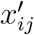 and the median gene expression in cell *i* is *med*(*x_i_*):

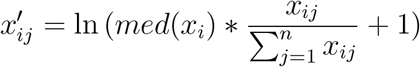

We then clustered the data using PhenoGraph Levine et al. [2015] on the most dominant principal components which we selected using the knee point of the PC vs. explained variance curve (calculated using the kneed package v.0.7.0 Arvai [2020]) or the lowest number of PCs explaining ≥ 20% of the total variance whichever was higher. Using this procedure we clustered the data with PhenoGraph on the first 26 PCs explaining 20.1% of total variance. We chose k=80 parameter for PhenoGraph because of its ability to delineate immune cell from non-immune cell populations while showing stable clustering in a window of adjacent k parameters (pairwise rand indices > 0.7). We then subsetted leukocytes for further analysis by their marker gene expression (myeloid cells, T cells, innate lymphoid cells = ILC, B cells and plasma cells) per PhenoGraph cluster (Supplementary Table 2).

##### 6.2.4 Annotating tumor-infiltrating leukocytes

We re-normalized leukocytes in the Bassez et al. [2021] and Zhang et al. [2021] data using scran because median library size normalization can generate artificial differential gene expression between cells of different library size such as leukocytes L Lun et al. [2016]. After testing all genes and a range between 5, 000 and 15, 000 HVG, we selected the top 15, 000 highly variable genes (HVG) for the Bassez et al. [2021] and all genes for the Zhang et al. [2021] data which led to the best separation of major immune cell subtypes using scanpy’s `pp.highly_variable_genes` function with the `seurat_v3` method on raw counts. To avoid discarding genes relevant to cell typing we added a manually curated list of 459 cell typing markers to HVGs (Supplementary Table 5). We then repeated the clustering procedure outlined above (Bassez et al. [2021]: 24 PCs explaining 20.1% of total variance Zhang et al. [2021]: 52 PCs explaining 20% of total variance, k=50) and annotated major immune cell subsets (T/ILC cells, B/plasma cells, myeloid cells) by assessing their mean marker gene expression per cluster (Supplementary Table 2). To obtain more granular annotations, we partitioned the data into major immune subtypes (T, ILC, B/plasma, myeloid), we re-normalized each subtype using scran, re-calculated HVGs and PCs and clustered as described above. The processing parameters for each subtype are indicated in Table 3 below:

We then annotated granular immune cell types by assessing the mean marker gene expression per cluster (Supplementary Table 2). In the Bassez et al. [2021] data, we detected clusters with low library size and lower complexity of gene-gene correlation patterns at this step (5, 509 cells) which we removed from the data. Finally, we combined the annotated major immune subtypes for downstream joint analysis.

**Table 3:** Clustering parameters for immune subtypes

|  | Bassez data |  |  | Zhang data |  |  |  |
| --- | --- | --- | --- | --- | --- | --- | --- |
|  | TNK | B | M | TNK | ILC | B | M |
| HVGs | 7,500 | 7,500 | 15,000 | 19,379 | 19,379 | 10,000 | 18,888 |
| PCs | 24 | 10 | 16 | 100 | 100 | 17 | 23 |
| variance[%] | 20.3 | 20.1 | 20.1 | 17.9 | 17.7 | 20.4 | 20.1 |
| k | 30 | 40 | 20 | 40 | 40 | 60 | 30 |

#### 6.3 Running Spectra

After the filtering steps above, the Bassez et al. [2021] and Zhang et al. [2021] data had 97, 863 and 150, 985 cells respectively. To run Spectra, we restrict the number of genes using scanpy’s `highly_variable_genes` function with the `cell_ranger` method selecting the 3, 000 most highly variable genes. We removed several genes which are highly abundant and may originate from ambient RNA in many cell types thus adding noise to the analysis. This included genes for immunoglobulins (genes starting with *IGHM*, *IGLC*, *IGHG*, *IGHA*, *IGHV*, *IGLV*, *IGKV*), T cell receptor variable domains (genes starting with *TRBV*, *TRAV*, *TRGV*, *TRDV*) and hemoglobins (genes starting with *HB*). For the Bassez et al. [2021] and Zhang et al. [2021] datasets, the total number of genes used (the union of genes included in a gene set and highly variable genes) were 6397 and 6398 respectively.

Gene sets were then converted into weighted adjacency matrices. One of Spectra’s strongest features is its ability to meaningfully modify the input gene-gene knowledge graph (gene sets) in a data driven matter: With the influence parameter *λ* set to 0.01, the median overlap coefficient across all factors in the Bassez et al. [2021] data was 88%, with 25% of factors relevantly deviating from the gene sets (overlap < 70%), and 7% of factors bearing little resemblance to the input gene sets (overlap < 20%). With the influence parameter set to 0.1, the median overlap coefficient across all factors was 82% with 42% of factors relevantly deviating from the gene sets (overlap < 70%) and 12% of factors with overlap less than 20%. In terms of graph edit distance to the input graph (defined as the mean absolute difference between input and output graphs), at *λ* set to 0.1, we had 0.011 and at *λ* set to 0.01 we had 0.0095 with diminishing returns in graph edit distance for lower *λ* (0.0095 again for *λ* = 1*e* − 4 and 0.0094 for *λ* = 1*e* − 5). For the analyses described below, we used an influence parameter *λ* between 0.1 and 0.01 depending on whether we wanted more (0.01) or less (0.1) adherence to the input gene sets. Because we obtained very similar results with these parameters in two independent datasets, it is likely that this also constitutes a good default for other datasets.

#### 6.4 Running Slalom

Because Slalom’s runtime scales linearly with the number of gene sets (Fig. 3d), we had to subset the number of gene sets used in order to run Slalom on our datasets of interest (*n* = 20 gene sets, runtime 63.49 CPU hours, 40GB memory on the Bassez et al. [2021] dataset). Expression data was preprocessed identically to Spectra. To compare results of Spectra and Slalom we chose a subset of 20 gene sets of scientific relevance to the immune microenvironment under immune checkpoint blockade: CD8 T cell tumor reactivity, Type II interferon response, Myeloid angiogenic effectors, Post-translational modification, MHC class I presentation, G2M transition, Oxidative phosphorylation, Type I interferon response, Macrophage IL4/IL13 response, Glycolysis, DNA synthesis, G1S transition, Lysine metabolism, MHC class II presentation, Hypoxia response, Pentose phosphate pathway, CD8 terminal exhaustion, PD-1 signaling, TCR activiation and cytoxicity effectors. These gene sets were used to determine the I parameter of the slalom `initFA()` function. The following additional input parameters were used: `nHidden=0, nHiddenSparse=0, do_preTrain=False, minGenes = 1 pruneGenes = False` with all other options set to default values.

#### 6.5 Running scHPF

scHPF was run with the `run_trials()` function from the schpf pypi package. Preprocessing was identical to all other methods tested. Specifically, the following setting is used: nfactors=100, ntrials=1, min_iter=20, max_iter=1000, check_freq=10, epsilon=0.001, better_than_n_ago=5, dtype=np.float32,verbose=True, return_all=False,reproject=True, batchsize=0, beta_theta_simultaneous=True, loss_smoothing=1.

#### 6.6 Assigning factor labels

Factor labels were assigned using the overlap coefficient of the top 50 marker genes (genes with the highest gene scores) with each gene set. We observed a bimodal distribution of overlap coefficients with one group of factors centered close to 0 and one group of factors centered close to 1 (Supplementary Fig. 2). We therefore chose a threshold of 0.2 to separate high overlap from low overlap factors. For every factor, if the maximum overlap coefficient was > 0.2 we assigned the gene set label with the maximal overlap coefficient to that factor, if the maximum overlap coefficient was ≤ 0.2 we did not assign a label to that factor.

#### 6.7 Classifying new and modified factors

We classified all factors as new, modified or unspecified based on their input gene-gene knowledge graph dependency parameter *η*. The dependence parameter is a scalar value between zero and one, that quantifies its reliance on the input gene set graph. We observed a bimodal distribution of *η* with one group of factors centered close to 0 and another group of factors centered close to 1 (Supplementary Fig. 2a). We therefore chose a threshold of 0.25 to separate high dependence from low dependence parameters. We defined new factors as factors with a graph dependency parameter *η* < 0.25 and modified factors as factors with a graph dependency parameter *η* ≥ 0.25.

#### 6.8 Analyzing tumor infiltrating leukocytes

We compared Spectra, Slalom and scHPF’s capacity to retrieve features of tumor infiltrating immune cells. We ran the three algorithms on all leukocytes in the Bassez dataset as described above using a *λ* parameter of 0.01 for Spectra. We also ran Spectra on the Zhang dataset using a *λ* parameter of 0.01. For cells with high library size, such as macrophages (Bassez dataset median library size = 8038), we calculated gene scores for Spectra factors using an `offset` of 1 which retrieved more stably expressed genes (e.g. mean scran normalized expression of top 50 marker genes of the macrophage factor 182: 1.15 with offset vs 0.41 with no offset). For remaining analyses with lower library size, such as T-cells (Bassez dataset median library size = 3127) or B cells (Bassez dataset median library size = 3954), we calculated gene scores for Spectra using an `offset` of 0 which allowed for more sensitive retrieval of lowly-expressed genes such as transcription factors involved in tumor reactivity and exhaustion (e.g. *EOMES, TOX*), as well as metabolic processes (e.g. *PIPOX, BBOX1*).

##### 6.8.1 Visualizing scRNAseq data

To visualize individual genes in embeddings and account for sparsity in scRNA-seq data, we imputed gene expression using scanpy’s implementation of MAGIC Setty et al. [2019] with a t parameter of 3 and the exact solver (Supplementary Fig. 3a, 6c,7a, 8b, 8c). For visualizing all leukocytes, we calculated t-SNE embeddings on 57 PCs explaining 25.0% (Bassez dataset) or 55 PCs explaining 20% of variance (Zhang dataset) with standard parameters including a learning rate of 1000 using the scanpy implementation (Fig. 2c, 2e, 5d; Supplementary Fig. 3a, 7d, 8a).

##### 6.8.2 Aggregating factor cell scores at the patient level

Spectra factors are sparse and bimodally distributed, with one mode centered around zero and one more positive mode. To aggregate factor cell scores at the sample level we calculated the average cell score of the positive fraction (Fig. 4f, Supplementary Fig. 3b, 6g). The positive fraction was defined as cells with cell score > 0.01 selected empirically as the threshold which best separated the two cell score modes. If all cells showed a cell score ≤ 0.01 for a given gene program the mean of the positive fraction was set to 0 for that gene program.

##### 6.8.3 Gene set enrichment analysis

To find the most representative factors for a gene set in the Spectra, Slalom and scHPF, we performed gene set enrichment analysis for the exhaustion and tumor reactivity input gene sets in the top 50 marker genes (genes with highest gene scores) of every factor using gseapy’s enrichr function Fang et al. [2022] (Supplementary Fig. 6d, 7b). The enrichr function calculates enrichement using a hypergeometric test to calculate the probability of drawing the observed number of genes belonging to a gene set of interest when sampling froma pool of all genes without replacement (here: the union of the 3000 most highly variable genes plus the genes contained in the gene sets, see 6.3 ‘Running Spectra’). We calculated enrichment of gene sets in the top 50 markers genes (genes with highest gene scores) for each factor:

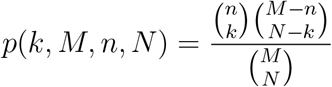

where *n* is the size of the gene set, *k* is the number of genes from the gene set in the top 50 marker genes in the factor, *M* is the total number of genes contained in the data, *N* is the number of draws from the all genes in the data (here: top 50 marker genes). From this, we calculated an FDR using the Benjamini-Hochberg correction. We assumed the factors with the lowest FDR for enrichment were representative for the respective gene sets if FDR < 0.05.

##### 6.8.4 CD8 T cell analysis

We took the subset of CD8 T cells (11 clusters) from Bassez dataset to explore CD8 T cell tumor reactivity (tumor reactivity) and CD8 T cell exhaustion (exhaustion). The most representative factors for the tumor reactivity and exhaustion gene sets were retrieved using the gene set enrichment procedure described above (6.8.3 ‘Gene set enrichment analysis’) for each factor analysis method (Supplementary Fig. 6d). Spectra factors were also compared to expression scores for the tumor reactivity and exhaustion gene sets using scanpy’s `score_genes` function (Fig. 4b, Supplementary Fig. 6c). To find genes driving `score_genes` expression we calculated the covariance of all genes within the tumor reactivity or exhaustion gene sets with the tumor reactivity and exhaustion gene scores (Supplementary Fig. 6b). To visualize of force-directed layouts we used scanpy’s `tl.draw_graph` function and the `ForceAtlas2` method on a nearest neighbors graph computed on CD8 T cells using scanpy’s `tl.neighbors` function with *n* = 10 nearest neighbors (Fig. 4a, 4b, 4c, Supplementary Fig. 6c). Contour plots were created using seaborn’s `jointplot` kernel density estimation with standard parameters (Fig. 4b, 4c, Supplementary Fig. 6f).

##### 6.8.5 Metabolism analysis

We assessed the expression pattern of metabolic factors across cell types and found highly specific expression of the lysine metabolism program in plasma cells. In the Bassez dataset we noticed a small (*n* = 114 cells, 0.1% of all cells, 3% of all plasma/B cells) group of heterotypic doublets expressing plasma cell and T cell markers (*CD3E*, *CD3D*, *CD3G*, *IGHG4*, *IGHG1*) which was not apparent in previous analyses and not detected by DoubletDetection. We removed these cells from further analyses involving plasma cells (Fig. 5). We also inspected the mean expression per cell type of the MAGIC Van Dijk et al. [2018] imputed (scanpy implementation, *t* = 3, exact solver) top 50 individual marker genes of the lysine factor genes with highest gene scores (Supplementary Fig. 7a)

##### 6.8.6 Macrophage analysis

To analyze differentiation gradients in macrophages and to capture all possible maturation stages, we retained the subset of 18 clusters (12, 132 cells) annotated as mature macrophages (12 clusters) or more immature myeloid derived (suppressor) cells/monocytic cells (6 clusters) in the Bassez dataset for further analysis (Supplementary Table 2). We embedded the data using diffusion components (DCs) which preserve differentiation trajectories better than many common linear and non-linear dimensionality reduction techniques Haghverdi et al. [2015]. Using a classification strategy, we selected the DCs which best captured the differentiation from more monocytic states to macrophages while separating patients with (responders) and without clonal T cell expansion (non-responders, Fig. 6a, 6b, Supplementary Fig. 8a, 8b). For every pair of the first 20 DCs we performed 1) a linear regression with the DCs as the independent variable and the scran-normalized expression of the monocyte marker *S100A8* as the dependent variable and 2) a logistic regression with the DCs as the independent variable and response status as the dependent variable. We chose the DC pair with highest sum of the coefficient of determination R^2^ (linear regression) and highest mean accuracy of (logistic regression). We calculated Spectra cell score trends over the DCs by fitting a generalized additive model as implemented in Palantir’s `_gam_fit_predict` and `calculate_gene_trends` functions Setty et al. [2019] using cell scores instead of gene expression and DCs instead of pseudotime (Fig. 6a):

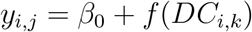

where *y_i,j,k_* is the cell score of cell *i*, factor *j* and the *k*th diffusion component and *DC_i,k_* the *k*th diffusion component for cell *i*. We then visualized cell score trends using the `plot_gene_trend_heatmap` function from Palantir.

To calculate compositional changes under immune checkpoint therapy we used the Milo package Dann et al. [2022]. Milo is analogous to differential gene expression analysis, but instead of identifying genes that differ between two groups of cells, it tests for differential cell density in (possibly overlapping) neighborhoods of a k-nearest neighbors (KNN) cell-cell similarity graph, across different conditions. We chose the default fraction of 0.1 to be sampled as index cells from the KNN graph, such that representative cellular neighborhoods were only constructed for those index cells. Milo counts the number of cells per sample in each neighborhood and uses a generalized linear model with a negative binomial distribution to test for differences in abundance. Milo also accounts for multiple comparison testing by computing a spatial false discovery rate (FDR).

For the Bassez dataset we constructed a KNN graph on macrophages and monocytic cells. The Milo paper gives the following heuristic to estimate an optimal *k* parameter Dann et al. [2022]:

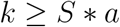

where *S* is the number of samples (here: 79) and *a* is an arbitrary scaling parameter. Following the authors’ suggestion of 3 ≤ *a* ≥ 5 resulted in an overly large *k* parameter 237 ≤ *k* ≥ 395. We therefore chose *k* to be smaller than the smallest population of cells identified by clustering (58 cells) but close to the *k* parameter obtained by the heuristic above resulting in *k* = 50 to construct the KNN graph and to identify the nearest 50 neighbors of the index cells.

We then assessed the fold change of cell states under PD-1 blockade using the following regression formula:

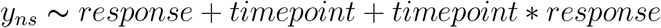

where *y* is the number of cells from sample *s* in neighborhood *n* and *response* status is defined as 0 for non-responders and 1 for responders. We defined the *timepoint* as 0 for pre-therapy and 1 for on-therapy. *timepoint response* indicates the interaction between the *timepoint* and *response* variables. We then identified the neighborhoods specifically enriched for non-responders under therapy by taking the subset of neighborhoods based on the estimated regression coefficients. First we identified the neighborhoods specifically enriched under therapy by retaining a subset all neighborhoods with an FDR < 0.05 and coefficient (log fold change) > 0 for the *timepoint* parameter for further analysis. From these, we took a subset of the neighborhoods enriched for non-responders as compared to responders under therapy by selecting neighborhoods with an interaction FDR < 0.05 and an interaction coefficient (log fold change) < 0 for further analysis. We then compared the mean factor cell scores for these neighborhoods with all remaining neighborhoods.

The Zhang dataset contained less patients than the Bassez dataset (*n* = 11 vs *n* = 42) and therefore did not allow for testing as many covariates. We thus chose a slightly different strategy to find macrophage neighborhoods enriched for non-responders under therapy. As for the Bassez dataset we took a subset of 16 clusters (11, 466 cells) annotated as mature macrophages (12 clusters, 9, 385 cells) or more immature myeloid derived (suppressor) cells/-monocytic cells (4 clusters, 2, 081 cells) (Supplementary Table 2), and then selected samples from patients classified as non-responders (see 6.1.2) for a total of 4, 318 cells and 15 samples. Analogously to the *k* parameter selection strategy above, we constructed a KNN graph using a *k* parameter of 20 which was smaller than the smallest cell population detected by clustering (22 cells). We then defined Milo neighborhoods as the 30 nearest neighbors of the index cells. We fitted the Milo model using the following regression formula:

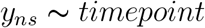

where *y* is the number of cells from sample *s* in neighborhood *n* and the timepoint defined as either 0 for pre-therapy or 1 for on-therapy. We took a subset of the neighborhoods with *FDR* < 0.2 and a coefficient (log fold change) > 0 for the *timepoint* parameter for further analysis. As for the Bassez dataset, we then compared the factor cell scores for this group with all remaining neighborhoods.

#### 6.9 Statistical analysis and visualization

P values were calculated as indicated above using the Milo, scipy and statsmodels python packages. No normality assumption was made. We used a Mann-Whitney U test for independent samples and a Wilcoxon matched pairs signed rank test for paired samples. If not indicated differently, all p values are two-sided and a multiple-comparisons corrected (Bejamini-Hochberg method) p value of 0.05 was considered statistically significant. Data was visualized using the matplotlib and seaborn python packages or GraphPad Prism (v9.4.1) for Windows 10 and edited in Adobe Illustrator Creative Cloud (v27.0).

### 7 Theoretical Analysis

#### 7.1 Spectra behaves like gene set averaging when *λ* → ∞

A standard real analysis argument shows this formally. For ease of exposition we assume that the gene scalings *g* are 1 and *ρ* = *κ* = 0. Simulations confirm the result. Recall that *B_kl_* ∈ [0, 1] and *θ_i_* ∈ Δ^*K−*1^.

##### Theorem 1

*Suppose that 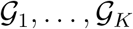 are a partition of* {1,…, *p*} *and A is constructed such that A_ij_* = 1 *if 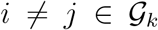 for some k. Define 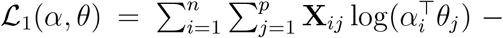 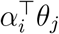 and 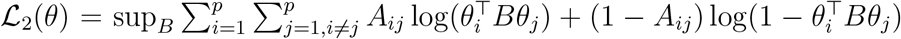. We have 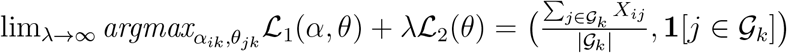 (or some permutation)*

##### Lemma 1

*If each block is at least size* 2, 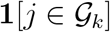 *is a unique solution up to permutations σ*(*k*) *for the loss* 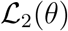. *Since θ is constrained to a compact set and* 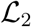 *is continuous, if* 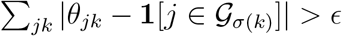 *we must have a δ* > 0 *with* 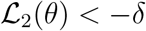

Proof of lemma. From the strict convexity of the Bernoulli log likelihood we have that the maximum must satisfy 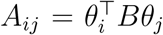 for *i* ≠ *j*. Define the following subsets of {1,…, *K*} : 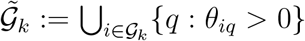. Note that each 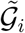 is nonempty. If one were empty we would have *θ_i_* = 0 for all 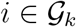 which is impossible if each block has size at least 2.

Consider 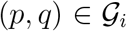 with *p* ≠ *q*. By assumption 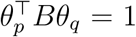. Thus it must be the case that we have *B_kl_* = 1 for some *k, l* (possibly *k* = *l*) and *θ_pk_* > 0, *θ_ql_* > 0. Further 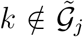 for all *j* ≠ *i* and 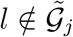 for all *j* ≠ *i*. To see this, we have by assumption, *θ_pk_* > 0 and 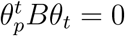 for all 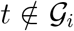. If some such *t* had *θ_tl_* > 0 then *B_kl_θ_pk_θ_tl_* = 0 a contradiction. This shows that *k* and *l* are both unique to 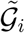. But this applies to each *i* = 1,…, *K*. So *k* = *l* and 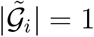 for all *i*. Then we have *B_kk_θ_pk_θ_qk_* = 1 for a unique *k* and *B_kk_* = *θ_pk_* = *θ_qk_* = 1.

Remark that 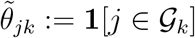 is a global minimizer of 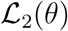. An upper bound of 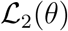 is:

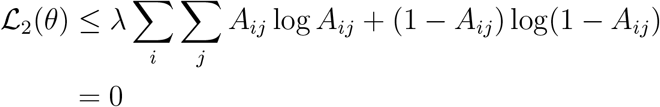

while setting *B* = *I* gives 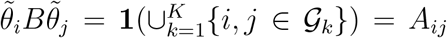 matching the upper bound. 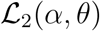 is convex in either argument and the solution to 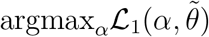 is given by 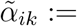 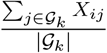, which is clear by the computation:

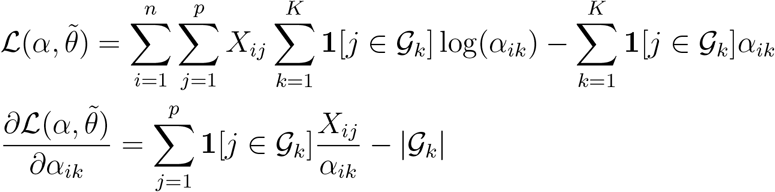

Now define 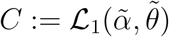 and define 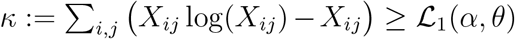 (with *x* log(*x*) defined as 0). *κ* is the unconstrained maximizer of the log likelihood function. For a given *ϵ* we can choose 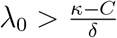 where *δ* is chosen such that 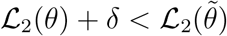 for all *θ* such that 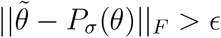 for all column permutations *P_σ_* (by the Lemma). For *λ*_0_, if we have:

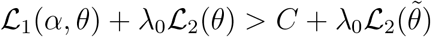

Then it follows that 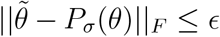 for some *σ*, since otherwise we would have 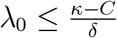. Further, for the chosen *λ*_0_ the argmax is nonempty as 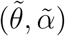 is a maximizer outside of a compact set (it is standard to show that the maximizing *α* is a continuous function of *θ* and the set 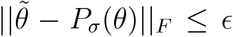 is compact so its image is compact). Since 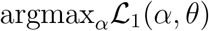 is a continuous function (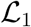 is smooth and under mild assumptions on *X* has a unique minimizer for each *θ*) it follows that for an *ϵ*_2_, *ϵ* can be chosen small enough so that 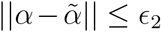.

##### Corollary 1

*For sufficiently large λ there is a global optimum of* 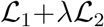 *which is arbitrarily close to the gene set averaging solution*.

This corollary explains why Spectra converges extremely quickly for large values of *λ*.

## References

1. Wolf, F.A., Angerer, P. & Theis, F.J. SCANPY: large-scale single-cell gene expression data analysis. Genome Biology 19, 15 (2018).

2. Bielecki, P., et al. Skin-resident innate lymphoid cells converge on a pathogenic effector state. Nature 592, 128–132 (2021).

3. Levitin, H.M., et al. De novo gene signature identification from single-cell RNA-seq with hierarchical Poisson factorization. Molecular Systems Biology 15, e8557 (2019).

4. Pelka, K., et al. Spatially organized multicellular immune hubs in human colorectal cancer. Cell.

5. Buettner, F., Pratanwanich, N., McCarthy, D.J., Marioni, J.C. & Stegle, O. f-scLVM: scalable and versatile factor analysis for single-cell RNA-seq. in Genome biology, Vol. 18 212 (2017).

6. Elyanow, R., Dumitrascu, B., Engelhardt, B.E. & Raphael, B.J. netNMF-sc: leveraging gene–gene interactions for imputation and dimensionality reduction in single-cell expression analysis. Genome research 30, 195–204 (2020).

7. Bassez, A., et al. A single-cell map of intratumoral changes during anti-PD1 treatment of patients with breast cancer. Nature Medicine (2021).

8. Stein-O’Brien, G.L., et al. Enter the Matrix: Factorization Uncovers Knowledge from Omics. Trends in Genetics 34, 790–805 (2018).

9. Boonyaratanakornkit, J.B., et al. Key gravity-sensitive signaling pathways drive T-cell activation. The FASEB Journal 19, 2020–2022 (2005).

10. Brauns, E., et al. Functional reprogramming of monocytes in patients with acute and convalescent severe COVID-19. JCI Insight 7(2022).

11. Satija, R., Farrell, J.A., Gennert, D., Schier, A.F. & Regev, A. Spatial reconstruction of single-cell gene expression data. Nature Biotechnology 33, 495–502 (2015).

12. Grasso, C.S., et al. Conserved Interferon-γ Signaling Drives Clinical Response to Immune Checkpoint Blockade Therapy in Melanoma. Cancer Cell 38, 500–515.e503 (2020).

13. Goswami, S., et al. Immune profiling of human tumors identifies CD73 as a combinatorial target in glioblastoma. Nature Medicine (2019).

14. Jorgovanovic, D., Song, M., Wang, L. & Zhang, Y. Roles of IFN-γ in tumor progression and regression: a review. Biomarker Research 8, 49 (2020).

15. van der Leun, A.M., Thommen, D.S. & Schumacher, T.N. CD8+ T cell states in human cancer: insights from single-cell analysis. Nature Reviews Cancer (2020).

16. Duhen, T., et al. Co-expression of CD39 and CD103 identifies tumor-reactive CD8 T cells in human solid tumors. Nature Communications 9, 2724 (2018).

17. Li, H., et al. Dysfunctional CD8 T Cells Form a Proliferative, Dynamically Regulated Compartment within Human Melanoma. Cell 176, 775–789 e718 (2019).

18. Thommen, D.S., et al. A transcriptionally and functionally distinct PD-1+ CD8+ T cell pool with predictive potential in non-small-cell lung cancer treated with PD-1 blockade. Nature Medicine 24, 994–1004 (2018).

19. Liu, B., Zhang, Y., Wang, D., Hu, X. & Zhang, Z. Single-cell meta-analyses reveal responses of tumor-reactive CXCL13+ T cells to immune-checkpoint blockade. Nature Cancer 3, 1123–1136 (2022).

20. Lee, Y.J., et al. CD39^+^ tissue-resident memory CD8^+^ T cells with a clonal overlap across compartments mediate antitumor immunity in breast cancer. Science Immunology 7, eabn8390 (2022).

21. Yost, K.E., et al. Clonal replacement of tumor-specific T cells following PD-1 blockade. Nature Medicine 25, 1251–1259 (2019).

22. Scott, A.C., et al. TOX is a critical regulator of tumour-specific T cell differentiation. Nature 571, 270–274 (2019).

23. Khan, O., et al. TOX transcriptionally and epigenetically programs CD8+ T cell exhaustion. Nature 571, 211–218 (2019).

24. Zhang, Y., et al. Single-cell analyses reveal key immune cell subsets associated with response to PD-L1 blockade in triple-negative breast cancer. Cancer Cell 39, 1578–1593.e1578 (2021).

25. Miller, B.C., et al. Subsets of exhausted CD8+ T cells differentially mediate tumor control and respond to checkpoint blockade. Nature Immunology 20, 326–336 (2019).

26. Siddiqui, I., et al. Intratumoral Tcf1+PD-1+CD8+ T Cells with Stem-like Properties Promote Tumor Control in Response to Vaccination and Checkpoint Blockade Immunotherapy. Immunity 50, 195–211.e110 (2019).

27. Schmid, P., et al. Pembrolizumab for Early Triple-Negative Breast Cancer. New England Journal of Medicine 382, 810–821 (2020).

28. Chowdhury, P.S., Chamoto, K., Kumar, A. & Honjo, T. PPAR-Induced Fatty Acid Oxidation in T Cells Increases the Number of Tumor-Reactive CD8+ T Cells and Facilitates Anti– PD-1 Therapy. Cancer Immunology Research 6, 1375–1387 (2018).

29. Caushi, J.X., et al. Transcriptional programs of neoantigen-specific TIL in anti-PD-1-treated lung cancers. Nature 596, 126–132 (2021).

30. Seo, H., et al. BATF and IRF4 cooperate to counter exhaustion in tumor-infiltrating CAR T cells. Nature Immunology 22, 983–995 (2021).

31. Chen, Y., et al. BATF regulates progenitor to cytolytic effector CD8+ T cell transition during chronic viral infection. Nature Immunology 22, 996–1007 (2021).

32. Boutet, M., et al. Memory CD8+ T cells mediate early pathogen-specific protection via localized delivery of chemokines and IFNγ to clusters of monocytes. Science advances 7, eabf9975 (2021).

33. Shanker, A., et al. CD8 T Cell Help for Innate Antitumor Immunity1. The Journal of Immunology 179, 6651–6662 (2007).

34. Franciszkiewicz, K., Boissonnas, A., Boutet, M., Combadière, C. & Mami-Chouaib, F. Role of Chemokines and Chemokine Receptors in Shaping the Effector Phase of the Antitumor Immune Response. Cancer Research 72, 6325–6332 (2012).

35. Yeong, J., et al. Intratumoral CD39+CD8+ T Cells Predict Response to Programmed Cell Death Protein-1 or Programmed Death Ligand-1 Blockade in Patients With NSCLC. Journal of Thoracic Oncology 16, 1349–1358 (2021).

36. Chow, A., et al. CD39 Identifies Tumor-Reactive CD8 T cells in Patients With Lung Cancer. bioRxiv, 2022.2001.2024.477554 (2022).

37. Leone, R.D. & Powell, J.D. Metabolism of immune cells in cancer. Nature Reviews Cancer 20, 516–531 (2020).

38. Artyomov, M.N. & Van den Bossche, J. Immunometabolism in the Single-Cell Era. Cell Metabolism 32, 710–725 (2020).

39. Costa da Silva, M., et al. Iron Induces Anti-tumor Activity in Tumor-Associated Macrophages. Frontiers in Immunology 8(2017).

40. Sun, J.-L., et al. Tumor cell-imposed iron restriction drives immunosuppressive polarization of tumor-associated macrophages. Journal of Translational Medicine 19, 347 (2021).

41. Lee, M.-S. & Bensinger, S.J. Reprogramming cholesterol metabolism in macrophages and its role in host defense against cholesterol-dependent cytolysins. Cell Mol Immunol 19, 327–336 (2022).

42. Behmoaras, J., et al. Macrophage Epoxygenase Determines a Profibrotic Transcriptome Signature. The Journal of Immunology 194, 4705–4716 (2015).

43. Vazquez Rodriguez, G., Abrahamsson, A., Turkina, M.V. & Dabrosin, C. Lysine in Combination With Estradiol Promote Dissemination of Estrogen Receptor Positive Breast Cancer via Upregulation of U2AF1 and RPN2 Proteins. Frontiers in Oncology 10(2020).

44. Misiewicz, M., et al. Identification of a Novel Endoplasmic Reticulum Stress Response Element Regulated by XBP1*. Journal of Biological Chemistry 288, 20378–20391 (2013).

45. Sasako, T., et al. Hepatic Sdf2l1 controls feeding-induced ER stress and regulates metabolism. Nature Communications 10, 947 (2019).

46. Sharma, R.B., Darko, C. & Alonso, L.C. Intersection of the ATF6 and XBP1 ER stress pathways in mouse islet cells. Journal of Biological Chemistry 295, 14164–14177 (2020).

47. Vekich, J.A., Belmont, P.J., Thuerauf, D.J. & Glembotski, C.C. Protein disulfide isomerase-associated 6 is an ATF6-inducible ER stress response protein that protects cardiac myocytes from ischemia/reperfusion-mediated cell death. Journal of Molecular and Cellular Cardiology 53, 259–267 (2012).

48. Lemarié, M., Chatonnet, F., Caron, G. & Fest, T. Early Emergence of Adaptive Mechanisms Sustaining Ig Production: Application to Antibody Therapy. Frontiers in Immunology 12(2021).

49. Ricci, D., Gidalevitz, T. & Argon, Y. The special unfolded protein response in plasma cells. Immunological Reviews 303, 35–51 (2021).

50. Dennler, P., Fischer, E. & Schibli, R. Antibody Conjugates: From Heterogeneous Populations to Defined Reagents. Antibodies 4, 197–224 (2015).

51. Wang, L., et al. Myeloid Cell–associated Resistance to PD-1/PD-L1 Blockade in Urothelial Cancer Revealed Through Bulk and Single-cell RNA Sequencing. Clinical Cancer Research (2021).

52. DeNardo, D.G. & Ruffell, B. Macrophages as regulators of tumour immunity and immunotherapy. Nature Reviews Immunology 19, 369–382 (2019).

53. Riihilä, P., et al. Complement Component C3 and Complement Factor B Promote Growth of Cutaneous Squamous Cell Carcinoma. The American Journal of Pathology 187, 1186–1197 (2017).

54. Haghverdi, L., Buettner, F. & Theis, F.J. Diffusion maps for high-dimensional single-cell analysis of differentiation data. Bioinformatics 31, 2989–2998 (2015).

55. Wolf, Y., et al. Autonomous TNF is critical for in vivo monocyte survival in steady state and inflammation. J. Exp. Med. 214, 905–917 (2017).

56. Lee, M.K.S., et al. Glycolysis Is Required for LPS-Induced Activation and Adhesion of Human CD14+CD16− Monocytes. Frontiers in Immunology 10(2019).

57. Lubbers, R., van Essen, M.F., van Kooten, C. & Trouw, L.A. Production of complement components by cells of the immune system. Clinical and experimental immunology 188, 183–194 (2017).

58. Dann, E., Henderson, N.C., Teichmann, S.A., Morgan, M.D. & Marioni, J.C. Milo: differential abundance testing on single-cell data using k-NN graphs. bioRxiv, 2020.2011.2023.393769 (2020).

59. Dykes, S.S., Fasanya, H.O. & Siemann, D.W. Cathepsin L secretion by host and neoplastic cells potentiates invasion. Oncotarget 10, 5560 (2019).

60. Rochefort, H. & Liaudet-Coopman, E. Cathepsin D in cancer metastasis: A protease and a ligand. APMIS 107, 86–95 (1999).

61. Vasiljeva, O., et al. Tumor Cell–Derived and Macrophage-Derived Cathepsin B Promotes Progression and Lung Metastasis of Mammary Cancer. Cancer Research 66, 5242–5250 (2006).

62. Lee, Y.S., et al. A small molecule targeting CHI3L1 inhibits lung metastasis by blocking IL-13Rα2-mediated JNK-AP-1 signals. Molecular Oncology 16, 508–526 (2022).

63. Huang, R., et al. Osteopontin Promotes Cell Migration and Invasion, and Inhibits Apoptosis and Autophagy in Colorectal Cancer by activating the p38 MAPK Signaling Pathway. Cellular Physiology and Biochemistry 41, 1851–1864 (2017).

64. He, Y., et al. Lipid Droplet-Related PLIN2 in CD68+ Tumor-Associated Macrophage of Oral Squamous Cell Carcinoma: Implications for Cancer Prognosis and Immunotherapy. Frontiers in Oncology 12(2022).

65. Baitsch, D., et al. Apolipoprotein E Induces Antiinflammatory Phenotype in Macrophages. Arteriosclerosis, Thrombosis, and Vascular Biology 31, 1160–1168 (2011).

66. Kemp, S.B., et al. Apolipoprotein E Promotes Immune Suppression in Pancreatic Cancer through NF-κB–Mediated Production of CXCL1. Cancer Research 81, 4305–4318 (2021).

67. Fuior, E.V. & Gafencu, A.V. Apolipoprotein C1: Its Pleiotropic Effects in Lipid Metabolism and Beyond. International Journal of Molecular Sciences 20, 5939 (2019).

68. Li, T., Chen, W. & Chiang, J.Y.L. PXR induces CYP27A1 and regulates cholesterol metabolism in the intestine. Journal of Lipid Research 48, 373–384 (2007).

69. Prabata, A., Ikeda, K., Rahardini, E.P., Hirata, K.-I. & Emoto, N. GPNMB plays a protective role against obesity-related metabolic disorders by reducing macrophage inflammatory capacity. Journal of Biological Chemistry 297, 101232 (2021).

70. Walle, T. Cytopus (2022).

## References

Edo M Airoldi, David Blei, Stephen Fienberg, and Eric Xing. Mixed membership stochastic blockmodels. Advances in neural information processing systems, 21, 2008.

Kevin Arvai. kneed, August 2020. URL https://doi.org/10.5281/zenodo.6944485. If you use this software, please cite it as below.

Dalia Barkley, Reuben Moncada, Maayan Pour, Deborah A Liberman, Ian Dryg, Gregor Werba, Wei Wang, Maayan Baron, Anjali Rao, Bo Xia, et al. Cancer cell states recur across tumor types and form specific interactions with the tumor microenvironment. Nature Genetics, 54(8):1192–1201, 2022.

Ayse Bassez, Hanne Vos, Laurien Van Dyck, Giuseppe Floris, Ingrid Arijs, Christine Desmedt, Bram Boeckx, Marlies Vanden Bempt, Ines Nevelsteen, Kathleen Lambein, et al. A single-cell map of intratumoral changes during anti-pd1 treatment of patients with breast cancer. Nature Medicine, 27(5):820–832, 2021.

Florian Buettner, Naruemon Pratanwanich, Davis J McCarthy, John C Marioni, and Oliver Stegle. f-sclvm: scalable and versatile factor analysis for single-cell rna-seq. Genome biology, 18(1):1–13, 2017.

David Croft, Gavin O’Kelly, Guanming Wu, Robin Haw, Marc Gillespie, Lisa Matthews, Michael Caudy, Phani Garapati, Gopal Gopinath, Bijay Jassal, Steven Jupe, Irina Kalatskaya, Shahana Mahajan, Bruce May, Nelson Ndegwa, Esther Schmidt, Veronica Shamovsky, Christina Yung, Ewan Birney, Henning Hermjakob, Peter D’Eustachio, and Lincoln Stein. Reactome: a database of reactions, pathways and biological processes. Nucleic Acids Research, 39(suppl 1):D691–D697, 11 2010. ISSN 0305-1048. doi: 10.1093/nar/gkq1018. URL https://doi.org/10.1093/nar/gkq1018.

Emma Dann, Neil C Henderson, Sarah A Teichmann, Michael D Morgan, and John C Marioni. Differential abundance testing on single-cell data using k-nearest neighbor graphs. Nature Biotechnology, 40(2):245–253, 2022.

Elizabeth A Eisenhauer, Patrick Therasse, Jan Bogaerts, Lawrence H Schwartz, Danielle Sargent, Robert Ford, Janet Dancey, S Arbuck, Steve Gwyther, Margaret Mooney, et al. New response evaluation criteria in solid tumours: revised recist guideline (version 1.1). European journal of cancer, 45(2):228–247, 2009.

Inbal Eizenberg-Magar, Jacob Rimer, Irina Zaretsky, David Lara-Astiaso, Shlomit Reich-Zeliger, and Nir Friedman. Diverse continuum of cd4¡sup¿+¡/sup¿ t-cell states is determined by hierarchical additive integration of cytokine signals. Proceedings of the National Academy of Sciences, 114(31):E6447–E6456, 2017. doi: 10.1073/pnas.1615590114. URL https://www.pnas.org/doi/abs/10.1073/pnas.1615590114.

Zhuoqing Fang, Xinyuan Liu, and Gary Peltz. GSEApy: a comprehensive package for performing gene set enrichment analysis in Python. Bioinformatics, 11 2022. ISSN 1367-4803. doi: 10.1093/bioinformatics/btac757. URL https://doi.org/10.1093/bioinformatics/btac757. btac757.

Adam Gayoso, Jonathan Shor, Ambrose J. Carr, Roshan Sharma, and Dana Pe’er. Doublet-detection (version v3.0). Zenodo, 2020.

Carmen Bravo González-Blas, Seppe De Winter, Gert Hulselmans, Nikolai Hecker, Irina Matetovici, Valerie Christiaens, Suresh Poovathingal, Jasper Wouters, Sara Aibar, and Stein Aerts. Scenic+: single-cell multiomic inference of enhancers and gene regulatory networks. bioRxiv, 2022.

Prem Gopalan, Jake M Hofman, and David M Blei. Scalable recommendation with hierarchical poisson factorization. In UAI, pages 326–335, 2015.

Laleh Haghverdi, Florian Buettner, and Fabian J Theis. Diffusion maps for high-dimensional single-cell analysis of differentiation data. Bioinformatics, 31(18):2989–2998, 2015.

Minoru Kanehisa, Miho Furumichi, Mao Tanabe, Yoko Sato, and Kanae Morishima. Kegg: new perspectives on genomes, pathways, diseases and drugs. Nucleic Acids Research, 45 (D1):D353–D361, 11 2016. ISSN 0305-1048. doi: 10.1093/nar/gkw1092. URL https://doi.org/10.1093/nar/gkw1092.

Zheng Tracy Ke, Yucong Ma, and Xihong Lin. Estimation of the number of spiked eigenvalues in a covariance matrix by bulk eigenvalue matching analysis. Journal of the American Statistical Association, pages 1–19, 2021.

DP Kingma and JL Ba. Adam: A method for stochastic optimization. 3rd int. conf. learn. represent. iclr 2015-conf. Track Proc., Dec, 2014.

Aaron T L Lun, Karsten Bach, and John C Marioni. Pooling across cells to normalize single-cell rna sequencing data with many zero counts. Genome biology, 17(1):1–14, 2016.

Daniel D Lee and H Sebastian Seung. Learning the parts of objects by non-negative matrix factorization. Nature, 401(6755):788–791, 1999.

Jacob H Levine, Erin F Simonds, Sean C Bendall, Kara L Davis, D Amir El-ad, Michelle D Tadmor, Oren Litvin, Harris G Fienberg, Astraea Jager, Eli R Zunder, et al. Data-driven phenotypic dissection of aml reveals progenitor-like cells that correlate with prognosis. Cell, 162(1):184–197, 2015.

Hanna Mendes Levitin, Jinzhou Yuan, Yim Ling Cheng, Francisco JR Ruiz, Erin C Bush, Jeffrey N Bruce, Peter Canoll, Antonio Iavarone, Anna Lasorella, David M Blei, et al. De novo gene signature identification from single-cell rna-seq with hierarchical poisson factorization. Molecular systems biology, 15(2):e8557, 2019.

Arthur Liberzon, Aravind Subramanian, Reid Pinchback, Helga Thorvaldsdóttir, Pablo Tamayo, and Jill P. Mesirov. Molecular signatures database (msigdb) 3.0. Bioinformatics, 27(12):1739–1740, 05 2011. ISSN 1367-4803. doi: 10.1093/bioinformatics/btr260. URL https://doi.org/10.1093/bioinformatics/btr260.

Arthur Liberzon, Chet Birger, Helga Thorvaldsdóttir, Mahmoud Ghandi, Jill P Mesirov, and Pablo Tamayo. The molecular signatures database hallmark gene set collection. Cell systems, 1(6):417–425, 2015.

Jun S Liu and Ying Nian Wu. Parameter expansion for data augmentation. Journal of the American Statistical Association, 94(448):1264–1274, 1999.

Emma Pierson and Christopher Yau. Zifa: Dimensionality reduction for zero-inflated single-cell gene expression analysis. Genome biology, 16(1):1–10, 2015.

Ruslan Salakhutdinov, Sam T Roweis, and Zoubin Ghahramani. Optimization with em and expectation-conjugate-gradient. In Proceedings of the 20th International Conference on Machine Learning (ICML-03), pages 672–679, 2003.

Manu Setty, Vaidotas Kiseliovas, Jacob Levine, Adam Gayoso, Linas Mazutis, and Dana Pe’Er. Characterization of cell fate probabilities in single-cell data with palantir. Nature biotechnology, 37(4):451–460, 2019.

Keith Stevens, Philip Kegelmeyer, David Andrzejewski, and David Buttler. Exploring topic coherence over many models and many topics. In Proceedings of the 2012 joint conference on empirical methods in natural language processing and computational natural language learning, pages 952–961, 2012.

The Gene Ontology Consortium. The gene ontology resource: 20 years and still going strong. Nucleic Acids Research, 47(D1):D330–D338, 11 2018. ISSN 0305-1048. doi: 10.1093/nar/gky1055. URL https://doi.org/10.1093/nar/gky1055.

Robert Tibshirani. Regression shrinkage and selection via the lasso. Journal of the Royal Statistical Society: Series B (Methodological), 58(1):267–288, 1996.

David Van Dijk, Roshan Sharma, Juozas Nainys, Kristina Yim, Pooja Kathail, Ambrose J Carr, Cassandra Burdziak, Kevin R Moon, Christine L Chaffer, Diwakar Pattabiraman, et al. Recovering gene interactions from single-cell data using data diffusion. Cell, 174(3): 716–729, 2018.

Thomas Walle. wallet-maker/cytopus: Cytopus v1.21, November 2022. URL https://doi.org/10.5281/zenodo.7306238.

F Alexander Wolf, Philipp Angerer, and Fabian J Theis. Scanpy: large-scale single-cell gene expression data analysis. Genome biology, 19(1):1–5, 2018.

Yuanyuan Zhang, Hongyan Chen, Hongnan Mo, Xueda Hu, Ranran Gao, Yahui Zhao, Baolin Liu, Lijuan Niu, Xiaoying Sun, Xiao Yu, et al. Single-cell analyses reveal key immune cell subsets associated with response to pd-l1 blockade in triple-negative breast cancer. Cancer Cell, 39(12):1578–1593, 2021.

