## Supplementary Figures for "Supervised discovery of interpretable gene programs from single-cell data"

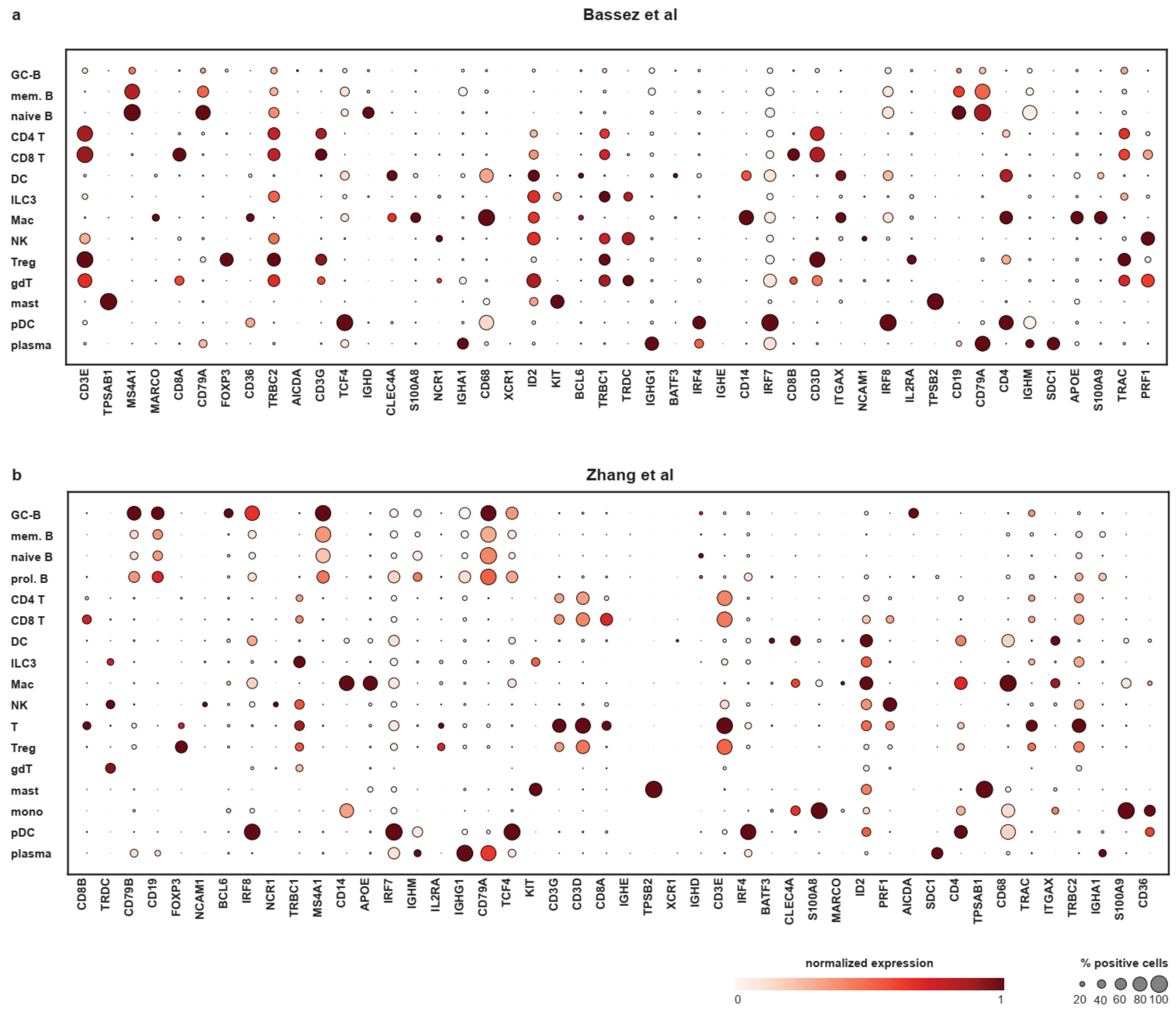


**Supplementary Fig. 1 | Cell type annotations in breast cancer datasets.** [Related to **Fig. 2**.]

Marker gene expression for each of 14 broad cell-type annotations in the Bassez[7] (top, n=97,863 cells) and Zhang[24] (bottom, n=150,985 cells) datasets. Each gene is normalized to its maximal expression (color bar, normalized expression bar), and the percentage of cells that express the gene within a cluster is indicated (dot size, % positive cells). ILC3: innate lymphoid cell type 3, T: T cell, gdT: gamma-delta T cell, pDC: plasmacytoid dendritic cell, NK: Natural Killer cell, B: B cell, Mac: macrophage, mast: mast cell, mem.: memory, mono: monocyte, prol.: proliferating, Treg: regulatory T cell, DC: dendritic cell, GC-B: germinal center B cell, plasma: plasma cell.

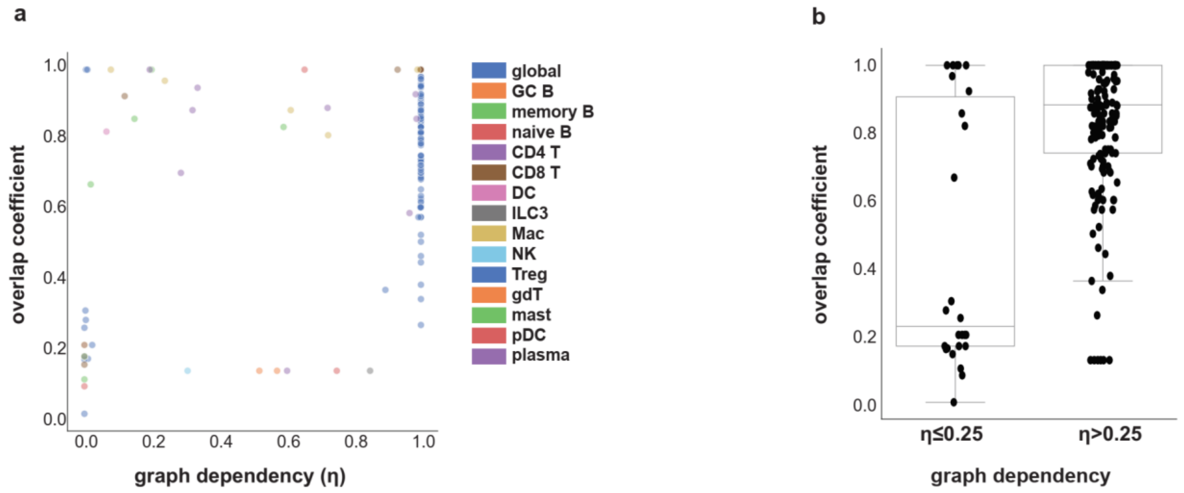


**Supplementary Fig. 2 | Dependence of Spectra factors on input gene sets.** [Related to **Fig. 2**.]

**a**, Maximum overlap coefficient for each factor (n=197), denoting overlap with input gene sets, plotted against Spectra's graph dependency parameter (eta, η); each factor is colored by cell type. Most factors are highly dependent on the gene-gene graph and exhibit correspondingly high overlap coefficients. **b**, Maximum overlap coefficient of each factors (n=197) with input gene sets and a high (η ≥ 0.25) or low (η < 0.25) dependence parameter. Boxes and line represent interquartile range (IQR) and median, respectively; whiskers represent ±1.5 x IQR. ILC3: innate lymphoid cell type 3, T: T cell, gdT: gamma-delta T cell, pDC: plasmacytoid dendritic cell, NK: Natural Killer cell, B: B cell, mast: mast cell, Treg: regulatory T cell, DC: dendritic cell, GC B: germinal center B cell, plasma: plasma cell.

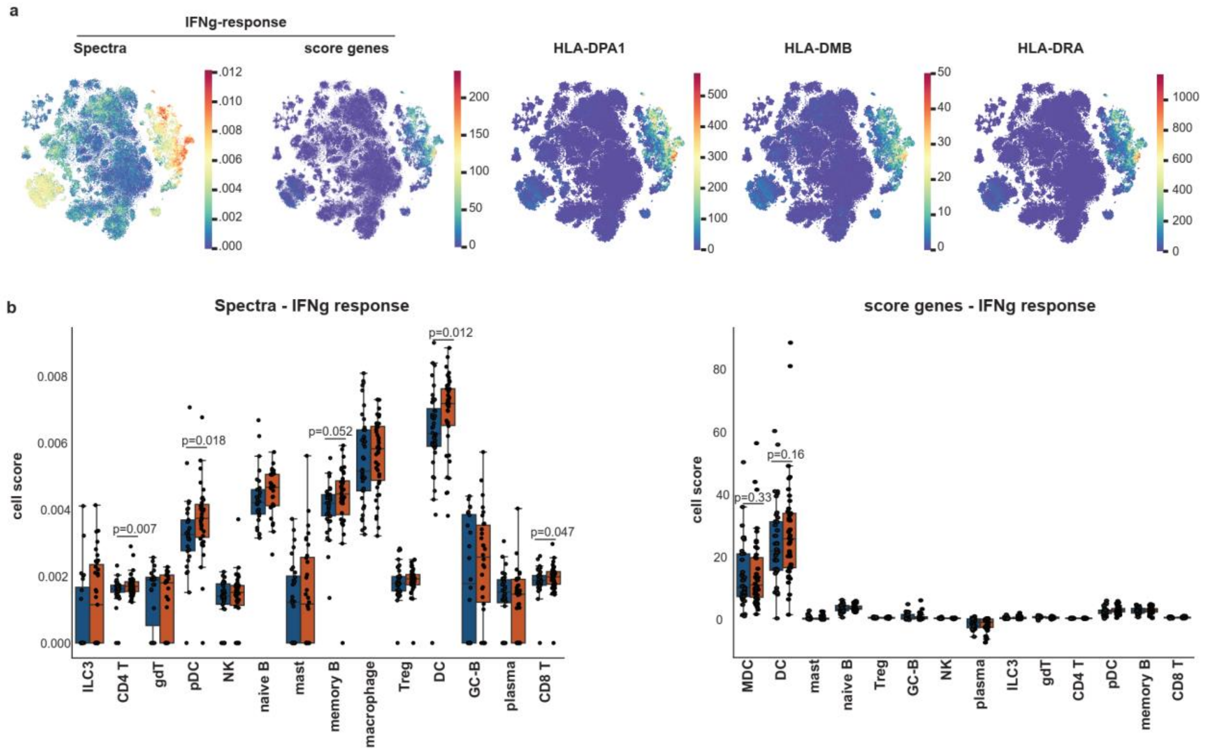


**Supplementary Fig. 3 | Spectra discerns the effects of immune checkpoint therapy on interferon signaling.** [Related to **Fig. 2**.]

**a**, t-SNE projections of the Bassez dataset[7] tumor-infiltrating leukocytes (n=97,863), colored by cell scores for Spectra or scanpy.score_genes interferon gamma (IFNg) response, or by expression of selected human leukocyte antigen (HLA) class II genes. HLA expression is scran-normalized and not imputed. **b**, Mean cell score of the positive fraction (cell score > 0.01) per sample and cell type before (blue, n = 40) and after (red, n = 40) anti-PD-1 immune checkpoint blockade in breast tumor infiltrating leukocyte data from the Bassez dataset. P values (two-sided) calculated using Wilcoxon matched-pairs signed rank tests. Test statistics, left panel: 282 (Treg), 184 (memory B), 75 (mast), 82 (pDC), 237 (CD4 T), 222 (DC), 129 (naive B), 289 (NK), 293 (CD8 T), 76 (gdT), 59 (GC-B), 339 (macrophage), 24 (ILC3), 220 (plasma). Test statistics, right panel: 340 (Treg), 241 (memory B), 168 (mast), 131 (pDC), 362 (CD4 T), 305 (DC), 121 (naive B), 349 (NK), 427 (CD8 T), 71 (gdT), 104 (GC-B), 356 (macrophage), 125 (ILC3), 262 (plasma). Cohen's d, left panel: 0.049 (NK), 0.320 (CD4 T), 0.228 (GC B), 0.105 (CD8 T), 0.075 (naïve B), 0.064 (Treg), 0.285 (DC), 0.142 (mast), 0.093 (Mac), 0.131 (plasma), 0.171 (ILC3), 0.341 (pDC), 0.067 (gdT), 0.197 (memory B). Cohen's d, right panel: 0.168 (NK), 0.142 (CD4 T), 0.039 (GC-B), 0.040 (CD8 T), 0.187 (naïve B), 0.008 (Treg), 0.166 (DC), 0.003 (mast), 0.064 (Mac), 0.139 (plasma), 0.008 (ILC3), 0.202 (pDC), 0.170 (gdT), 0.146 (memory B). Boxes and line represent interquartile range (IQR) and median, respectively; whiskers represent ±1.5 x IQR. ILC3: innate lymphoid cell type 3, T: T cell, gdT: gamma-delta T cell, pDC: plasmacytoid dendritic cell, Mac: macrophage, NK: Natural Killer cell, B: B cell, mast: mast cell, Treg: regulatory T cell, DC: dendritic cell, GC-B: germinal center B cell, plasma: plasma cell.

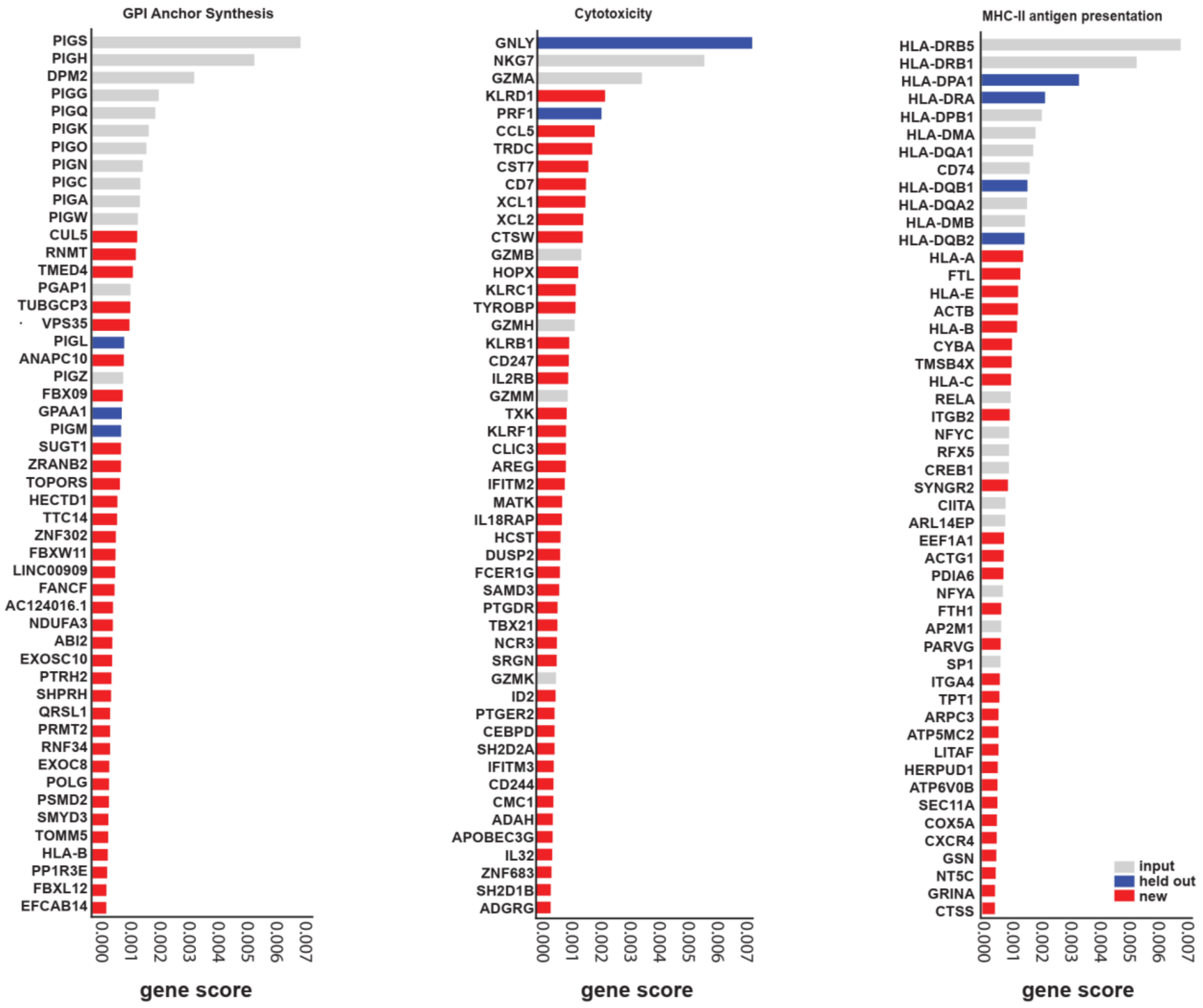


**Supplementary Fig. 4. | Spectra recovers genes involved in diverse cellular processes.** [Related to **Fig. 3.**]

Top 50 marker genes for three cellular process factors identified by Spectra in the Bassez dataset[7], after holding out a randomly selected 40% subset from each corresponding input gene set. Marker genes are ranked by Spectra factor score. New genes (absent from the original input gene set) and held-out genes recovered by Spectra are highlighted in red and blue, respectively.

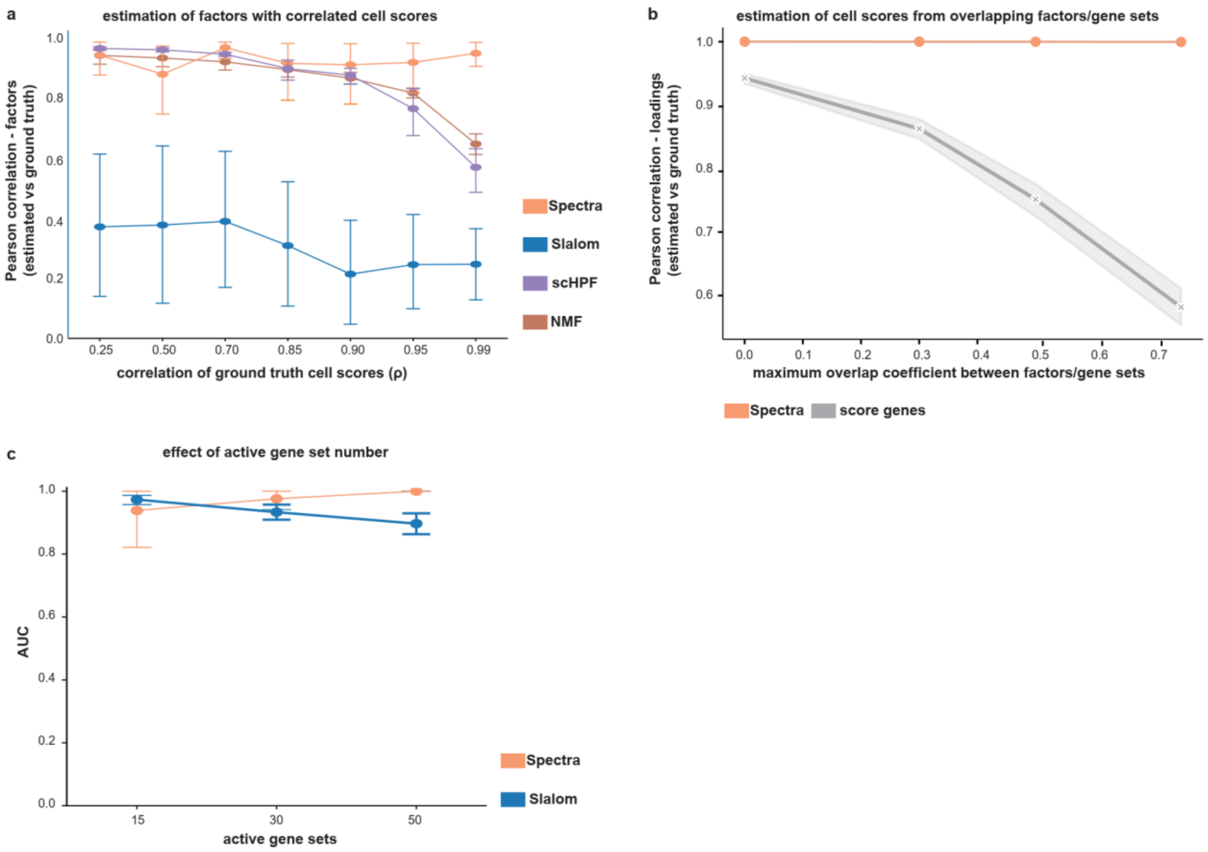


**Supplementary Fig. 5 | Spectra accurately quantifies the activity of inferred factors in simulated data.** [Related to **Fig. 3**.]

**a**, Spectra retrieves highly correlated simulated gene programs more accurately than other methods. Synthetic data was generated by sampling random ground truth loadings and factors from log-normal and half-Cauchy distributions, respectively, and introducing correlations in two factors via off-diagonal entries in the log-normal covariance matrix. After mutiplying loadings and factor matrices and introducing noise, models were fit to the data and output factors were correlated with ground truth factors. **b**, Correlation between ground truth and inferred loadings (cell scores) for score_genes and Spectra, as a function of gene set overlap in simulated data. Data consisted of overlapping synthetic gene sets and a random cell loading vector representing the expression of each gene set in a cell (Methods). The sum of gene sets weighted by the individual cell loadings for each cell was used to represent a mean for sampling Poisson gene expression counts. **c**, Slalom and Spectra are robust to highly overlapping gene sets, but Slalom suffers worse performance when the number of active gene sets is large. Gene expression data was simulated from a factor analysis model in which only a subset of gene sets are active in the data, similar to **a** and the original Slalom publication[5]. y axis, area under the ROC curve. Intervals and lines represent 95% CI and mean, respectively, across n = 10 (**a**), n = 3 (**b**), and n = 5 (**c**) independent simulations.

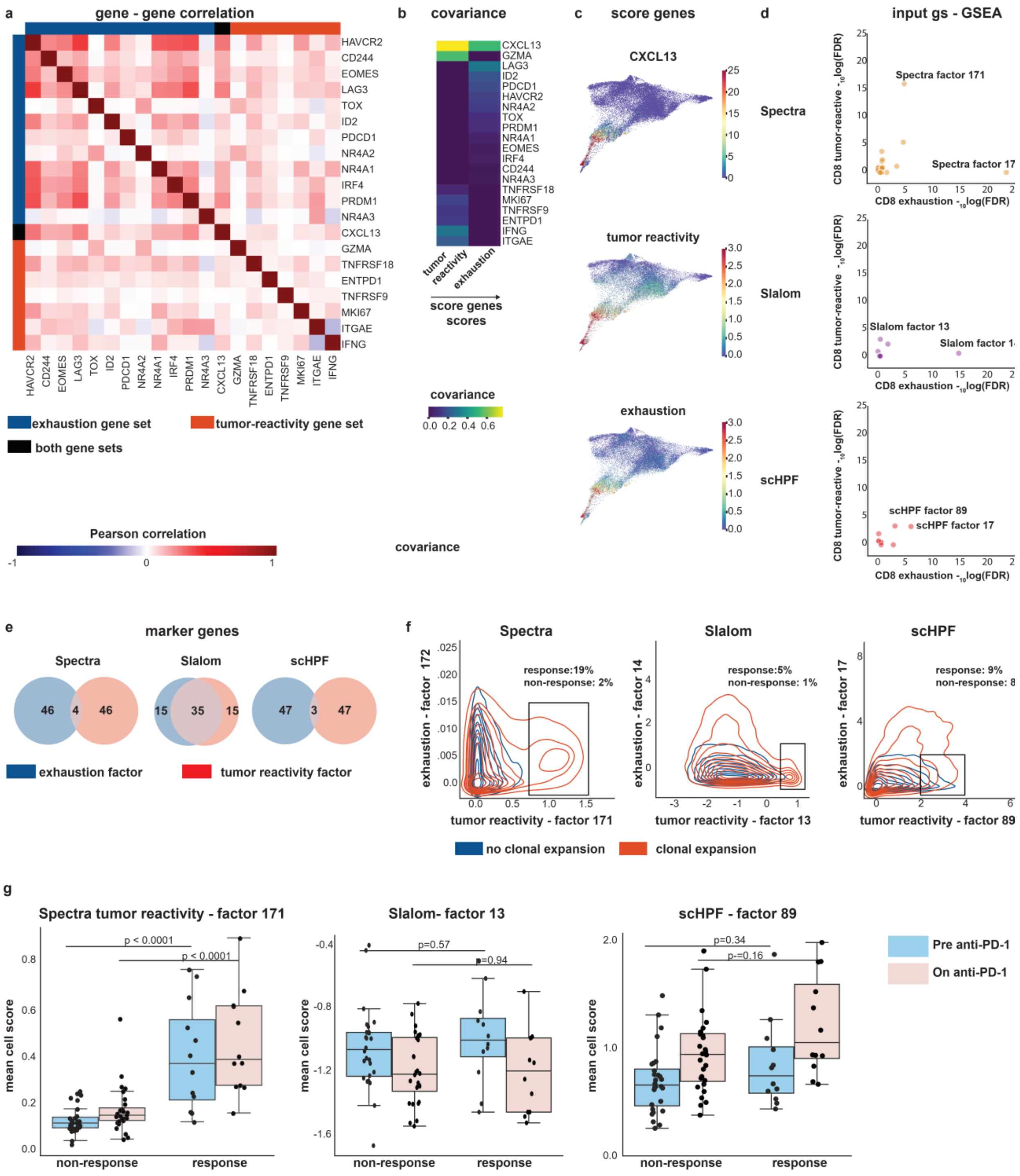


**Supplementary Fig. 6 | Existing methods fail to separate highly correlated features in CD8$^+$ T cell data.** [Related to **Fig. 4**.]

Analysis of breast cancer infiltrating leukocytes from the Bassez scRNA-seq dataset (n = 42 patients)[7]. **a**, Pearson gene-gene correlations reveal that CD8$^+$ T cell exhaustion and tumor reactivity gene sets are highly correlated (n=43,066 cells). **b,** Covariance between individual genes and their respective average gene set scores (n=43,066 cells). *CXCL13* is most highly correlated with both tumor reactivity and exhaustion. **c**, *CXCL13* expression maps to similar cells as both gene set scores in FDL of CD8$^+$ T cells (n=43,066). **d**, Significance (–10log(FDR)) of CD8$^+$ T cell exhaustion or tumor reactivity factors by gene set enrichment analysis (ssGSEA, Spectra: n=159, Slalom: n=20, scHPF: n=100 factors). **e**, Overlap between the top 50 marker genes of CD8+ T cell exhaustion and tumor reactivity factors. **f**, Contour plots indicating density of Spectra, Slalom or scHPF loading scores for CD8+ T cell exhaustion and tumor reactivity factors grouped by clonal T cell expansion status (n=43,066 cells). **g**, Spectra, Slalom and scHPF average cell scores per sample of the positive fraction (> 0.01). Patient samples are grouped by clonal T cell expansion status and time (baseline or on-treatment). Boxes and line represent interquartile range (IQR) and median, respectively; whiskers represent ±1.5 x IQR. P values (two-sided) were calculated using Mann-Whitney-U tests (n=40 pre anti-PD-1, n=40 on anti-PD-1 samples): Spectra: $p=3.835 \times 10^{-5}$ statistic: 308; on anti-PD-1: $p = 2.001 \times 10^{-5}$, statistic = 313; Slalom pre anti-PD-1: p=0.57 statistic: 198 Cohen's d: 0.249 ; Slalom on anti-PD-1: p=0.94 statistic: 155 Cohen's d: 0.059; scHPF pre anti-PD-1: p=0.34 statistic: 216 Cohen's d: 0.337; scHPF on anti-PD-1: p=0.16 statistic: 201 Cohen's d: 0.592.

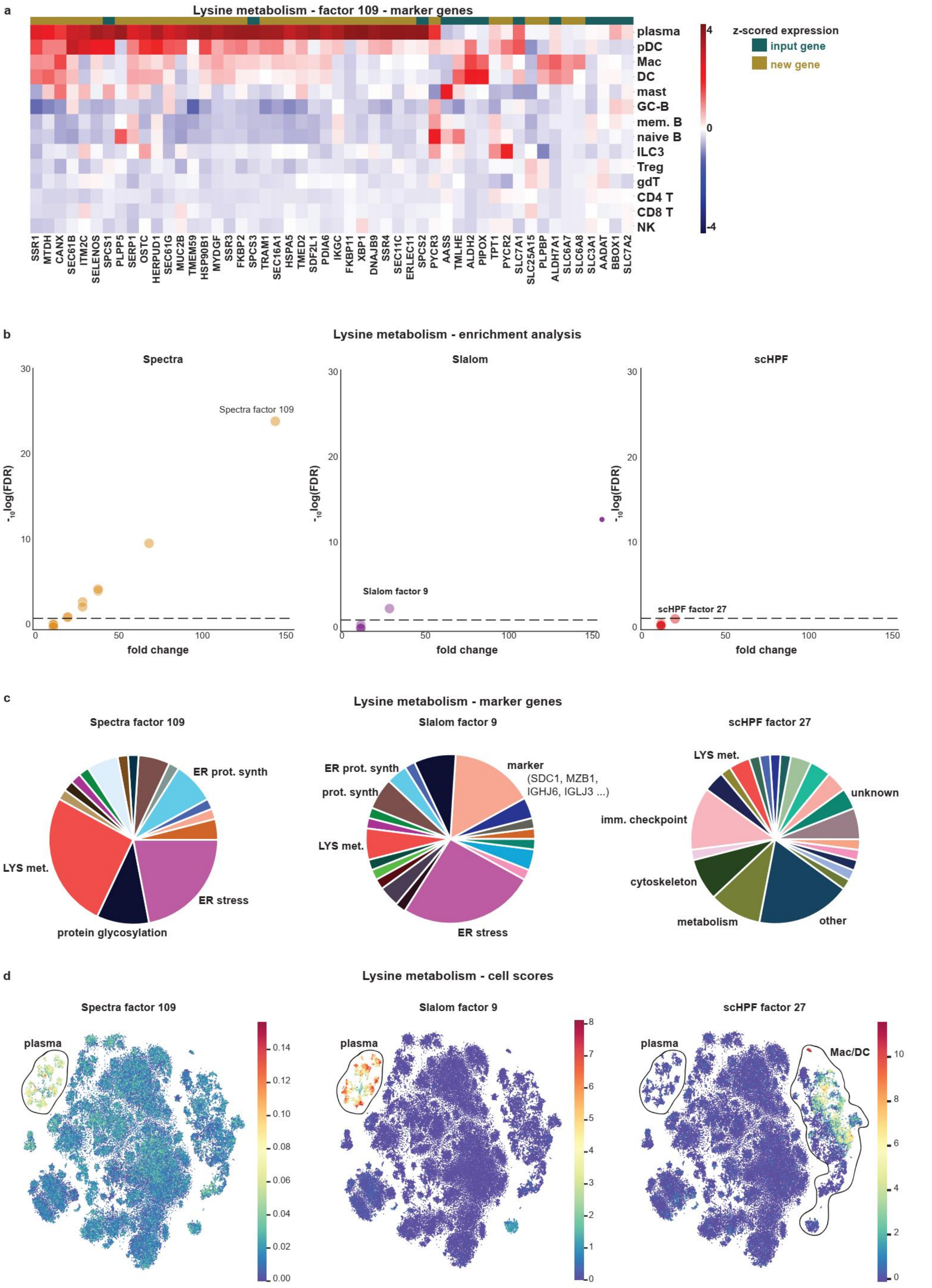


**Supplementary Fig. 7 | Spectra finds more specific and biologically coherent lysine metabolism factors than other methods.** [Related to **Fig. 5**.]

**a**, Z-scored average cellular expression (per cell type) of lysine factor genes (n=97,863 leukocytes). **b**, Significance (-10log(FDR)) and fold enrichment (odds ratio) of the lysine metabolism input gene set, among genes with highest gene scores (top 50 marker genes), as calculated by gene set enrichment analysis (Spectra: n=152 factors; Slalom: n=20 factors;

scHPF: n=100 factors). **c**, Functional categories of the top 50 marker genes of lysine metabolism factors identified by different factorization methods. **d**, t-SNE embeddings colored by cell scores for the top-performing lysine metabolism factors (labeled in **b**) from different factorization methods (n=97,863 leukocytes). Plasma, macrophage (Mac) and dendritic cell (DC) populations are outlined. ILC3: innate lymphoid cell type 3, T: T cell, gdT: gamma-delta T cell, pDC: plasmacytoid dendritic cell, Mac: macrophage, mono: monocyte, NK: Natural Killer cell, B: B cell, mast: mast cell, Treg: regulatory T cell, DC: dendritic cell, GC B: germinal center B cell, plasma: plasma cell.

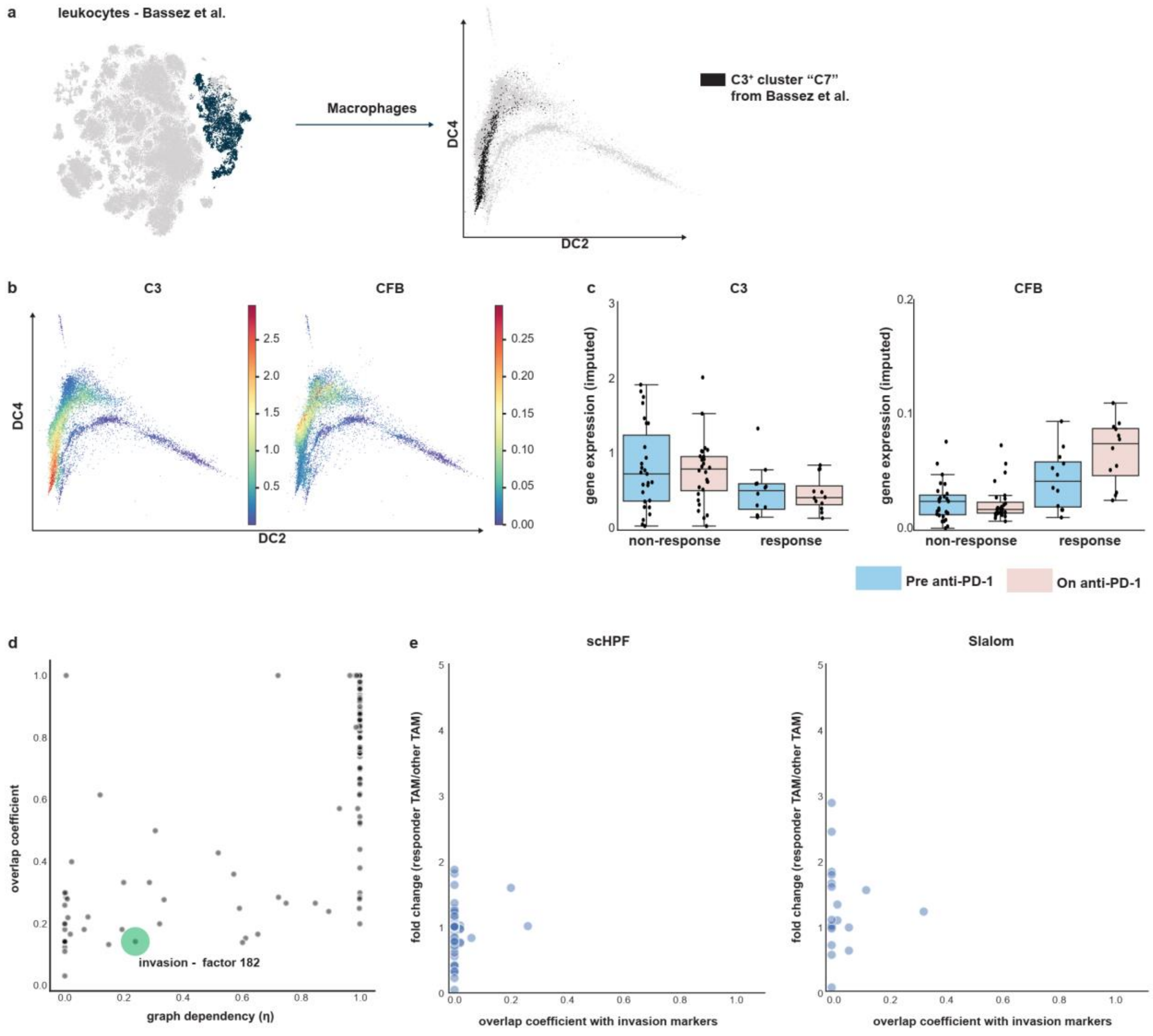


**Supplementary Fig. 8 | Tumor-infiltrating macrophage cell states exist along continua that change under therapy.** [Related to **Fig. 6**.]

**a**, t-SNE embedding of all leukocytes (left) and distribution of macrophages/monocytes (n=12,132) along diffusion components 2 and 4 (DC2 and DC4, right), highlighting cells of C3-positive macrophage cluster C7 from the Bassez dataset[7]. **b**, Macrophages (n=12,132) along DC2 and DC4, colored by MAGIC-imputed (t = 3) complement gene expression. **c**, mean MAGIC-imputed (t = 3) complement gene expression per sample in macrophages from responsive (E) or non-responsive (NE) patients sampled before (pre, n = 40) or during (on, n = 40) anti-PD-1 therapy. Boxes and line represent interquartile range (IQR) and median, respectively; whiskers represent ±1.5 x IQR. **d**, Overlap coefficient and graph dependency parameter for Spectra factors (n=197). **e**, scHPF (n=100) and Slalom (n=20) factors do not resemble the Spectra invasion factor (factor 182). Each factor is plotted by the fold change of its cell score in macrophage neighborhoods enriched in non-responders under therapy compared to its cell score in all remaining macrophages (y axis), and by the coefficient of overlap between the top 50 marker genes of each factor and of the Spectra invasion factor (x-axis) (analogous to **Fig. 6c,d**).
